# Micro-scale spatial metagenomics: revealing high-resolution spatial biogeography of gut microbiomes

**DOI:** 10.1101/2025.09.30.679663

**Authors:** Carlotta Pietroni, Bryan Wang, Amalia Bogri, Jorge Langa, Iñaki Odriozola, Zoé Horisberger, Marta Contreras-Serrano, Jonas Greve Lauritsen, Nanna Gaun, Amalia Toffano, Anders Miki Bojesen, Ida Thøfner, Victoria Drauch, Søren Johannes Sørensen, Urvish Trivedi, Antton Alberdi

## Abstract

Spatial organisation is a fundamental yet poorly resolved aspect of gut microbial ecology. Conventional shotgun metagenomics provides rich functional information but relies on homogenised, macro-scale samples that obscure the micron-scale distributions critical for understanding microbial community dynamics. Here, we introduce Micro-Scale Spatial Metagenomics (MSSM), a new methodological framework that couples laser micro-dissection of tissue sections, ultra-low-input library preparation, and genome-resolved bioinformatics to reconstruct microbial communities from intestinal microsamples measuring as little as ∼500 µm² (≈100 bacterial cells). We describe a fully optimised laboratory and computational pipeline that enables quantitative, strain-resolved, and functionally informed spatial profiling directly from intact gut tissue. Using chicken intestinal samples, we validated MSSM through combinatorial single-cell fluorescence *in situ* hybridisation (FISH) imaging and comparisons with macro-scale metagenomics, demonstrating its robustness and accuracy. MSSM captured fine-scale heterogeneity in taxonomic and functional composition across intestinal cryosections, hinting at spatially structured assemblages and segregation of metabolic capacities. Strain-level analyses uncovered coexisting *Lawsonibacter* lineages exhibiting distinct spatial distributions and host-specific occurrence patterns, while SNP-level microdiversity analyses showed that genetically coherent clonal populations cluster at spatial scales below ∼200 µm. By enabling shotgun metagenomics at micron resolution, MSSM closes a longstanding methodological gap and provides a scalable platform for studying microbial ecosystems *in situ*. This approach unlocks a previously inaccessible view of microbial biogeography, offering new opportunities to investigate host–microbe and microbe-microbe interactions, and the spatial principles governing gut ecosystems.

**Significance statement:** Understanding how microbial communities are organised in space is essential to explaining their ecological and functional roles, yet microbiome research still relies overwhelmingly on bulk, spatially averaged measurements. We introduce micro-scale spatial metagenomics (MSSM), the first method that brings shotgun metagenomics to the microscale, enabling direct measurement of functional and taxonomic variation across regions containing as few as ∼100 cells. Unlike existing spatial approaches, MSSM reconstructs complete genomes and resolves strain-level diversity within intact tissue, allowing researchers to map metabolic potential, microdiversity, and community structure *in situ*. By coupling high-resolution sequencing with spatial context, MSSM reveals a previously inaccessible layer of microbial organisation, transforming how host-associated ecosystems can be studied.

## Main

Microbiomes are fundamental to the functioning of most natural ecosystems (1–3), owing to the coordinated activity of the diverse microbial communities that comprise them (4). While metagenomics has significantly advanced our ability to characterise the functional potential of microorganisms (5, 6), understanding the interactions that underpin these functions remains a major challenge. This limitation constrains our capacity to predict the assembly, behaviour, and evolution of complex microbial consortia (7). Without accurate models of community functioning, our capacity to forecast the spread of infectious diseases, design targeted probiotics, or engineer optimal synthetic communities remains restricted.

One key barrier to deeper insights into microbiome functionality is our limited understanding of their spatial organisation (8). Although the importance of spatial structure in shaping biological interactions has long been recognised (9, 10), micro-scale research has been severely impeded by technical limitations in generating spatial data. Most existing knowledge about spatial architecture of microbiomes stems from fluorescence-based imaging techniques (11–13), which offer high-resolution insights through labelled microscopy. Advances in optics, molecular biology, and image analysis have enabled the simultaneous visualisation of dozens of microbial taxa (14, 15). However, the complexity of most natural microbiomes often surpasses what can be captured through fluorescence *in situ* hybridisation (FISH). To address this, various spatially resolved amplicon sequencing techniques have emerged as potential alternatives (16–18).

Despite these advancements, both imaging and amplicon-based sequencing still fall short in capturing the functional diversity of microbiomes, which frequently comprise hundreds of coexisting strains. Notably, strains with identical 16S rRNA gene sequences, which are primarily targeted by both aforementioned methods, can differ significantly in their functional capabilities and response to environmental stimuli that influence population- and community-level dynamics. Shotgun metagenomics offers a path toward strain-level functional resolution by enabling comprehensive genomic profiling (6, 19, 20). However, conventional implementations of this approach require relatively large amounts of template DNA derived from homogenised samples, which lack the spatial resolution needed to discern microbial interactions (21, 22).

To address this knowledge gap, we introduce micro-scale spatial metagenomics (MSSM) technology. MSSM integrates laser micro-dissection (LMD) with ultra-low-biomass shotgun metagenomic sequencing to enable the reconstruction of spatial patterns of microbial functional traits at micron-scale resolution (Fig. 1). Here, we present both the laboratory and computational workflows for MSSM, and showcase its application in the context of intestinal microbiomes of chickens, while validating and complementing the results with combinatorial FISH (Fig. 1). We show that MSSM can accurately recover spatially referenced microbial communities comprising down to ∼100 cells, unveiling fine-scale spatial heterogeneity within microbiomes.

**Fig. 1:**
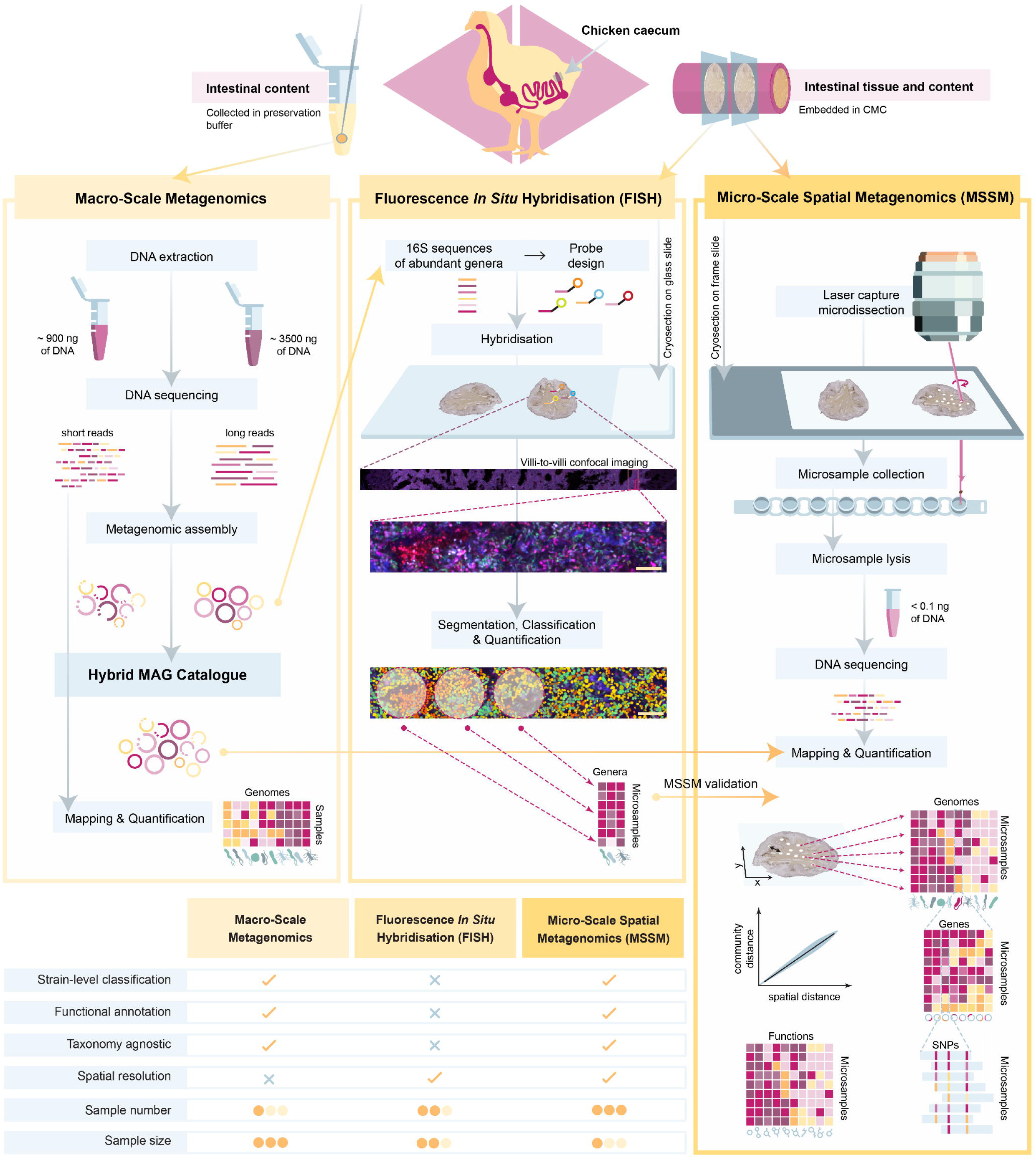
MSSM: micro-scale spatial metagenomics. Overview of the MSSM data generation and analysis workflow, compared with macro-scale metagenomics and fluorescence *in situ* hybridisation (FISH) for bacterial community characterisation. Arrows indicate how the three methods were applied to improve, complement, and validate one another. The accompanying table summarises the differences across key features, from sample processing to the resolution and type of microbial information each method can achieve.

## Results

### Microbial genomic reference data for micro-scale spatial metagenomics (MSSM)

Quantifying microbial communities using shotgun metagenomics depends on a comprehensive reference catalogue of metagenome-assembled genomes (MAGs) to accurately map sequencing reads (19). To test whether micro-scale spatial metagenomics (MSSM) can generate a catalogue on its own despite the low biomass of microsamples, or if it needs a reference catalogue from high-biomass macro-scale samples, we compared catalogue generation at both scales and evaluated their strengths and limitations.

#### Micro-scale MAG catalogue captures microbiome complexity

The MAG catalogue reconstructed from microsamples comprised 122 genomes, with a median completeness of 96.94% (IQR = 91.49-99.49%) and contamination rate of 0.58% (IQR = 0.21-1.83%). These values were comparable to those obtained using a hybrid long-and short-read sequencing strategy applied to macro-scale samples (completeness: median = 99.98%, IQR = 94.60-100%; contamination: median = 0.23%, IQR = 0-1.71%) (Fig. 2a), demonstrating the feasibility of generating a comprehensive reference catalogue that captures functional microbiome attributes directly from microsamples. The micro-scale approach effectively captured the complexity of the microbiome, with a comparable number of reads mapping against both catalogues (S. Note 1). The results also revealed similar alpha diversity and community composition between the two catalogues (S. Note 1), demonstrating that MSSM can serve as a standalone method for species-level microbial quantification without relying on macro-scale inputs. However, the macro-scale approach yielded 98 circularised genomes (Fig. 2b), while the micro-scale approach, lacking long-read sequences, yielded none (Supplementary Fig. 1). The circularised genomes enabled more detailed and complementary micro-scale analyses, including SNP-based investigations detailed in the method implementation section.

**Fig. 2:**
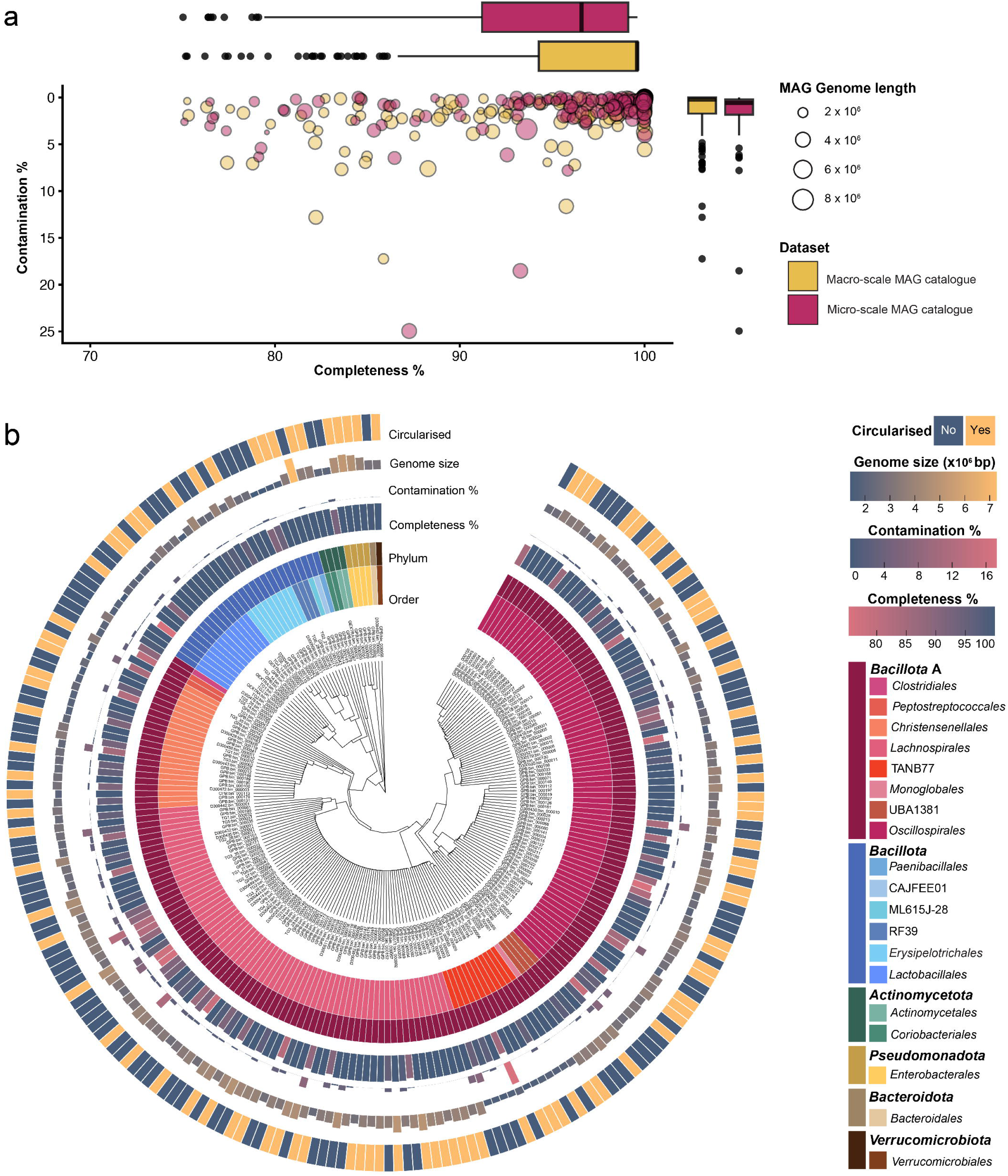
Comparative assessment of metagenome-assembled genome (MAG) catalogues derived from conventional macro-scale metagenomics and micro-scale spatial metagenomics (MSSM). **a,** Comparison of completeness and contamination levels across MAGs obtained from the two datasets. **b,** Summary of key features of the macro-scale MAG catalogue used as primary reference, including metrics on completeness, contamination, taxonomic diversity, genome size, and circularisation status.

#### Macro-scale MAG catalogue enables the design of unique fluorescence *in situ* hybridisation (FISH) targets

Another advantage of generating the macro-scale MAG catalogue based on long-read sequencing was the recovery of full-length 16S rRNA gene sequences, which were essential for designing accurate fluorescence *in situ* hybridisation (FISH) probes to validate MSSM (Supplementary Fig. 2). Our initial probe design relied on publicly available 16S rRNA gene sequences (S. Note 2). However, reference sequences were available for only 10 of the 15 most abundant genera detected by MSSM, out of which hybridisation signals were detected for only 5 genera due to sequence mismatches (S. Note 2). In contrast, by utilising full-length 16S rRNA gene sequences recovered from the macro-scale MAG catalogue, we were able to design genus-specific probes without cross-hybridisation for 9 of the 15 most abundant taxa observed in MSSM data (Supplementary Fig. 2; S. Note 2).

### MSSM method development

The development of MSSM involved optimising each step of the workflow by systematically evaluating histological, biochemical, and experimental design variables.

#### Segmentation, embedding and laser micro-dissection (LMD)

The protocol (Fig. 1) starts by embedding snap-frozen intestinal sections and cryosectioning them into 10 μm-thick tissue sections. To ensure compatibility with mass spectrometry-based methods such as metabolomics, we tested the spectrometry-compatible carboxymethylcellulose (CMC) along with the commonly used optimal cutting temperature (OCT) compound, which is known to interfere with mass spectrometry (23, 24). Both embedding materials were used on the same intestinal section, with no differences observed in cryosection quality and downstream MSSM processing (Supplementary Fig. 3a).

While fixing and staining procedures are commonly used to improve tissue preservation and imaging contrast (25, 26), washing steps led to significant loss of the more soluble intestinal content where the microbial communities of interest were located (Supplementary Fig. 3b). To maximise intestinal content retention, laser micro-dissection (LMD) procedures were developed with unfixed and unstained tissue sections in brightfield mode, which enabled to visibly identify regions with digesta and distinguish them from empty membrane regions caused by sectioning artifacts (Supplementary Fig. 3b). The LMD settings were optimised to ensure complete cuts through the mounted cryosection and slide membrane (Supplementary Table 1), as repeated laser exposure can lead to sample deterioration.

A major challenge in LMD was the tendency of microdissections to adhere to the underside of the membrane instead of falling into the collector reaction well. To improve sampling success, 8-strip caps were used as collectors instead of 96-well plates, enabling visual confirmation of collected microsamples and allowing for resampling from an adjacent area when necessary (Supplementary Fig. 4a). As a result of this, 97.8 % of the microsamples were successfully collected in a maximum of three attempts (Supplementary Fig. 4b).

#### Tissue lysis and sequencing library preparation

Due to the importance of cell lysis for unbiased microbial characterisation, we evaluated eight lysis formulations on a balanced mock community (S. Note 3.2). Two lysis formulations containing Triton X-100, Lysobac, and Proteinase K outperformed the others, demonstrating minimal lysis bias and providing more accurate composition statistics for the mock community. When applied to intestinal tissue microsamples, these buffers were able to capture the complexity of microbial communities in the caecum and colon, confirming their suitability for compositional analysis (S. Note 3.2).

As a baseline for library-building performance, the manufacturer’s protocol was used to evaluate MSSM’s performance in caecum, colon, and ileum using 5,000 μm^2^ microsamples. Library-building performance, evaluated using sequencing and diversity metrics, was highest in the caecum, lower in the colon, and poorest in the ileum, reflecting the microbial biomass differences across intestinal sections (S. Note 3.3).

#### Design considerations: controls and microsample area

When processing low-biomass microsamples, the risk of environmental DNA contamination and cross-sample carryover poses a significant concern (27). To monitor contamination, 25% of each sequencing batch consisted of randomly arranged negative control reactions, allowing for effective identification and removal of contaminants through data filtering strategies. Although controls yielded sequencing data, they consistently exhibited poorer quality metrics (S. Note 3.4). Notably, a high percentage of reads was discarded due to quality filtering or assignment to adapter and human contamination. Furthermore, applying a minimum genome coverage cutoff of 30% for bacterial quantification to exclude false-positive identifications arising from cross-mapping (28), resulted in 91% of controls retaining no data (S. Note 3.4).

We determined the lower limit of microsample area supported by our methodology using caecum microsamples ranging from 50,000 μm^2^ (∼250 μm in diameter, ∼10,000 bacterial cells) to 500 μm^2^ (∼25 μm in diameter, ∼100 bacterial cells) (Fig. 3a). The median percentage of reads mapped to the reference MAG catalogue was above 80% down to 5,000 μm^2^, reaching 66% at 500 μm^2^ (Fig. 3b). Although alpha-diversity decreased as microsample area was reduced (Fig. 3b), phylogenetic diversity remained largely stable until 1,500 μm^2^, implying that the smaller areas contain fewer taxa, which is expected due to spatial constraints, yet the overall phylogenetic composition is largely maintained (Fig. 3b). Variation between microsamples was primarily driven by their area (Fig. 3c), with the separation being largely influenced by sparsely distributed rare taxa only captured in the larger microsamples (Fig. 3d). Microsamples down to 500 µm² still provide usable data, demonstrating that this sampling scale is technically achievable, although recovering the full community requires denser spatial sampling because very small areas naturally capture communities of lower diversity and higher clonal populations.

**Fig. 3:**
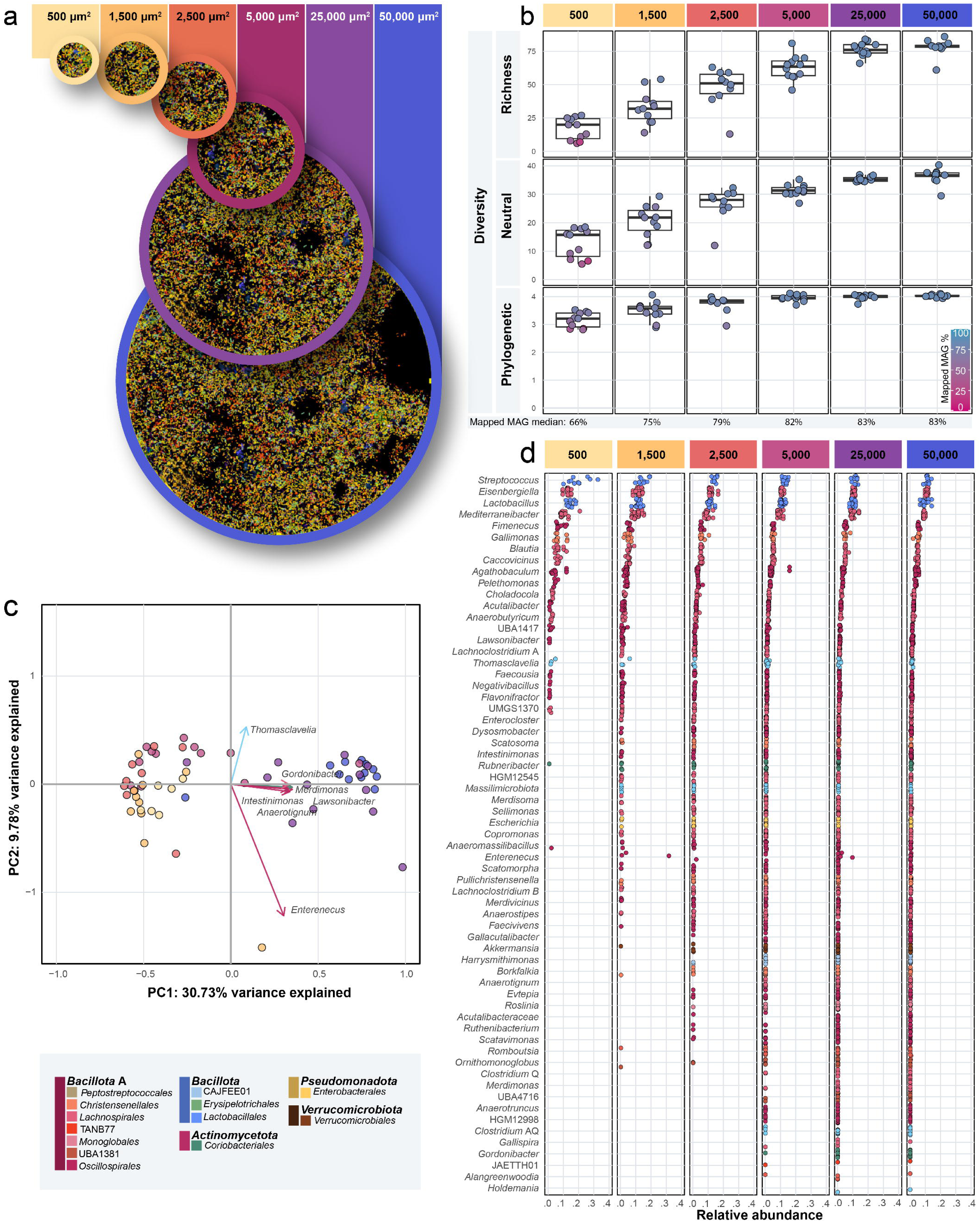
Spatial sampling scale evaluation of micro-scale spatial metagenomics (MSSM). **a,** Overview of the range of microsample areas tested represented over a FISH image obtained from the same tissue. **b,** Boxplots displaying differences in alpha diversity (based on Hill numbers) across microsample areas. **c,** Principal component analysis (PCA) illustrating microbial community variation among microsamples of varying areas, indicated by colour. **d,** Genus-level comparison of relative abundances among microsamples of different areas.

#### Resource optimisation and throughput

To detect spatial patterns of microbiome variation, MSSM inherently requires processing a larger number of samples than conventional metagenomic analyses. To address the need for processing many low-biomass microsamples, we validated half-volume reagent reactions for MSSM library preparation in a scalable 96-well plate format using an automated liquid handler (Fluent, Tecan). Our enhancements maximised resource use, doubling throughput and reducing costs by half, with no notable differences observed in sequencing performance (S. Note 3.5). The optimised workflow, featuring a 2-plate automation routine, enables the generation of 192 sequencing libraries in around 9 hours (S. Note 3.5), positioning MSSM as a promising high-throughput method with significant scalability for future large-scale applications.

### MSSM method validation

#### Discriminative power and replicability of MSSM data

To validate MSSM, we first assessed the variability of caecal micro-scale communities (5,000 μm^2^) reconstructed from animals of different ages (14 and 35 days, Supplementary Fig. 5). MSSM data across individuals revealed highly distinct microbial communities (PERMANOVA, permutations=999, *p=*0.001), underscoring the discriminatory power of our technology. In contrast, differences among sequential cryosections of the same individual, processed in different batches, were negligible (PERMANOVA, *R²* value of 0.008), supporting the reproducibility of our method and indicating minimal batch effects. While PERMANOVA detected some compositional differences between cryosections (999 permutations, *p=*0.042), the significant beta-dispersion (*F*(_3,124_)=12.22, *p=*0.001) suggests that this effect was likely driven by within-cryosection variation rather than true compositional differences. Beyond demonstrating the discriminative power and replicability of MSSM data, these results also show that sequential cryosections harbour similar microbial communities, which facilitates generation of different types of comparable and complementary data, such as spatial metabolomics or FISH imaging (29, 30), from consecutive cryosections.

#### MSSM compared to FISH and macro-scale metagenomics

To evaluate the accuracy of MSSM, we first compared the metagenomic sequencing results from caecum cryosections of 14- and 35-day-old animals with FISH-based quantification. Sequential cryosections were analysed using both MSSM and FISH, and the relative abundances of nine genera targeted by FISH were compared across the datasets. MSSM and FISH showed high agreement in the relative abundance of most genera (Fig. 4a). The variation between MSSM and FISH was minimal, accounting for only 7.5% of the variance, whereas the animal explained 65.3% according to PERMANOVA (Fig. 4b).

**Fig. 4:**
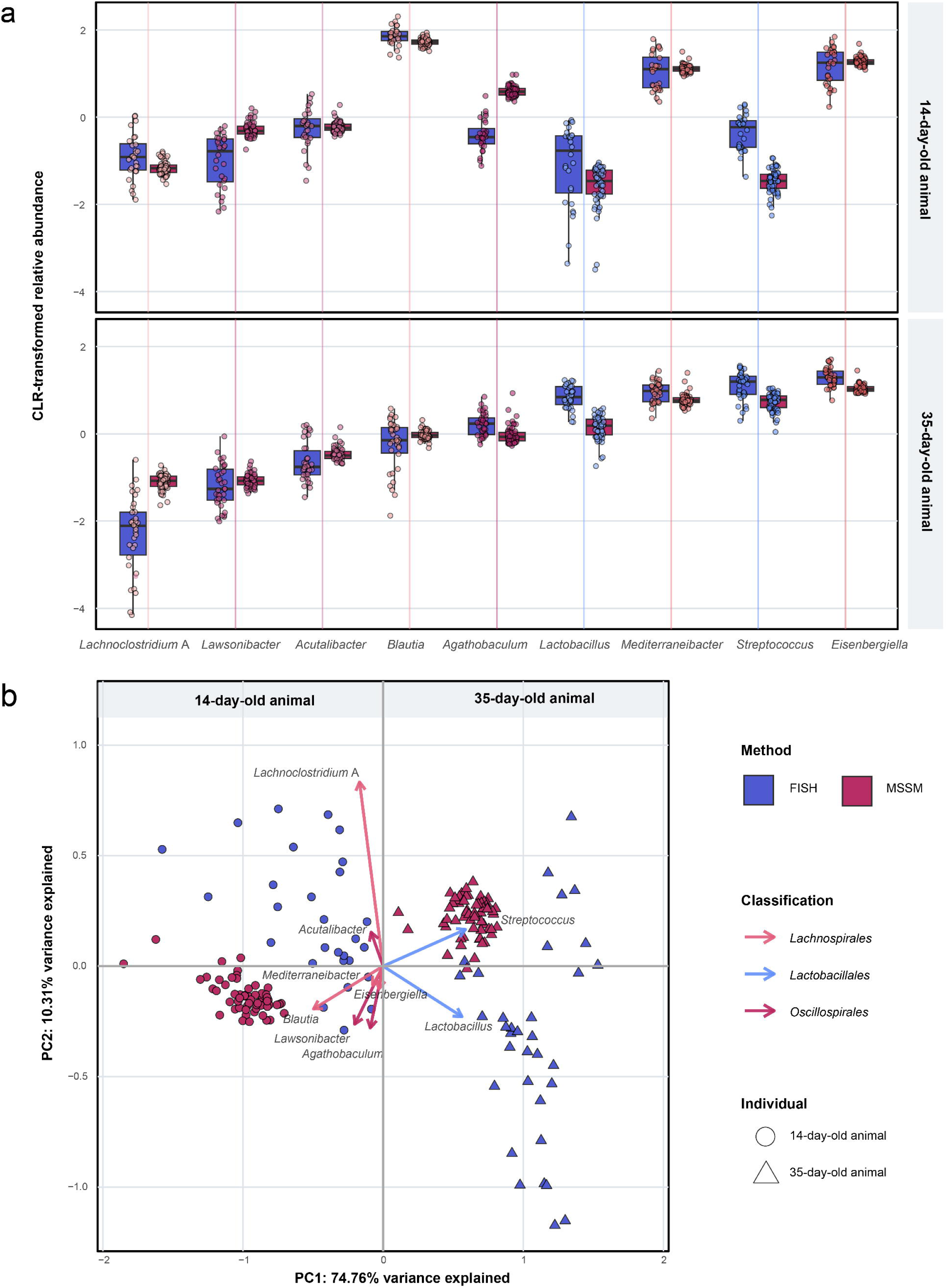
Comparative assessment of bacteria quantification between fluorescence *in situ* hybridisation (FISH) and micro-scale spatial metagenomics (MSSM). a, Centered log-ratio (CLR)-transformed relative abundances of nine bacterial genera quantified using FISH and MSSM in caecal samples from a 14-day- and 35-day-old chicken. **b,** Principal component analysis (PCA) comparing genus-level abundance profiles between FISH (blue, 5,000 μm² area) and MSSM (red, 5,000 μm² microsample). Shape distinguishes the animal: circles represent the 14-day-old and triangles the 35-day-old chicken.

Comparison of the MSSM-derived dataset with macro-scale metagenomic data from the two examined animals revealed a strong correlation in microbial community composition at the species level. In total, 88 species (85%) were shared between the two approaches, while 13 species (13%) and 2 species (2%) were unique to the micro- and macro-scale datasets, respectively. The degree of concordance was influenced by sampling effort: the 35-day-old animal, represented by 263 microsamples, showed 85% overlap, whereas the 14-day-old animal, with 71 microsamples, showed 95% overlap, with two species uniquely identified in the macro-scale dataset. Notably, MSSM identified 15 species absent from the macro-scale dataset in the 35-day-old animal (Fig. 5a). Three of these taxa were consistently detected across microsamples (>30%), suggesting they are prevalent community members that escaped detection in macro-scale sampling due to their low overall abundance.

**Fig. 5.**
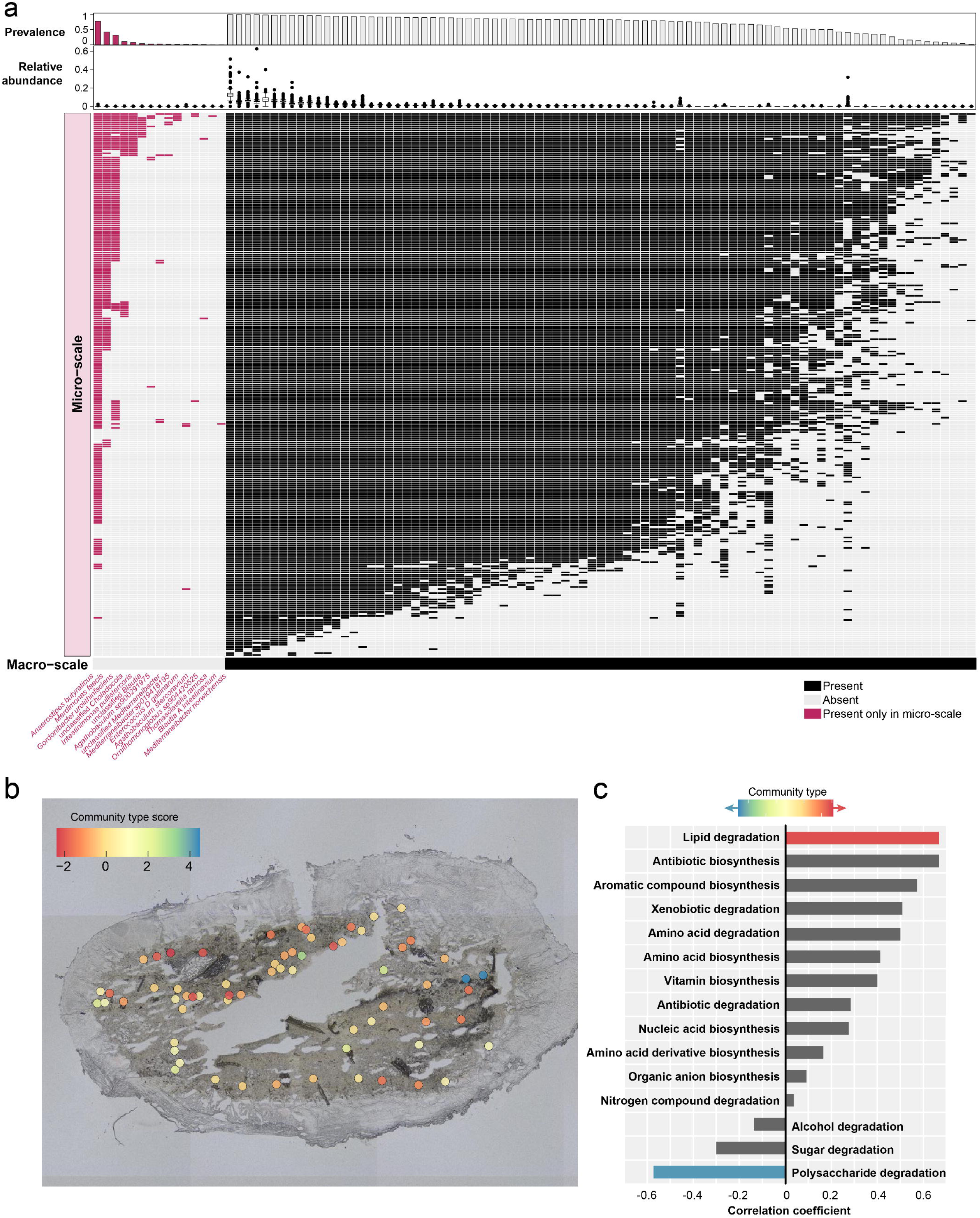
Comparative assessment of bacterial detection using macro-scale metagenomics and micro-scale spatial metagenomics (MSSM), and evaluation of MSSM capacity in colon to resolve spatial taxonomic and functional variation via RLQ analysis. **a,** Species-level presence/absence comparison between the macro-scale metagenomic sample (total caecal content, ∼200□ng input) and 5,000□μm² microsamples obtained via MSSM from a 35-day-old chicken. For each species, bar plots indicate prevalence, and box plots display relative abundances across microsamples. MSSM approach identified 15 unique species, three of which exhibited high prevalence but low relative abundance. **b,** The global coordinates for the first axis are defined as the combination of patterns derived from the environmental variables (microbial alpha diversity and sequence counts) and spatial descriptors. The color indicates the values of site coordinates, ranging from red for strongly negative values to blue for strongly positive values. **c,** For the quantitative functional traits, Pearson correlation (based on raw data) for the first axis between the traits and coordinates of species on the canonical axis are given.

### MSSM method implementation

To demonstrate the scope of our method, we analysed spatial variation in microbial community composition, strain segregation across the highly diverse *Lawsonibacter*, and single nucleotide polymorphism (SNP)-level microdiversity in the most widespread *Lawsonibacter* strains using spatially referenced MSSM data.

#### MSSM enables capturing spatial patterns of taxonomic and functional variation

By breaking down species diversity into three components (within individual microsamples [alpha], between microsamples [beta], and across the whole cryosection [gamma]) we found that diversity within microsamples was lower than expected by chance in both the caecum and the colon. In contrast, the differences in community composition between microsamples were greater than expected by chance, indicating that microbes were spatially organised into distinct, localised groups (S. Note 4). In the colon, dissimilarity between these microbial micro-communities increased with greater spatial separation between microsamples (permutations=10000, *p<*0.001; S. Note 4), whereas no such spatial pattern was observed in the caecum (permutations=10000, *p=*0.110). This likely reflects the structural and functional differences between these two intestinal regions: the colon has a directional flow of contents, promoting spatial separation of microbial communities, whereas the avian caecum serves as a reservoir, where contents accumulate over time (31), reducing spatial structuring.

To further investigate how microbial communities are arranged in the colon, we examined the spatial segregation of functional and phylogenetic characteristics of microbiomes. Our extended RLQ analysis revealed a spatially structured gradient in the microbial community (Fig. 5b), ranging from microsamples with low diversity and sequence counts to those with higher species richness and sequencing depth (S. Note 4). Microsamples with low diversity were typically dominated by *Lachnospirales* and *Lactobacillales* (S. Note 4), and associated with a higher capacity to degrade polysaccharides and sugars (Fig. 5c). In contrast, high-diversity microsamples were dominated by *Oscillospirales* (S. Note 4), with a higher capacity for lipid degradation, suggesting functional segregation of microbial taxa across microsamples (Fig. 5c).

#### Spatial and host distribution of strains resolved by MSSM

To demonstrate MSSM’s ability to detect multiple co-existing strains, we focused on the widespread and abundant *Lawsonibacter*. The eight analysed genomes, carrying different gene contents (Fig. 6a), segregated spatially within and across caecum microsamples (Fig. 6a), although without clear spatial patterning (Mantel statistic: *p>*0.05, permutations=500). Nonetheless, differences in strain representation were evident at the host level, with the 35-day-old animal exhibiting a higher number of co-existing *Lawsonibacter* strains (Fig. 6b), reflecting greater community complexity.

**Fig. 6.**
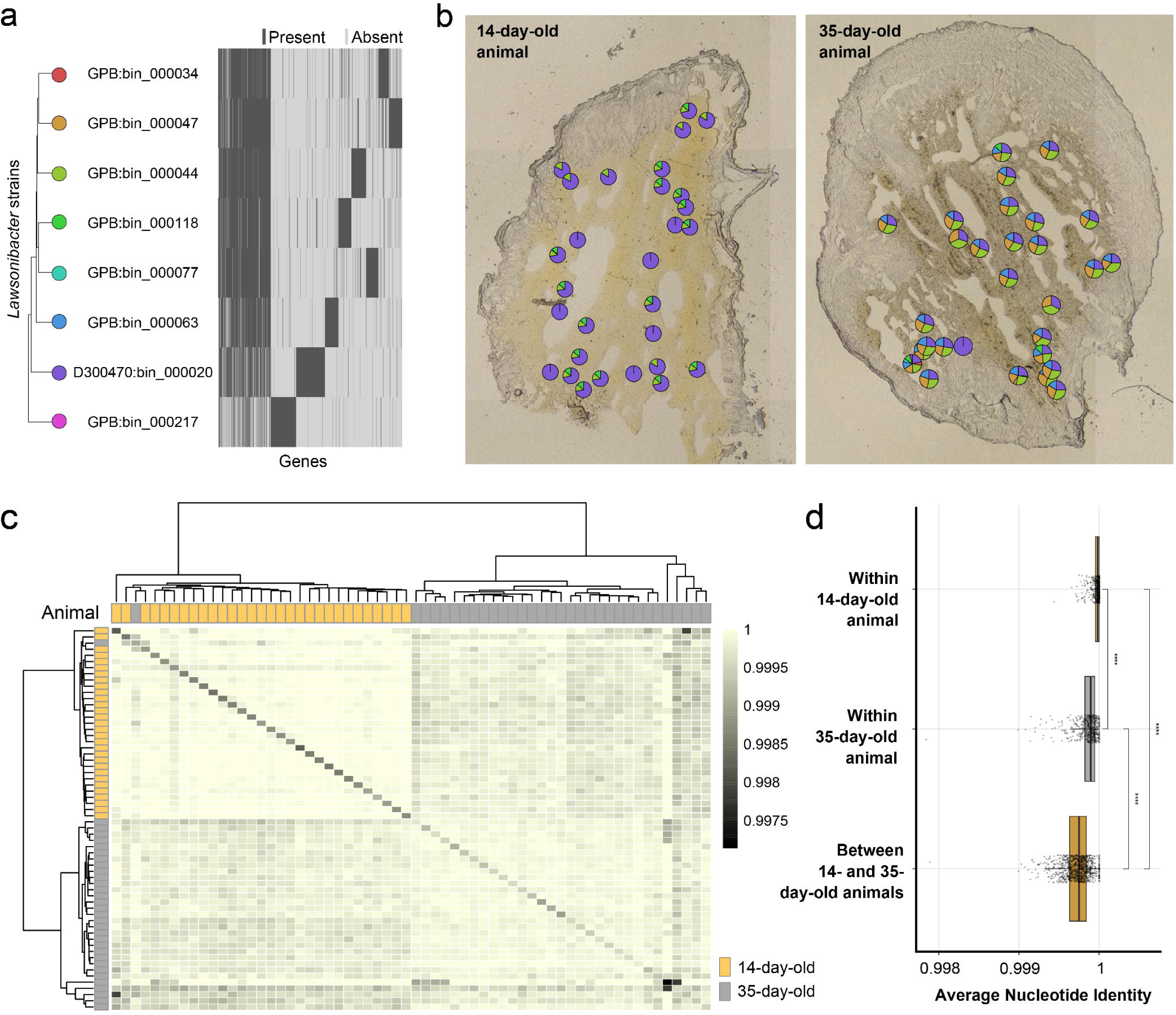
Strain distribution and assessment of within-strain genetic variation using single-nucleotide polymorphisms (SNPs). **a,** Heatmap showing the presence of gene families (x-axis) across *Lawsonibacter* MAGs (y-axis) in the pangenome. Each tile represents a gene family in a specific MAG: dark grey indicates presence, while light gray tiles indicate absence. MAGs are ordered by hierarchical clustering based on shared gene family content, as illustrated by the dendrogram. **b,** Pie charts representing the estimated relative abundance of co-existing *Lawsonibacter* strains within each microsample, based on reads mapping to metagenome-assembled genome (MAG) catalog dereplicated at 98% ANI to represent strains. Each pie chart is plotted according to the spatial coordinates of the corresponding microsample. **c,** Population Average Nucleotide Identity (*pop*ANI) values calculated by Lorikeet for one *Lawsonibacter* strain, across microsamples from a 14-day-old and 35-day-old chicken. Mapping had revealed strong evidence of the presence of this strain in most of the microsamples from the 35-day-old chicken and in a subset of microsamples from the 14-day-old chicken. Pairwise *pop*ANI values were computed between each microsample pair to capture genomic similarity and potential strain-level variation. The diagonal represents the *pop*ANI of each microsample compared to the reference genome, rather than self-comparison. Each microsample is annotated with its corresponding animal host. **d,** Comparison of *pop*ANI values within animals and between animals. Statistical significance between comparison groups was evaluated using Wilcoxon rank-sum tests with Benjamini-Hochberg correction for multiple comparisons. Significance levels are indicated by stars (\**p*<0.05; \*\**p*<0.01; \**p*<0.001 and “ns” for *p*>0.05) shown above relevant comparisons.

#### MSSM can recover spatial variation of SNP-level microdiversity within strains

To examine within-strain microdiversity in the caecum, we analysed SNP-level variation in the most widespread *Lawsonibacter* strains (Fig. 6b). Microdiversity was significantly lower within animals than between animals (Fig. 6c-d, Supplementary Fig. 6), consistent with host-specific colonisation. Notably, microsamples from the 14-day-old animal exhibited lower microdiversity than those from the 35-day-old animal (Fig. 6c-d, Supplementary Fig. 6), suggesting an association with age. In the 35-day-old animal, the Mantel correlogram analysis revealed that, for two of the three *Lawsonibacter* strains, microsamples located within shortest distances (< 200 μm) of each other were significantly more similar than expected by chance (Supplementary Fig. 6). Although the third strain also exhibited this trend, the evidence of a significant signal at the shortest distance was weak (*p*=0.058). These results suggest that variant-level structuring and spatial clustering of clonal populations sharing similar SNP signatures are restricted to the finest spatial scales.

## Discussion

We developed, optimised, and showcased micro-scale shotgun metagenomics (MSSM), a novel molecular technology for high-resolution, spatially-resolved microbiome research, enabling the study of bacterial biogeography at the micron scale. Unlike previous sequencing-based microscale methodologies (16–18, 32), MSSM allows the study of spatial variation in functional metagenomic features at the microscale.

Our MSSM analysis of chicken gut samples revealed differences in the spatial structure across two intestinal sections, the caecum and the colon, likely reflecting variations in bacterial density and diversity. In the colon, our data revealed spatial separation between bacterial consortia primarily capable of carbohydrate degradation and those specialised in lipid breakdown. This implies that local nutrient availability shapes community structure, a hypothesis that could be tested by combining MSSM with spatial metabolomics (29). In the caecum, we showed that bacteria form non-random local aggregates, although these patterns may occur at a finer spatial scale than analysed (5,000 μm^2^), as further supported by microdiversity analysis. This implies that reliable detection of spatial patterns in the chicken caecum requires higher-resolution sampling with smaller microsample sizes, alongside continuous efforts to increase data generation throughput. Our preliminary analyses showed that MSSM could be applied to smaller microsamples; however, fluorescence *in situ* hybridisation imaging revealed substantial spatial heterogeneity in biomass distribution. As a result, further reducing the laser micro-dissection samplingarea may increase the risk of capturing regions with low bacterial biomass, which could potentially affect sequencing efficiency and the completeness of community reconstruction.

Despite these limitations, we were able to resolve the spatial segregation of strains within species at SNP-level. This was made possible by long-read sequencing, which enabled the recovery of circularised reference genomes from macro-scale caecum digesta samples, since long-read library preparation requires much more biomass than short-read methods. Nonetheless, our analyses showed that MSSM can generate a genome catalogue, offering a standalone method for micro-scale genome-resolved metagenomics when circularised genomes are not required.

Our multiple tests across different intestinal sections and host animals showed that MSSM performance depends on the specific properties of the studied environment. Therefore, pilot analyses are necessary to optimise key sampling and data generation parameters based on the microbial diversity, density, and spatial heterogeneity, which are factors that can vary across species, intestinal sections, or treatment groups.

Our comprehensive assessments spanning sample collection, processing, and data analysis establish a robust platform for future testing and refinement across diverse host species and microbial communities. By enabling high-throughput, spatially resolved microbiome profiling, MSSM sets the stage for previously inaccessible insights into host–microbe interactions, microbial niche partitioning, and ecosystem dynamics, reshaping how microbial communities are studied and understood.

## Methods

### Generation of macro-scale metagenome-assembled genomes (MAG) catalogue and metagenomic data

A reference metagenome-assembled genomes (MAG) catalogue was created using the total caecal content from all 114 commercial Ross 308 broilers in the *in vivo* experiment, which provided the intestinal specimens for micro-scale spatial metagenomics (MSSM) analysis. The macro-scale data from two focal animals of different ages (14- and 35-days old), which were also utilised for MSSM development, were subsequently used for comparison in the method validation section.

#### Sample collection and DNA extraction

Details of the *in vivo* experiment can be found in Drauch *et al.* (33). At the time of euthanasia, approximately 100 mg of caecal digesta was collected in 1 mL of DNA/RNA Shield Reagent (Zymo, cat. Number R1100-250) for short-term stabilisation at ambient temperature. Samples were then stored at -18°C until further processing. DNA extraction was performed using the magnetic bead-based method described in Lauritsen *et al.* (34), preceded by mechanical lysis using Lysing Matrix E 96-tube Rack (1.2 mL, MP Biomedicals™, cat. Number 116984010) and two 6-minute bead-beating steps at 30 Hz on a TissueLyser II (Qiagen). The amount of DNA was quantified using the Qubit™ dsDNA Quantification Assay (ThermoFisher, cat. Number Q32851) on a Qubit Flex Fluorometer (Thermofisher, cat. Number Q33327).

#### Short- and long-read sequencing data generation

Different strategies were employed to generate short- and long-read sequencing data. For short-read sequencing, a maximum of 200 ng in 24 μL from each caecal content DNA extract was fragmented using a Covaris LE220R focused ultrasonicator (Covaris, USA) to 320–420 bp-long DNA fragments. A total of 114 individual sequencing libraries were prepared using the optimised version of Blunt-End Single Tube (BEST (35)) protocol described in Mak *et al.* (*36*), with 20 μM BEDC3 adaptors. Libraries were purified using Solid Phase Reversible Immobilisation (SPRI) beads, prepared as described in Rohland & Reich (37), both before and after PCR indexing, with approximately 1.7 and 1.2 times the library volume, respectively. The mixture was incubated for 5 minutes (min) at room temperature, then placed on a 96S Super Magnet (Alpaca, SKU: A001322) to remove the supernatant. The immobilised beads were washed twice with 80% ethanol and air-dried for up to 5 min. DNA was then eluted using Elution Buffer Tween (EBT; Buffer EB, Qiagen, cat. No. 19086, and TWEEN® 20, Sigma-Aldrich, cat. No. P9416-50ML) with a 10-minute incubation at 37°C, followed by recovery of the cleared supernatant. Most of the libraries were amplified using unique dual-index Illumina primers with the following PCR conditions: 1) 1 cycle at 95°C for 12 min, 2) 7 cycles of a) 95°C for 20 seconds, b) 60°C for 30 seconds, and c) 72°C for 40 seconds, 3) 1 cycle at 72°C for 5 min, and 4) hold at 4°C. The PCR reaction mixture consisted of 5 μL of 10x PCR Gold Taq buffer, 5 μL of MgCl2 (25 mM), 0.4 μL of dNTP mix (10 mM each), 1 μL (10 μM) of each P7 and P5 primers, 1 μL of AmpliTaq GOLD DNA polymerase (Applied Biosystems™ cat. Number 4311820), 26.6 μL of sterile dH_2_O, and 10 μL of sample library, for a total reaction volume of 50 μL. Seven libraries showed lower efficiency in library preparation, as assessed by qPCR, and therefore required additional PCR cycles (10 cycles). The final molarity of the indexed libraries was determined using the HS NGS Fragment Kit (Agilent, cat. No. DNF-474-0500) on a Fragment Analyzer (Agilent), and the libraries were then pooled equimolarly. DNA sequencing was performed on an Illumina NovaSeq X Plus platform, using 10B flow cells and 150 paired-end chemistry, aiming for around 4 Gbp (4×10^9^ basepairs) of data per library.

For long-read sequencing, the 14 DNA extracts with the highest diversity and lowest amount of host DNA (based on short-read data) were pooled to create a single library. Individual libraries for each sample were not necessary, as the long-read sequencing only aimed to improve the quality of our MAGs reference genomes. Fragmented DNA (∼7,000 bp) was prepared for PacBio sequencing using the SMRTbell express template prep kit 2.0 (Pacific Biosciences). SMRTbell libraries were bound with sequencing primer v5 and Sequel II DNA Polymerase 2.0 using Sequel II Binding Kit 2.2. Bound complexes were sequenced on a PacBio Revio platform. Circular consensus reads were generated onboard, and only High-Fidelity reads (HiFi, Phred score >Q20) were used for downstream analysis.

#### Macro-scale bioinformatic processing

All methods for generating the macro-scale MAG catalogue, derived from the processing of short- and long-read sequencing data, along with the MAG abundance quantification, are implemented in a Snakemake (38) pipeline, which is accessible on GitHub (https://github.com/3d-omics/mg_assembly).

#### Short-read sequencing MAG Catalogue

Raw reads were trimmed using fastp (39) to remove adapters (5’-AGATCGGAAGAGCACACGTCTGAACTCCAGTCA-3’ and 5’-AGATCGGAAGAGCGTCGTGTAGGGAAAGAGTGT-3’), poly-X fragments, and sequences shorter than 75 bp. Host reads were removed by aligning them to the *Gallus gallus* (Chicken) reference genome (bGalGal1.mat.broiler.GRCg7, GCF_016699485.2) from NCBI using Bowtie2 (40) and Samtools (41). Reads retained after quality filtering and host removal were then assembled with MEGAHIT (42). We omitted samples exhibiting high levels of adapters or poly-G, or high host contamination from the metagenome assembly and used their retained quality-filtered reads only for bacteria quantification by mapping reads against the generated MAG catalogue. Since genomic context matters when producing metagenome assemblies (43), multiple individual assemblies and co-assemblies were performed. This included one individual assembly for each of the 114 animals, one co-assembly per experimental treatment (five total), 25 co-assemblies combining treatment and sampling day (five treatments across five days), and one general co-assembly containing all samples. In total, 145 metagenome (co-)assemblies were performed. Once the reads were assembled into contigs, they were binned (assigning them to specific microbial taxa) by employing multiple binning tools that included MaxBin2 (44), MetaBAT2 (45), and CONCOCT (46), followed by refinement of the binning predictions using MAGsCot (47).

The set of redundant genomes yielded by the multiple (co-)assembly and binning procedures was de-replicated using dRep (48) with a 95% ANI threshold. The resulting dereplicated MAG catalogue was taxonomically classified with GTDB-Tk (49) and functionally annotated with DRAM (50) (database updated on 2023-08-11). MAG’s completeness and contamination assessments were performed with CheckM2 (51). To quantify MAGs in the animals’ total caecal content, reads retained after quality filtering and eukaryotic removal were mapped to the dereplicated MAG catalogue using Bowtie2, and a table of MAG abundances was generated with CoverM (52). These results guided the selection of 14 samples with the lowest host DNA content and highest diversity for long-read sequencing.

#### Long-read MAG Catalogue

HiFi reads were aligned to the *Gallus gallus* reference genome using Minimap (53). Unmapped reads were then processed through PacBio’s HiFi MAG assembly pipeline (54). This Snakemake-based pipeline involves binning with MetaBAT2 and SemiBin2 (55), followed by refinement using DAS Tool (56). Taxonomic classification is performed with GTDB-Tk, and quality assessment is conducted with CheckM2.

#### Macro-scale MAG catalogue and quantification of MAG abundance for macroscale and MSSM comparison

After generating the short- and long-read MAG catalogues, we combined the de-replicated bins from the short-read assemblies with those from long-read sequencing. The combined set was then de-replicated using dRep at 95% ANI to define species-level representatives and 98% ANI to capture subspecies-level diversity. All bins were re-annotated using DRAM and GTDB-Tk, following the annotation workflow outlined above. MAG quantification in the total caecal content of the animals utilised for MSSM comparison was repeated using the updated catalogues, following the previously described approach. To minimise false-positive identifications due to cross-mapping, the resulting MAG abundance table was filtered using a 30% genome coverage threshold (57, 58) before comparison with the MSSM data.

### Micro-scale spatial metagenomics (MSSM)

#### Sample collection

Intestinal sections were obtained from two Ross 308 broiler chickens assigned to the negative control treatment (no *Salmonella* Infantis infection and no synbiotic supplement) in the *in vivo* experiment referred above (33). Most of the MSSM data was generated from an animal euthanised at 35 days of age (id: G121), while the contrast subject (id: G103) for bacterial community comparisons was euthanised at 14 days of age, in compliance with the ethical guidelines for animal experimentation. Immediately post-mortem, a ∼2 cm segment from the ileum, the right caecum, and the colon of each animal was dissected and placed in individual tissue cassettes (Simport™, cat. Number M512), ensuring minimal disturbance to the integrity of the central section by handling the segment at the cut edges. The intestinal sections were rinsed with 50% glycerol to preserve cell integrity before being snap-frozen in liquid nitrogen and long-term ultra-low-temperature (-80°C) storage until further processing.

#### Segmentation, embedding and cryo-sectioning

The frozen ∼2 cm tissue sections were cut into 0.5-0.7 cm sub-segments in a pre-PCR laboratory using a mitre cutter (RP TOOLZ, cat. Number RP-CUTR). A thin layer of the embedding matrix was applied to the cryomolds to mount the sub-segments for cryo-slicing. Each sub-segment was then positioned transversely (perpendicular to the length of the intestinal tract) in the cryomold and immediately transferred to -80°C for 2 min to ensure proper orientation. After freezing, an additional layer of embedding matrix was added to fully submerge the sub-segments, which were then frozen again at -80°C. Cryosectioning was conducted using a Leica CM3050S cryostat (Leica Microsystems, Wetzlar, Germany) with Accu-Edge Low-Profile Disposable Blades CryoSectioning (C35, cat. Number 4810), at a chamber temperature of -26°C and an object temperature of -16°C. The embedded sub-segments were mounted onto the chuck using a fresh embedding matrix, using the tissue shelf heat-extractor to freeze the block onto the chuck. Consecutive cuts of 10 μm of thickness were mounted on PEN (polyethylene naphthalate) membrane frame slides (Leica Microsystems, Wetzlar, Germany), which were kept frozen at -80°C until further processing. As part of our initial methodology development, we compared two embedding matrices with distinct physico-chemical properties: Optimal Cutting Temperature (OCT) compound and 2% Carboxymethyl Cellulose (CMC). Two sub-segments from the same caecal tissue sections were embedded in both OCT and CMC to assess and compare the performance of each embedding material.

#### Laser micro-dissection (LMD)

Laser micro-dissection (LMD) was carried out using a Leica LMD7 microscope (Leica Microsystems, Wetzlar, Germany) placed in a dedicated cold room (10°C). The microscope is equipped with a custom-made enclosure and stage refrigeration systems (Okolab, Naples, Italy) to keep the slide at a maximum temperature of 4°C and physically isolated from the environment during micro-dissecting. This set-up reduces the pace of decay of nucleic acids while minimising the risk of environmental contamination. Moreover, LMD procedures, along with the preceding tissue section processing steps, were performed in a pre-PCR laboratory, minimising the risk of cross-contamination between laboratory processing stages. Laser settings were adjusted to the areas of the microsamples collected for the different experiments conducted in this study (Supplementary Table 1). Areas of interest for sample collection were identified and outlined using Leica microscope software. Microsamples were then collected into 8-strip lids (8 AFA-TUBE TPX Strip Caps, Covaris cat. Number 500639). The microsample collection was validated through direct visualisation under the microscope (Supplementary Fig. 3), and resampling was conducted if the lid was visually confirmed to be empty. The resampling attempts were documented and accounted for in the statistical analysis. A custom script, available at https://github.com/3d-omics/Bioinfo_Micro_LMD_export, was developed to log and visualise LMD information.

#### Method development: tissue lysis and sequencing library preparation

Tissue lysis was achieved in a pre-PCR laboratory using one of the eight different pretreatment methods tested (Table S2, S. Note 3.2), with each method involving an initial incubation at 37°C for 30 min, followed by overnight incubation at 60°C. Prior to further processing, enzyme heat-inactivation was achieved by incubation at 75°C for 30 min. Following tissue lysis and before DNA shearing, 15 μL of sterile dH_2_O was added to the crude lysate. A clean-up step was incorporated into the MSSM workflow after DNA shearing, as preliminary tests indicated it improved library yields (S. Note 3.1). A total of 50 μL of AMPure XP beads (Agencourt) was transferred to the reactions, and the mixture was incubated for 30 min at room temperature, prior to placing it on a 96S Super Magnet (Alpaca, SKU: A001322) and removal of the supernatant. Two bead washes with 80% ethanol were performed, and the beads were air-dried for up to 5 min. DNA was then eluted using 11 μL of sterile dH_2_O, whereas 10 μL of clear supernatant was used as input for library preparation.

MSSM library construction was initially carried out using the workflow and reaction volumes (end-repair, ligation, and qPCR/library amplification) outlined in the User Guide (M01379 V6.1) for the Ovation® Ultralow Library Systems V2 (Tecan, Switzerland), hereafter referred to as the manufacturer workflow. This workflow was applied throughout most of the method development and validation, including testing lysis conditions (S. Notes 3.1 and 3.2), evaluating MSSM performance across different intestinal sections (S. Note 3.3), as detailed in the sections below. Subsequently, we evaluated a workflow that employed half-volume reagent reactions, which was then used to assess the spatial sampling scale of MSSM, assessing the discriminative power and replicability of MSSM data, and utilised an automated workstation (Fluent® Automation Workstation, Tecan) for library preparation (S. Note 3.5). Prior to sequencing, amplified libraries were analysed using a Fragment Analyzer (Agilent) to assess DNA fragment sizes and molarities. Individually indexed libraries were then pooled equimolarly and sequenced across multiple lanes of NovaSeq X 10B flow cells using 150 bp paired-end chemistry.

#### Impact of lysis conditions on library preparation

As part of our initial methodology development, we assessed library construction performance from crude lysates using the manufacturer workflow, and investigated potential inhibition from two of the eight lysis methods listed in Table S3.1: one containing chaotropic salts (L01) and one without (L06). For this experiment (S. Notes 3.1), previously fragmented caecal DNA was diluted to 0.05Cng/μL, and 1 μL of this DNA was added to 9 μL of heat-inactivated lysis solution, yielding a 10 μL reaction volume, prepared in triplicate. These 10 μL crude lysates were used directly as input for library preparation to evaluate lysis-related inhibition. In contrast, for reactions intended for clean-up, 15 μL of sterile dHCO was added, bringing the total volume to 25 μL, and the mixture underwent magnetic bead-based purification using either AMPure XP (Beckman Coulter) or HighPrep PCR (MagBio). Clean-up was performed as described previously, and 10 μL of the recovered clear supernatant was used as input for library preparation. Library construction performance was evaluated by estimating DNA molarity using a Fragment Analyzer (Agilent).

#### Impact of lysis conditions on microbial taxonomic profiling and MSSM performance evaluation across intestinal sections

For the mock community test (S. Notes 3.2), we selected four bacterial species: two Gram-positive (*Microbacterium oxydans* and *Paenibacillus amylolyticus*) and two Gram-negative (*Stenotrophomonas rhizophila* and *Xanthomonas retroflexus*). Each species was cultivated individually in fresh Tryptic Soy Broth (TSB) medium at 25°C with shaking at 250 rpm for 20 hr. Following centrifugation at 5,000 x g for 2 min, the cells were washed twice and resuspended in sterile 1x phosphate buffered saline (PBS). Each monoculture was adjusted to approximately 1 x 10^8^ Ccells/mL with 1x PBS before being fixed with 4% paraformaldehyde (PFA) at room temperature for 15 min. After fixation, the cultures were washed twice and resuspended in sterile 1x PBS. A balanced mock community was then prepared in triplicate (technical replicates), containing approximately 25,000 cells of each species per μL, for a total concentration of 100,000 cells/μL. To evaluate the eight lysis methods (Table S3.1), 1 μL of the mock community was combined with 9 μL of each lysis solution. To detect potential false-positive identifications, each lysis method was also tested on individual monocultures (100,000 cells/μL), with two technical replicates per species.

For the intestinal section comparison (S. Notes 3.3), 36 microsamples were collected from each region of the intestine (ileum, caecum, and colon). This dataset was also used to compare the taxonomic identification outcomes resulting from two lysis protocols (S. Notes 3.2). Each set was evenly divided between two lysis methods, L06 and L07, with 18 reactions per treatment. The comparisons described above were performed using the lysis, inactivation, and DNA fragmentation procedures outlined previously, followed by manual library preparation according to the manufacturer’s workflow and double-indexed adapters (Part No. 0344NB-A01-FG and S02215-FG, Tecan). All libraries were sequenced to a target depth of approximately 4CGB per sample.

#### Resource optimisation and throughput

A total of 14 reactions derived from 5,000 μm^2^ caecum microsamples were used to evaluate the performance of utilising half the standard reagent volume per library preparation (half-reactions, S. Note 3.5). Library preparation was carried out manually following the manufacturer’s workflow, using double-indexed adapters (Part Nos. 0344NB-A01-FG and S02215-FG, Tecan). Subsequently, the validated half-reaction protocol was implemented on a Tecan Fluent automation system (DreamPrep 780). The module and labware software provided by the manufacturer were customised to support reduced reaction volumes and enable processing of a single plate within approximately seven hours. The performance of the automated half-volume library preparation was evaluated using 14 reactions derived from 5,000 μm^2^ caecum microsamples (S. Note 3.5). To enhance throughput, the automated workflow was extended to a two-plate configuration, incorporating interleaved processing steps that increased library preparation capacity, enabling completion within approximately 9 hours. To assess and compare the performance of the one-plate and two-plate automated implementations, human control DNA diluted to 0.1 ng/μL (24 replicates) was used as input for library preparation using single-indexed adapters (Part No. 0344NB-A01-FG and S02366A-FG, Tecan).

#### Design considerations: microsample area

The spatial sampling scale of MSSM in the caecum was evaluated using a range of microsample area: 50,000 μm^2^, 25,000 μm^2^ 5,000 μm^2^, 2,500 μm^2^, 1,500 μm^2^, and 500 μm^2^. According to microscopy-based estimates, the largest microsample area (50,000 μm^2^) corresponds to approximately 10,000 cells. Library preparation was conducted manually using half-reactions and single-indexed adapters (Part No. 0344NB-A01-FG and S02366A-FG, Tecan). For each microsample area, 12 reactions were prepared. Sequencing was carried out for all reactions targeting a depth of approximately 4 GB.

### Micro-scale data processing

#### Quantification of MAGs abundance and micro-scale MAG Catalogue

MAG abundance quantification for most of the MSSM analysis was conducted by mapping retained reads against the macro-scale MAG catalogues using the Snakemake pipeline mg_quant, available at https://github.com/3d-omics/mg_quant. Before quantification, the pipeline applies preprocessing and contamination removal steps, following a similar approach to the “Short-read Sequencing MAG Catalogue” but with key modifications.

To eliminate potential eukaryotic contamination, filtered reads (fastp) were first mapped to the *Gallus gallus* reference genome, followed by additional mapping to the human (GRCh38.p14) and pig (Sscrofa11.1) reference genomes. The inclusion of human and pig references was necessary due to the low-biomass nature of the samples, which increases the risk of human contamination, and because parallel experiments within the project utilise the same LMD instrumentation on swine intestinal sections. In addition to MAG abundance quantification for each microsample, microbial fraction estimation was conducted with SingleM (59).

A sequencing batch, containing data from two animals (72 microsamples per animal) with a target sequencing depth of approximately 2 GB, was used to assess the feasibility of generating a MAG catalogue directly from microsamples (S. Note 3.1). The micro-scale MAG catalogue was constructed following the same approach as the short-read macro-scale catalogue. MAG abundance quantification was performed by mapping reads against either the micro-scale or macro-scale MAG catalogues (S. Note 3.1).

As previously mentioned we employed a 30% genome coverage threshold, requiring at least 30% of a reference genome to be sequenced before it was considered detected, in order to balance detection sensitivity and minimize false positives due to genome overlap.

#### Evaluation of library preparation performance, sequencing efficiency, and microbial community profiles

We evaluated multiple experimental conditions—such as lysis method, intestinal section, and reaction volume—using a suite of metrics to provide an integrated overview of data generation performance. Sequencing efficiency across experimental conditions was quantified based on sequencing yield pre- and post-quality filtering. Library preparation performance was assessed by evaluating the proportion of low-quality reads and the extent of adapter contamination, with the latter defined as the percentage of bases trimmed due to adapter sequences. Environmental DNA contamination was estimated by calculating the proportion of reads aligning to the human genome or other known contaminant references. To determine whether any conditions enhanced bacterial detection, we compared the proportion of reads mapped to microbial reference genomes, quantified microbial fraction using the complementary computational tools SingleM, and assessed within-sample diversity using Hill number-based alpha diversity indices (60). Statistical comparisons between experimental conditions, used as explanatory variables along with microsample collection attempts, were performed within a generalised linear modeling (GLM) framework. Count data were modeled using a quasi-Poisson distribution to account for overdispersion relative to a standard Poisson model. Proportional response variables were analysed using a quasi-binomial distribution, which similarly accommodates overdispersion in binomial data. Continuous response variables were analysed using a linear model (LM) assuming normally distributed (Gaussian) errors. Model assumptions and fit were evaluated through residual diagnostics using simulated residuals, and the significance of explanatory variables was assessed using analysis of deviance with F-tests appropriate to each model family using the Anova() function from the car package (61).

Microbial community composition was compared using beta diversity analyses based on Aitchison distances calculated from centered log-ratio (CLR) transformed abundance profiles Zero counts were imputed prior to CLR transformation using a Bayesian multiplicative replacement strategy implemented via the cmultRepl() function from the zCompositions R package (63). Principal Component Analysis (PCA) was performed on compositionally centered, scaled, and then CLR-transformed data, using the prcomp() function in R to visualise inter-microsample relationships and group-level variance. The influence of explanatory variables on microbial community structure was assessed using permutational multivariate analysis of variance (PERMANOVA) via the adonis2() function in the vegan package (64), with statistical significance determined by the *p*-value and associated *R*² values (65). Differential abundance testing was performed using ALDEx2 (Analysis of Differential Abundance with Exclusion) (66), a compositional data-aware method tailored for microbial count data. Taxa were considered differentially abundant between groups when the q-value was 0.05 after Holm’s multiple testing correction and the associated effect size was equal or above 1. RMSE values were calculated to quantify the deviation between observed and expected species-level relative abundances in the data from the mock microbial community, as a measure of lysis efficiency and compositional bias (S. Note 3.1).

#### Spatial Partitioning and Community Structure Analysis

To explore the spatial scales at which the community varies in the caecum and the colon of the chicken intestines, we performed an additive partitioning of species diversity into alpha, beta, and gamma components (67). Alpha diversity represents the average number of species in a microsample, beta diversity represents the average number of species added when including microsamples within a cryosection, and gamma diversity, which is defined as gamma = alpha + beta, represents average species richness of a cryosection. The partitioning was performed independently for the caecum and the colon. The caecum analysis was based on seven cryosections that ranged between 32-50 microsamples each, whereas the colon analysis used three cryosections that ranged between 28-65 microsamples. The observed richness at each scale was compared with the expected richness computed under a null model, and the null hypothesis of no difference between the observed and expected richness at each scale was tested using a permutation procedure. The additive diversity partitioning was conducted with the adipart() function of the vegan R package (64), and the default “r2dtable” null model was used to compute the expected values.

To assess the spatial structure of microbial community composition within the caecum and the colon we selected the cryosection with the most informative spatial layout. The caecum cryosection contained 50 microsamples and the colon cryosection contained 65 microsamples. First, we built distance decay plots in community similarity, a well known biogeographical pattern (68). To generate these plots, we applied a CLR transformation to MAG sequence counts after replacing the zero counts using a bayesian multiplicative replacement procedure through the cmultRepl() function from zCompositions R package Then, we computed the Euclidean distance between the CLR transformed communities for all microsample pairs within the cryosection. The resulting Aitchison distances were lastly plotted against the spatial distance between sample pairs and tested the null hypothesis of no relationship using a permutation procedure through the lmperm() function from permuco R package (69). It should be noted that, in our analysis the distance decay in similarity is depicted as a distance increase in Aitchison distance. Additionally, we performed the mantel correlogram to test the null hypothesis of no spatial autocorrelation in microbial communities in the caecum and colon cryosections.

Finally, we used an extended version of the so-called RLQ analysis (70) to delve deeper into the spatial patterns of microbial communities within the chicken intestine. A significant spatial structure was only found in the colon cryosection; therefore, the extended RLQ analysis was performed exclusively for the colon. Standard RLQ ordination allows the simultaneous ordination of three tables; linking a trait matrix Q, weighted by species abundance matrix L, to an environmental matrix R. The extended version rlqESLTP utilises five matrices instead (71): the matrix E with environmental variables, the matrix S with spatial variables, the matrix L with microbial community composition and abundances, the matrix T with functional traits of the microbial species and a matrix P with the phylogeny. We therefore related the spatially structured environmental variables (E and S matrices) with phylogenetically structured species traits (T and P matrices), via community composition and abundance (as described by matrix L). The L matrix consisted of Hellinger-transformed MAG sequence counts (the RLQ analysis is based on correspondence analysis of the species matrix and CLR transformation is not compatible with the method). The matrix E consisted of bacterial species richness and log-transformed total sequence count from each sampling unit. The matrix S included X and Y cartesian coordinates of each microsample in the cryosection. This geographic table was used to generate a Gabriel graph and its corresponding connectivity matrix, as suggested by Dray et al. (72). Then, the Moran’s eigenvector maps (MEM) method was applied through the adespatial() R package (73). The method results in a n microsample (rows) x n-1 MEM spatial variables (columns) matrix, which are spatial filters that can be used in canonical analysis models to quantify community spatial structure (74). In the matrix T each element represented pairwise distances among MAGs based on their average functional trait values: since these biological traits are quantitative variables we used the euclidean distance. The last P matrix contained phylogenetic information, where each element consisted of pairwise phylogenetic distances among MAGs. The analysis was conducted using functions described by Pavoine et al. (71) and the ade4 R package (75). Before performing the joint ordination of the five matrices we verified the spatial autocorrelation of species richness and sequence count using the Moran’s I correlation through sp.correlogram() function in vegan, and we evaluated the phylogenetic signal of the functional traits through a root skewness test (76).

#### Strain-Level Differentiation: single nucleotide polymorphisms (SNPs)-based analyses

Leveraging its widespread, high-quality MAGs, we used the genus *Lawsonibacter* as a model to assess whether MSSM-derived sequences could resolve genomic variation at the strain level within two caecum cryosections, surpassing the conventional 95% ANI threshold typically used for species delineation in metagenomic studies. A genome catalogue dereplicated at 98% ANI, containing eight *Lawsonibacter* genomes (6 of which were circularised), was used to quantify strain-level abundance across microsamples. Pairwise community dissimilarities were calculated using the Jaccard index on presence-absence data, and Mantel correlograms were used to test for spatial autocorrelation in strain distributions within each cryosection. Pangenome analysis was performed using PPanGGOLiN (77), which generated a presence-absence matrix of gene families across the assembled *Lawsonibacter* MAGs. While mapping to 98% ANI-dereplicated MAGs enables strain level resolution, it does not capture finer-scale genetic variation such as SNPs, which are key for examining within-strain microdiversity. To address this, we performed microdiversity analysis using Lorikeet (78), a variant-calling tool built as a reimplementation of GATK’s HaplotypeCaller (79). Lorikeet computes metrics such as population ANI (*pop*ANI), allowing for SNP-level comparisons between microsamples. For each circularised *Lawsonibacter* strains detected in most microsamples (three total), we computed pairwise *pop*ANI values and tested for differences within and between animals using Wilcoxon rank-sum tests with Benjamini–Hochberg correction (significance threshold: *p*<0.05). In the 35-day-old animal, we further calculated pairwise genetic distances (1 – *pop*ANI) between all microsample pairs and used a Mantel correlogram to evaluate spatial autocorrelation in SNP-level genetic variation.

### Fluorescence *In situ* Hybridisation (FISH)

#### 16S rRNA Probe Design

16S rRNA gene sequences, obtained either from the SILVA Database (80) or PacBio long-read sequencing, were aligned with MEGA11 (81) using Muscle (82). The aligned FASTA files were then used as input to design genus specific fluorescence *in situ* hybridisation (FISH) probes using DECIPHER package (83). Potential probe sequences were then searched against all 16S rRNA sequences in the target genus. If the probe aligned to the same genetic location of all members of the genus, it was selected and individually tested for signal strength and specificity (Supplementary Fig. 2).

#### FISH on cryosections

Cryosections were prepared and mounted on Tissue Path Superfrost Plus Gold slides, allowed to adhere at room temperature for 5 min and subsequently fixed with 4% PFA at room temperature for 15 min. Bacteria on the slide were then permeabilised using 1 mg/mL of lysozyme and 20 mg/mL of Proteinase K sequentially, incubated at room temperature for 10 and 15 min respectively. Hybridisation buffer (900 mM NaCl, 20 mM Tris-HCl [pH 7.6], 0.01% sodium dodecyl sulfate [SDS], 20% formamide) containing DNA probes was pipetted directly onto each cryosection and incubated at 46°C in a humidity chamber for 3 hours. The slides were submerged in 50 mL of wash buffer (102 mM NaCl, 20 mM Tris-HCl [pH 7.6], 0.01% SDS) and incubated at 48°C for 15 min to wash away non-specific probes. The hybridisation and wash process was repeated with read-out probes when carrying out 2-step FISH. After rinsing and drying the slides, the samples were mounted with Prolong Gold Antifade Mounting Media and sealed with nail polish (Sally Hansen Insta-Dri, B079G4W25R). Slides were incubated in the dark at 4°C before imaging the next day or shortly after. Details on FISH probe testing and optimisation can be found in S. Note 2.

#### Confocal imaging and image analysis

Fluorescence confocal imaging was performed using LSM800 confocal laser-scanning microscope (Zeiss) to generate a villi-to-villi representative tile across the cryosection. Images were processed using the Zeiss ZEN 3.7 Black Edition software. Background correction and image stitching was carried out using BaSiC (84) and Stitching plug-ins in Fiji 2.17.0 respectively (85). Segmentation was carried out with the StarDist plugin in QuPath 0.5.1 (86) on the Alexa Fluor 594 channel to detect single cells tagged by the universal EUB388 probe. Classification and quantification was carried out using QuPath native features (87) using mean intensity measurements from each fluorescent channel.

## Data availability

The raw sequencing and imaging data generated in this study have been deposited in the European Nucleotide Archive (ENA) under Bioproject PRJEB86258. Complementary source data derived from these sequences, along with their associated metadata, are available in Zenodo (doi: 10.5281/zenodo.17091770). The reference genome catalogues at both the macro- and micro-scale are also archived in Zenodo, under DOIs 10.5281/zenodo.17091731 and 10.5281/zenodo.17091749, respectively.

## Code availability

The complete code required to reproduce all analyses presented in this manuscript is available in a dedicated GitHub repository (https://github.com/3D-omics/MSSM) and rendered as an HTML web book at https://3d-omics.github.io/MSSM. A snapshot of the repository has also been archived in Zenodo with doi: 10.5281/zenodo.17144377.

## Acknowledgements

We thank all 3D’omics colleagues and collaborators for their collective contributions, with special appreciation to project manager Ella Z. Lattenkamp for her support at every stage, and for organising both the administrative and scientific aspects of the project.

## Ethics declaration

Approval for the animal trial was given by the institutional ethics committee and licensed by the national authority according to the Austrian law for animal experiments (license number GZ: 2022-0.474.400).

