## Supplementary material for "Micro-scale spatial metagenomics: revealing high-resolution spatial biogeography of gut microbiomes"

Table of contents

**Supplementary figures**	2

**Supplementary tables**	8

**Supplementary notes**	9

1. Comparative analysis of metagenome-assembled genomes (MAG) catalogues	10

2. Design and validation of fluorescence in situ hybridization (FISH) probes	12

3. Micro-scale spatial metagenomics (MSSM) method development	16

1. Impact of lysis conditions on library preparation	18

2. Impact of lysis conditions on microbial taxonomic profiling	19

3. Performance evaluation across intestinal sections	22

4. Design considerations: control reactions	27

5. Resource optimisation and throughput	31

4. Micro-scale Spatial Metagenomics (MSSM) method implementation	34

References	38

### Supplementary figures

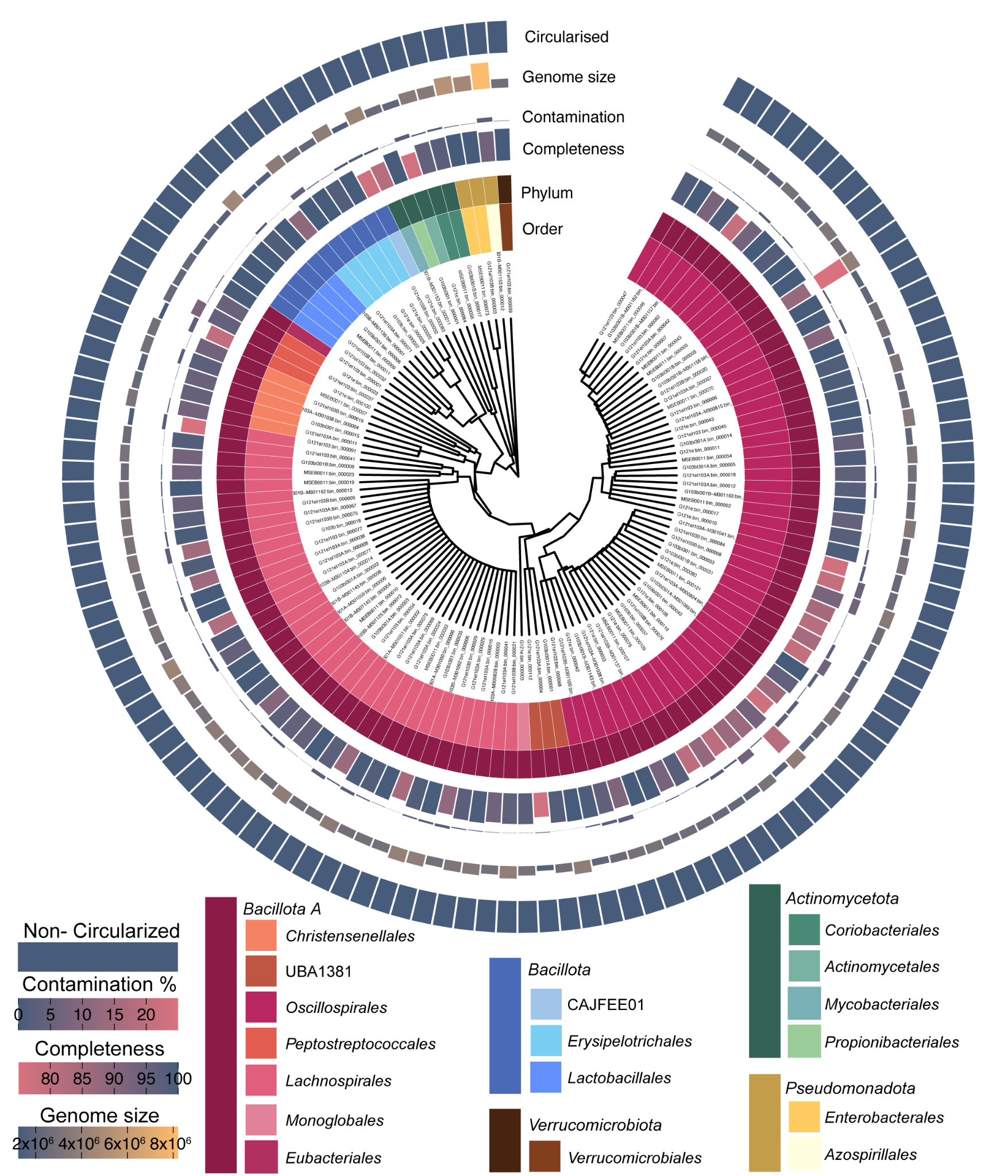

**Supplementary Fig. 1: Metagenome-assembled genomes (MAG) catalogue derived from micro-scale spatial metagenomics (MSSM).** Summary of key features of the micro-scale MAG catalogue, including metrics on completeness, contamination, taxonomic diversity, genome size, and circularisation status.

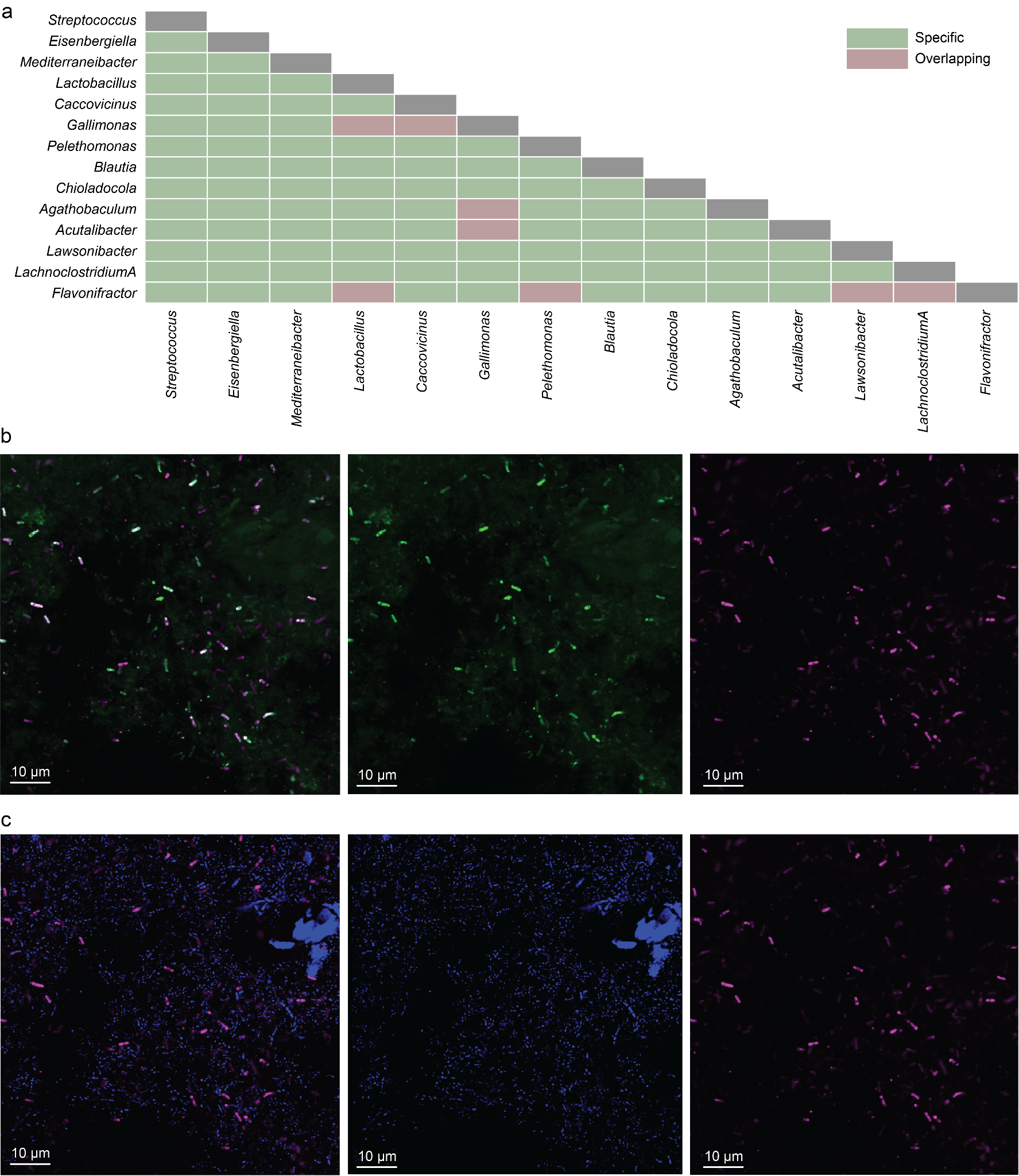

**Supplementary Fig. 2: Validation of fluorescence *in situ* hybridisation (FISH) probe specificity and assessment of cross-reactivity. a,** Heatmap illustrating the specificity and potential cross-reactivity of probes designed for each indicated bacterial genus, based on pairwise testing. **b,** Confocal microscopy image showing hybridisation of probes targeting *Lactobacillus* (pink) and *Gallimonas* (green). Scale bar, 10 μm. **c,** Confocal microscopy image showing hybridisation of probes targeting *Lactobacillus* (pink) and *Caccovicinus* (blue). Scale bar, 10 μm.

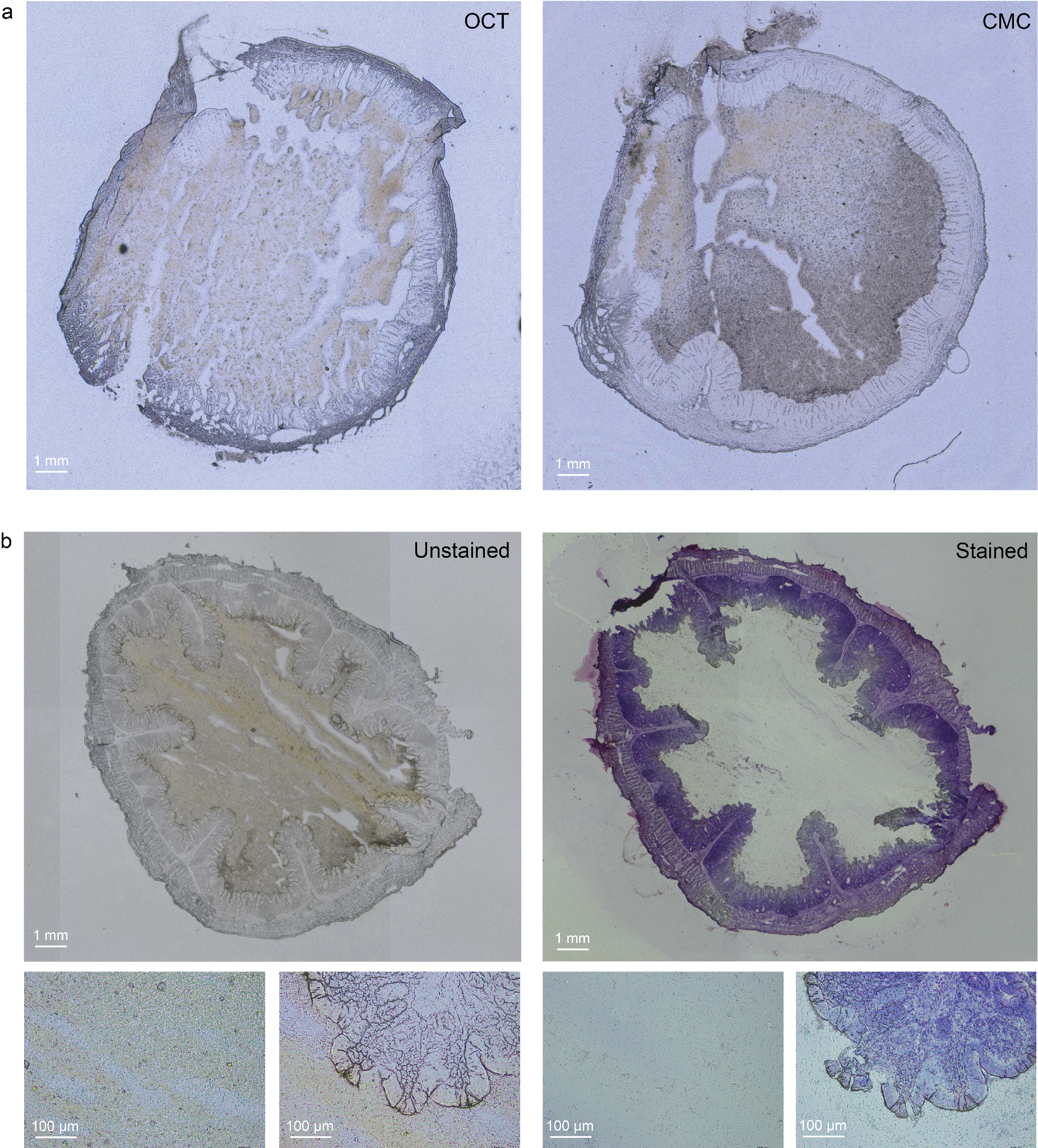

**Supplementary Fig. 3: Comparing embedding material and cryosection processing methods for micro-scale spatial metagenomics (MSSM). a,** Comparison of cryosection from the same tissue segment embedded in optimal cutting temperature (OCT) compound and carboxymethylcellulose (CMC), showing no observable differences in section quality or tissue morphology. **b,** Comparison of the same cryosection before and after Cresyl Violet staining (top). Enlarged views (20x magnification) of the lumen and villi regions from the same cryosection (bottom).

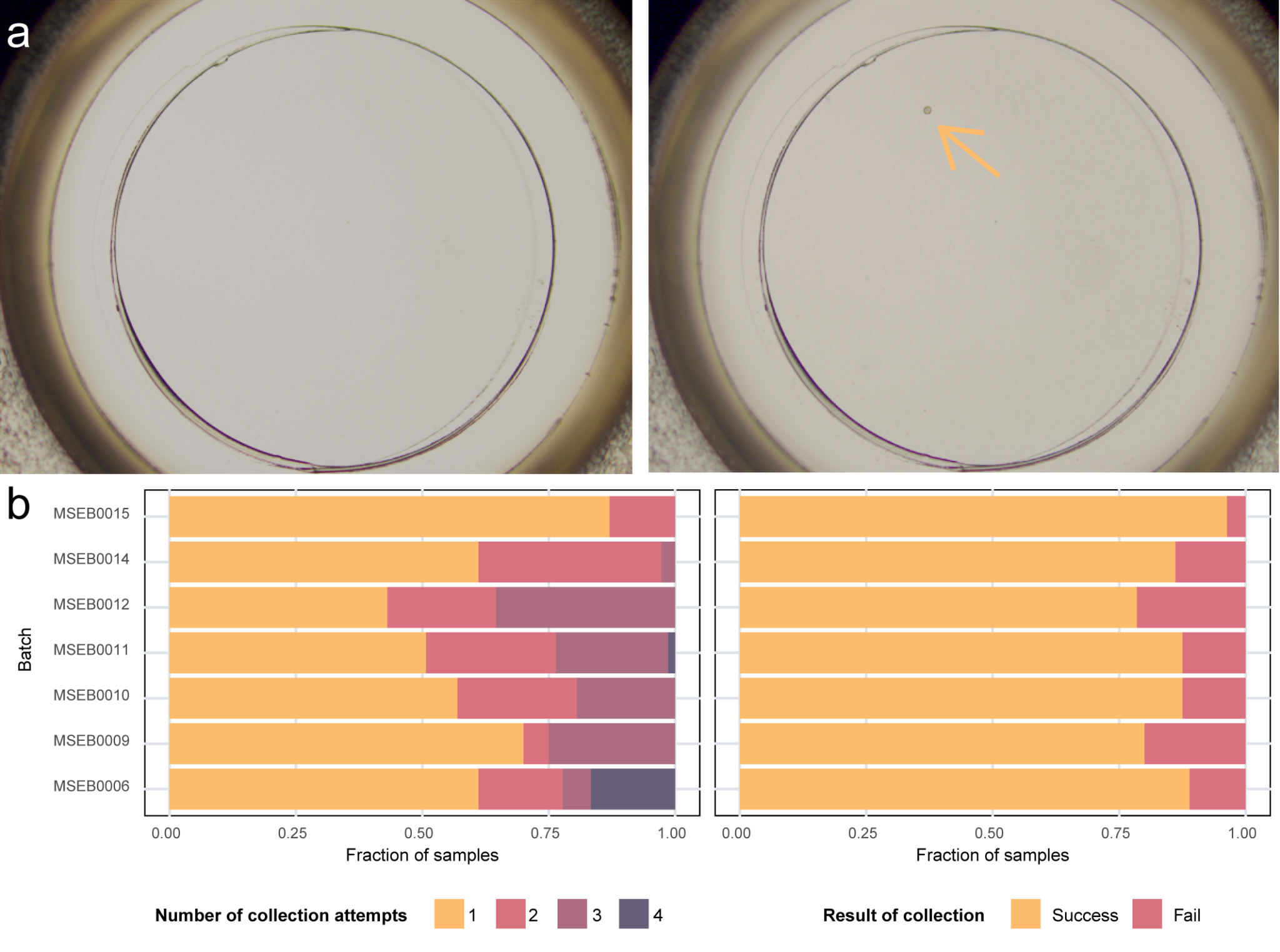

**Supplementary Fig. 4: Microsamples collection using 8-strip cap lids. a,** Visual confirmation of achieved microsamples collection. Left: Empty lid (approximate diameter of 5 mm) without microsamples; Right: Lid with one microsamples (size 5,000 μm^2^), indicated by the arrow. **b,** Comparison of the number of collection attempts and successful microsection collection across batches. Left: Bar plot showing the fraction of sequencing reactions based on the number of collection attempts, separated by sequencing batch. Right: Bar plot illustrating the fraction of successful microsection collections, confirmed visually, and separated by sequencing batch.

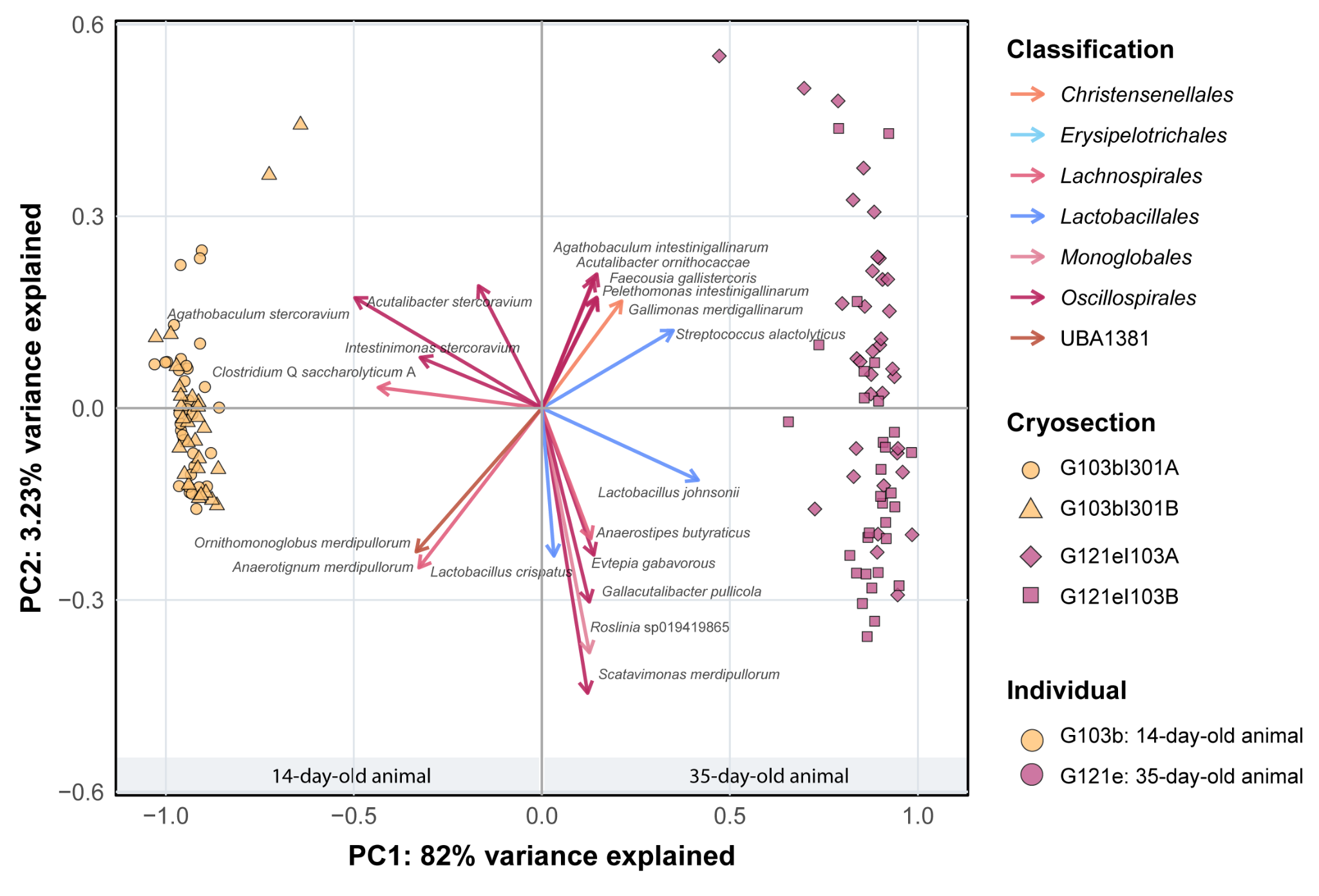

**Supplementary Fig. 5: Principal Component Analysis (PCA) illustrating variation in caecal micro-scale microbial communities, based on 5,000 μm² microsamples.** Each point represents a microsample, with the shape indicating the cryosection of origin. The points’ colour indicates the animal: yellow for the 14-day-old individual, pink for the 35-days-old one. The PCA illustrates both age-related differences and variation between sequential cryosections from the same individual.

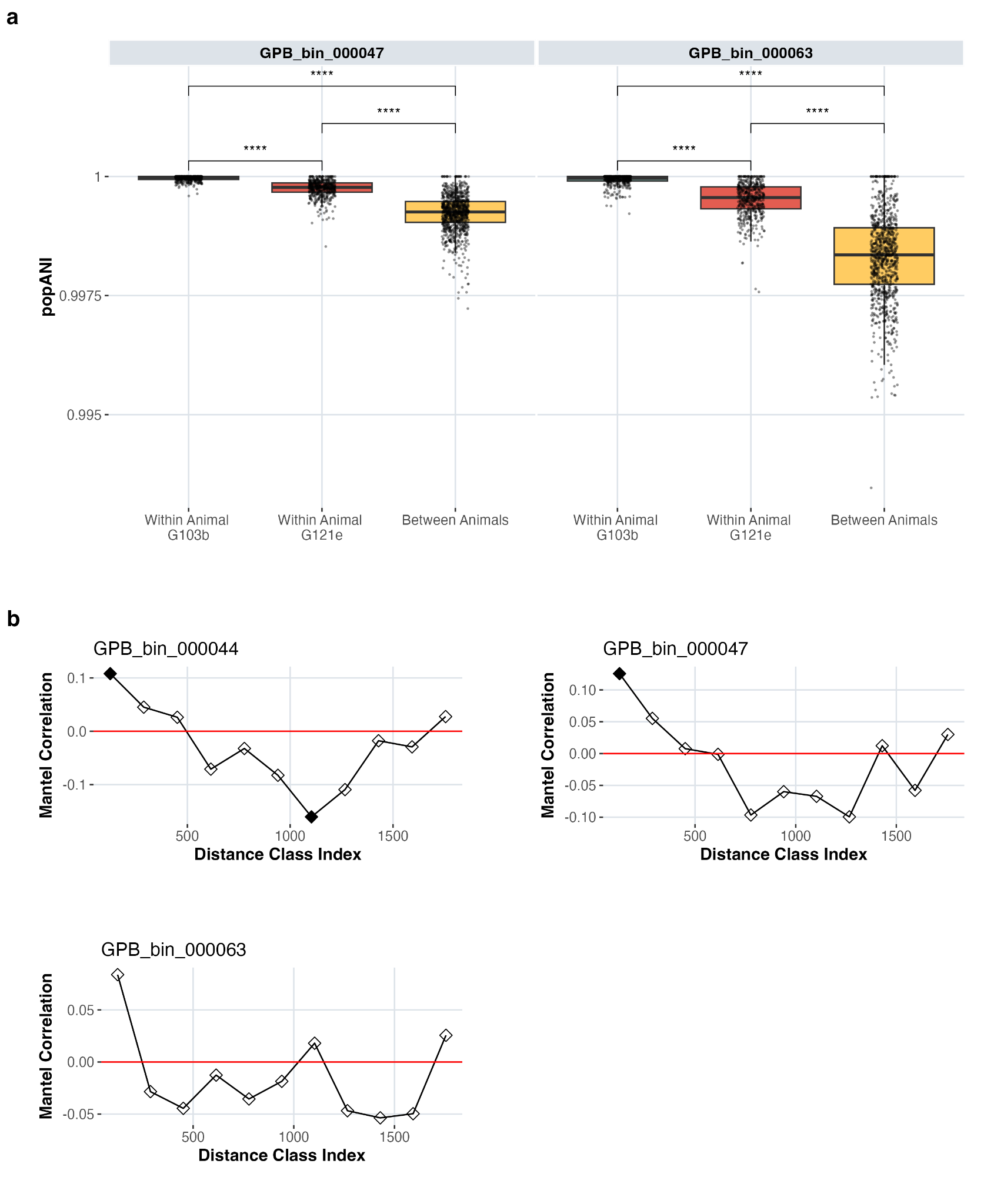

**Supplementary Fig. 6: Assessment of within-strain genetic variation using single-nucleotide polymorphisms (SNPs). a,** Comparison of Population Average Nucleotide Identity (*pop*ANI) values detected within animals and between animals (G103 indicates the 14-day-old animal, G121 indicates the 35-day-old animal). Statistical significance between comparison groups was evaluated using Wilcoxon rank-sum tests with Benjamini-Hochberg correction for multiple comparisons. Significance levels are indicated by stars (*p<0.05; **p<0.01; *p<0.001 and “ns” for p>0.05) shown above relevant comparisons. **b,** Mantel correlogram showing spatial autocorrelation of *pop*ANI distances across defined distance classes; filled squares indicate significant autocorrelation.

### Supplementary tables

**Supplementary Table 1: Laser settings for laser micro-dissection (LMD) on the LMD7 (Leica Microsystems, Wetzlar, Germany).**

| **Cryosection areas (μm^2^)** | **Objective** | **Power** | **Aperture** | **Speed** | **Specimen**  **Balance** | **Head**  **Current (%)** | **Pulse Frequency (Hz)** |
| --- | --- | --- | --- | --- | --- | --- | --- |
| **≥ 100,000** | 2.5x | 59 | 45 | 7 | 25 | 100 | 120 |
| **30,000 - 100,000** | 10x | 42 | 28 | 12 | 15 | 84 | 120 |
| **5,000 - 30,000** | 20x | 52 | 20 | 10 | 15 | 100 | 120 |
| **2,000 - 5,000** | 40x | 15 | 28 | 10 | 20 | 100 | 120 |
| **500 - 2,000** | 63x | 45 | 35 | 10 | 7 | 46 | 200 |
| **≤ 500** | 150x | 10 | 1 | 15 | 5 | 62 | 120 |

### Supplementary notes

#### 1. Comparative analysis of metagenome-assembled genomes (MAG) catalogues

The number of genomes (122) reconstructed from 5,000 µm² caecum microsamples was lower than that from macro-samples (223). This result was expected, as the macro-scale catalogue was created using data from all animals in the *in vivo* experiment, while the micro-scale catalogue was based on samples from just two animals. We compared the effectiveness of the catalogues in capturing microbiome complexity by mapping reads from the same sequencing batch used to generate the micro-scale MAG catalogue to both macro-scale and micro-scale catalogues. Read mapping results were strongly correlated across the two catalogues (Spearman's ρ = 0.99, *p<*0.001, n = 144), with marginally higher read counts observed for the micro-scale genome catalogue. However, a paired Wilcoxon signed-rank test revealed significant differences in alpha diversity between the two reference catalogues. Specifically, the median richness was significantly higher when the reads were mapped to the micro-scale catalogue compared to the macro-scale one (n=141; richness: *V*=7486.5, *p*<0.001 Fig. S1.1a). Up to 40 species were uniquely identified through mapping against the micro-scale catalogue, compared to only 8 species exclusive to the macro-scale one. Additionally, the micro-scale catalogue revealed 28 unique genera, whereas the macro-scale catalogue did not detect any unique genera. The most variation at the species level between catalogues was observed in the 35-day-old animal (G121e, Fig. S1.1b). This separation was mainly driven by species absent from either catalogues, except for a *Lawsonibacter* unclassified species that displayed differential abundance (CLR scale).

The main benefit of the macro-scale catalogue were the circularised genomes required for designing effective 16S rRNA fluorescence *in situ* hybridisation (FISH) probes and enabling strain-level resolution analyses. However, the data above demonstrate the feasibility of creating a comprehensive reference catalogue that captures functional microbiome attributes directly from micro-scale spatial metagenomics (MSSM) data generated from laser micro-dissection (LMD) of the caecum at 5,000 µm².

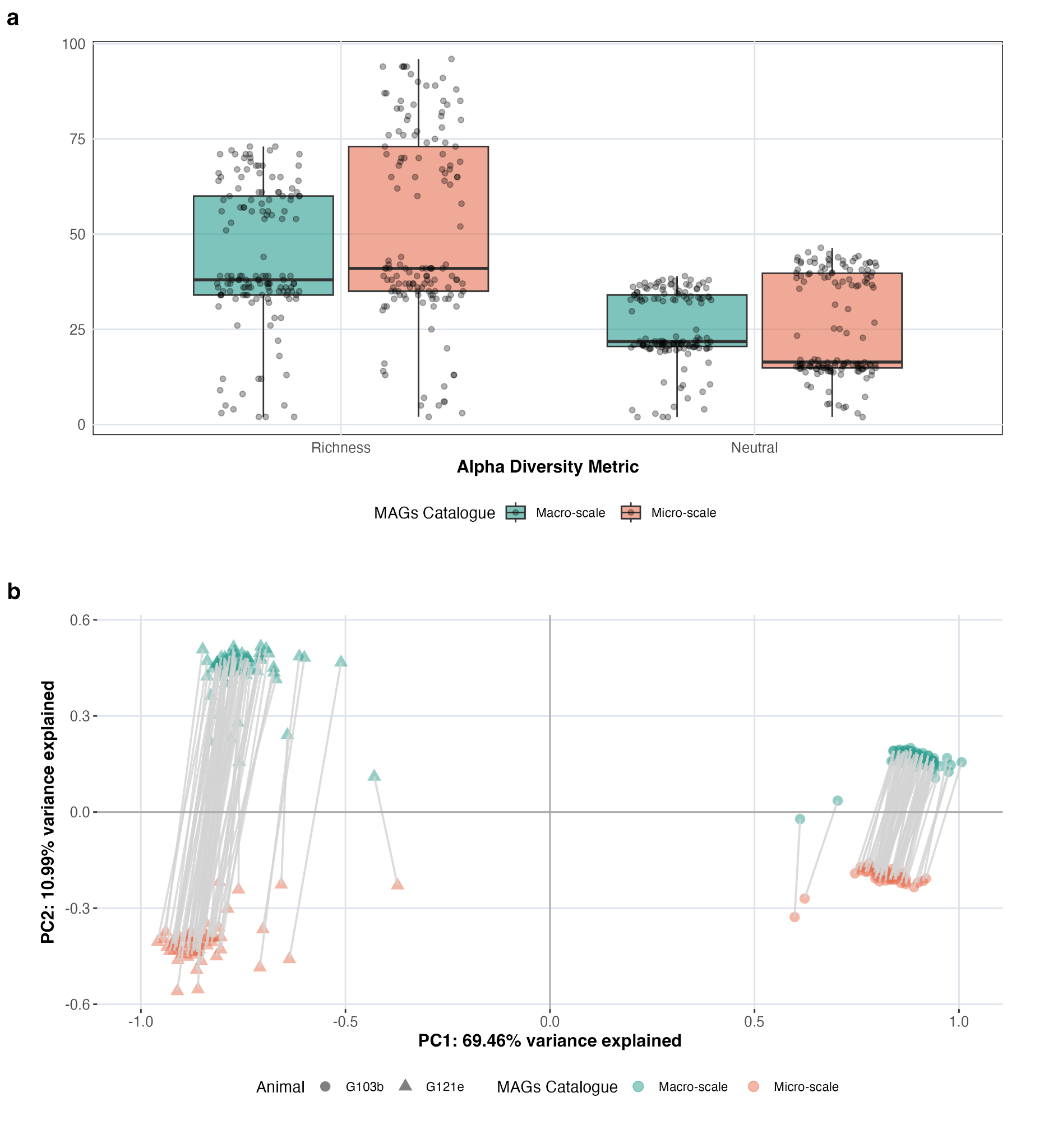
**Fig. S1.1: Comparative analysis of microbial community profiling using micro- and macro-scale MAG catalogues. a,** Richness and Neutral diversity indices based on read mapping to each reference catalogue. **b,** Principal Component Analysis (PCA) illustrating variation in bacterial community composition at the species level, with lines connecting paired samples across mapping strategies.

#### 2. Design and validation of fluorescence *in situ* hybridization (FISH) probes

To balance signal strength and cost between single-probe (CLASI-FISH (1)) and dual-probe (HiPR-FISH (2)) approaches, we adopted a hybrid strategy for probe development and validation. Initially, to economize on hybridization performance testing, we used a two-probe system: an untagged probe complementary to the target 16S rRNA sequence, linked via a 3-bp spacer to a unique barcode, followed by a second fluorophore-tagged probe complementary to that spacer barcode for “read-out” (Table S2.1). After confirming probe specificity, we switched to directly fluorophore-labeled versions of the same probes to enhance signal intensity and simplify image analysis. The initial design of FISH probes was conducted based on 16S rRNA sequences available in the SILVA database (3). The reference sequences available in the database corresponded to bacteria in the same genera but not species directly found in the chicken gut. Among the 15 most abundant genera detected in our samples, 16S sequences were available for 10 of them. However, after testing 45 different probes, a positive signal was only obtained for 5 out of the 10 genera (Table S2.2). This likely reflects the greater 16S rRNA sequence diversity of chicken gut bacteria compared to those represented in available databases. This is evident from sequence alignments, which highlight the mismatches and help explain why these probes failed to detect their targets in the chicken gut (Fig. S2.1).

**Table S2.1: Information on read-out probes utilised in the two-probe strategy.** Probes BW17 to BW154 were purchased from IDT, while BW155 was obtained from Thermo Fisher Scientific. Excitation and emission wavelengths are reported in nanometers (nm). The "Site" column indicates the oligonucleotide terminus (5' or 3') to which the fluorophore is conjugated.

| **No.** | **Name** | **Sequence** | **Fluorophore** | **Site** | **Excitation** | **Emission** |
| --- | --- | --- | --- | --- | --- | --- |
| **BW17** | AF488 | TATCCTTCAATCCCTCCACA | Alexa Fluor 488 | 5' | 490 | 525 |
| **BW18** | AF546 | ACCACAACCCATTCCTTTCA | Alexa Fluor 546 | 5' | 561 | 572 |
| **BW19** | AF594 | TTCTCCCTCTATCAACTCTA | Alexa Fluor 594 | 5' | 590 | 618 |
| **BW20** | AF647 | TTTACTCCCTACACCTCCAA | Alexa Fluor 647 | 5' | 650 | 671 |
| **BW155** | PaBl | ACTCCACTACTACTCACTCT | PacificBlue | 5' | 404 | 455 |

Since long-read sequencing enabled the circularisation of multiple MAGs per genus, we aligned each 16S rRNA sequence captured, chose a representative strain to run DECIPHER against, and selected probe targets that would hybridise across all MAGs in the genus. This strategy proved more successful, resulting in a significantly higher probe success rate (75.5%, Table S2.3). Each probe that gave a signal was subsequently assessed for specificity in a cross-reactivity matrix (Supplementary Fig. 2a). Probes targeting the top 12 genera were then employed simultaneously. However, we found that 3 sets of probes (targeting *Pelethomonas*, *Caccovicinus*, and *Choladocola*) that had generated positive signals when used individually, produced close to background-like signals when combined, making automated identification inaccurate. Consequently, these probes were excluded from quantification.

Additionally, to streamline image analysis of multiplexed FISH experiments, we implemented an image analysis pipeline combining multiple open-source and user-friendly platforms. Cryosections hybridised with fluorescent probes were imaged villi-to-villi using confocal microscopy to generate a representative tile across the cryosection (Fig. 1). Illumination and background correction were carried out using the BaSiC plugin in Fiji before being imported into QuPath. Segmentation was carried out on the Alexa Fluor 594 channel using the StarDist plugin in QuPath to get objects of single cells detected by the universal EUB388 probe. Classification and quantification were then completed using the default QuPath interfaces using mean intensity measurements from the other 4 channels (Fig. 1).

**Table S2.2: Probes designed and tested on publicly available 16S rRNA sequences (silvaDB).** In the SG column, "Y" indicates the presence of a detectable signal above background noise, while "N" denotes the absence of such a signal. In the SP column, "Y" confirms specific binding to the target genus, whereas "N" indicates non-specific signals. An example of this is shown in Supplementary Fig. 2b**.**

| **No.** | **Probe Name** | **Sequence** | **Target genus** | **SG** | **SP** |
| --- | --- | --- | --- | --- | --- |
| **BW88** | Bork 488 | CACGTGAAGTCGGAGTTGCTAGTA tgtggagggattgaaggata | *Borkfalkia* | **N** |  |
| **BW89** | Blaut 546 | GACTGCACGAAGCTGGAATCGCTA tgaaaggaatgggttgtggt | *Blautia* | **N** |  |
| **BW90** | Medite 647 | GTCACTTCACTTTCCTTCCG ttggaggtgtagggagtaaa | *Mediterraneibacter* | **N** |  |
| **BW92** | Bork 488 II | GTCCTGGGCTACACACGTG gtg tgtggagggattgaaggata | *Borkfalkia* | **N** |  |
| **BW93** | Blaut 546 II | ACAAAGGGAAGCGAGCCTGCGA cga tgaaaggaatgggttgtggt | *Blautia* | **N** |  |
| **BW94** | Medite 647 II | CTCCGAAGAGAAGGCGACATTACTCG tcg ttggaggtgtagggagtaaa | *Mediterraneibacter* | **N** |  |
| **BW95** | Eisen 594 II | ACAGGTGCGTCATTGGGATGT tgt tagagttgatagagggagaa | *Eisenbergiella* | **N** |  |
| **BW98** | Eintest 594 | TTTGATTCCATCCGAAAACTTCCTCG tcg tagagttgatagagggagaa | *Eisenbergiella* | **N** |  |
| **BW99** | Eintest 594 II | GGTCCATACCCACCTACAC cac tagagttgatagagggagaa | *Eisenbergiella* | **Y** | **Y** |
| **BW100** | Amuri 546 II | GTACTTCATCAGCGAATATCTCCGT cgt tgaaaggaatgggttgtggt | *Acutalibacter* | **N** |  |
| **BW101** | Gintest 488 | CCTCTCTGTGGTCATATGCG gcg tgtggagggattgaaggata | *Gallimonas* | **N** |  |
| **BW102** | Gintest 488 II | CGCTGCCAACTTGCCAAACC acc tgtggagggattgaaggata | *Gallimonas* | **N** |  |
| **BW103** | Adesm 647 | CCACTAAGTTATGAATTCCATCTCCG ccg ttggaggtgtagggagtaaa | *Agathobaculum* | **N** |  |
| **BW104** | Adesm 647 II | GCCACTAAGTTATGAATTCCATCTCC tcc ttggaggtgtagggagtaaa | *Agathobaculum* | **N** |  |
| **BW105** | Lmass 546 | ATTCCATCTCCGAAGAGATTTCCAA caa tgaaaggaatgggttgtggt | *Lachnoclostridium* | **N** |  |
| **BW106** | Lmass 546 II | CAAGGATGCCCCCAAAACGTATTATG atg tgaaaggaatgggttgtggt | *Lachnoclostridium* | **N** |  |
| **BW107** | Fplau 594 | AGTCACTTAAGCTTCACCCC tag tagagttgatagagggagaa | *Flavonifractor* | **N** |  |
| **BW108** | Fplau 594 II | GGCACTACTGTTTTATGCGGTG gtg tagagttgatagagggagaa | *Flavonifractor* | **N** |  |
| **BW109** | Mmass 647 | CATACCACCTCAGTTTTTACCT cct ttggaggtgtagggagtaaa | *Mediterraneibacter* | **N** |  |
| **BW110** | Mmass 647 II | GCTCAGTCACTTCACTTTCC tcc ttggaggtgtagggagtaaa | *Mediterraneibacter* | **N** |  |
| **BW111** | Borth 488 | CCCCTTATGATTTGGGCTACAC cac tgtggagggattgaaggata | *Blautia* | **N** |  |
| **BW112** | Borth 488 II | TCCCAAAAATAACGTCCCAGTTCGG cgg tgtggagggattgaaggata | *Blautia* | **N** |  |
| **BW113** | Lasa 546 | CTTTCTAAGGCACTCCGTTCGAC gac tgaaaggaatgggttgtggt | *Lawsonibacter* | **Y** | **N** |
| **BW114** | Lasa 546 II | TGGACGAATCCTCTTTCTAAGGCA gca tgaaaggaatgggttgtggt | *Lawsonibacter* | **Y** | **N** |
| **BW115** | Bceft 594 | GCAACCCGCCCACGTGAAG aag tagagttgatagagggagaa | *Borkfalkia* | **N** |  |
| **BW116** | Bceft 594 II | GATAGTCTCAGTTCGGATCGTGG tgg tagagttgatagagggagaa | *Borkfalkia* | **N** |  |
| **BW120** | Lmass 594 III | GTTGTGTCATGCGGCACTACTG ctg tagagttgatagagggagaa | *Lachnoclostridium* | **N** |  |
| **BW121** | Lmass 594 IV | CCGACGTTACTCGGCTGTCAAAGG agg tagagttgatagagggagaa | *Lachnoclostridium* | **N** |  |
| **BW122** | Fplau 594 III | GGTAACTCAATCCATTGGACGAATCC tcc tagagttgatagagggagaa | *Flavonifractor* | **Y** | **N** |
| **BW123** | Fplau 594 IV | CGCCACTAGGTAACTCAATCCAT cat tagagttgatagagggagaa | *Flavonifractor* | **N** |  |
| **BW124** | Cacco II 488 | CTCCGAAGAGAAGGCGACATTACTCG tcg tgtggagggattgaaggata | *Caccovicinus* | **Y** | **N** |
| **BW126** | Medite 546 II | CTCCGAAGAGAAGGCGACATTACTCG tcg tgaaaggaatgggttgtggt | *Mediterraneibacter* | **N** |  |
| **BW127** | Mfae 546 | CTCAGTCACCAAATTTTTCATTCCGA tgaaaggaatgggttgtggt | *Mediterraneibacter* | **N** |  |
| **BW128** | Adesm III 488 | CTTTGACGTTTCAAGGATGCCC ccc tgtggagggattgaaggata | *Agathobaculum* | **N** |  |
| **BW129** | Adesm IV 488 | GTTTCAAGGATGCCCCCAAAACG acg tgtggagggattgaaggata | *Agathobaculum* | **N** |  |
| **BW130** | Mmass III 546 | AGACTTGCTGCTCCGTCT tct tgaaaggaatgggttgtggt | *Mediterraneibacter* | **Y** | **N** |
| **BW131** | Mmass IV 546 | TCTCGGCTGCTCCGAAGAGAAG aag tgaaaggaatgggttgtggt | *Mediterraneibacter* | **N** |  |
| **BW132** | Borth III 594 | GGACTGCAGTCTGCAACTCGAC gac tagagttgatagagggagaa | *Blautia* | **N** |  |
| **BW133** | Borth IV 594 | GTCAAATCATCATGCCCCTTATGATT att tagagttgatagagggagaa | *Blautia* | **N** |  |
| **BW134** | Bceft III 594 | CTTACGTCCTGGGCTACACA aca tagagttgatagagggagaa | *Borkfalkia* | **N** |  |
| **BW135** | Bceft IV 594 | GGATCGTGGGCTGCAACC acc tagagttgatagagggagaa | *Borkfalkia* | **N** |  |
| **BW138** | Cacco II 594 | CTCCGAAGAGAAGGCGACATTACTCG tcg tagagttgatagagggagaa | *Caccovicinus* | **N** |  |
| **BW140** | Meditte II 546 | CTCCGAAGAGAAGGCGACATTACTCG tcg tgaaaggaatgggttgtggt | *Mediterraneibacter* | **Y** | **N** |
| **BW142** | Gintest 488 III | GAGCTCATCCATGTCCGATAAAATCT tct tgtggagggattgaaggata | *Gallimonas* | **N** |  |
| **BW143** | Gintest 488 IV | ACAAGGATGGTCACGCGATGTCAAG aag tgtggagggattgaaggata | *Gallimonas* | **N** |  |
| **BW144** | Adesm 488 V | GATGCCCCCAAAACGTATTATGC tgc tgtggagggattgaaggata | *Agathobaculum* | **N** |  |
| **BW145** | Borth V 594 | ACTGCAGTCTGCAACTCGACTG ctg tagagttgatagagggagaa | *Blautia* | **N** |  |
| **BW146** | Borth VI 594 | GGAGCGAATCCCAAAAATAACGTCCC tac tagagttgatagagggagaa | *Blautia* | **N** |  |

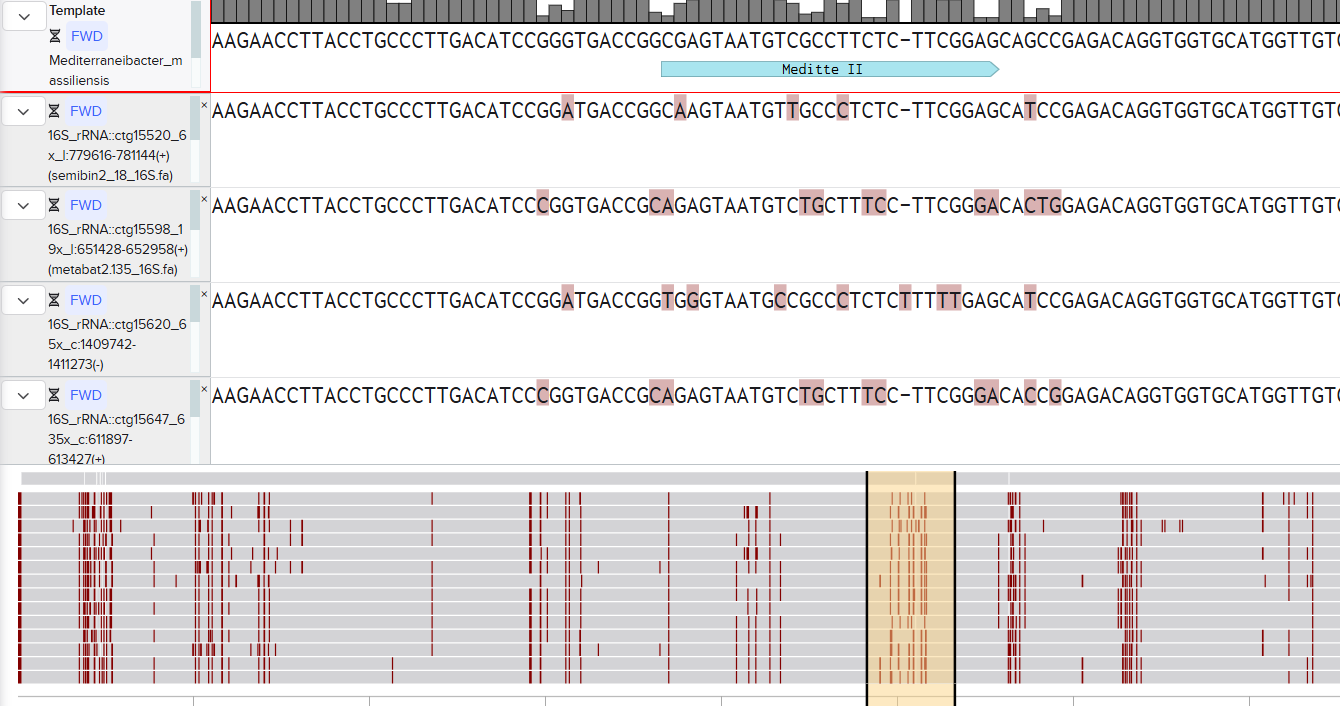

**Fig. S2.1: Screenshot of benchling.com alignment of publicly available 16S rRNA sequence of *Mediterraneibacter massiliensis* against *Mediterraneibacter* 16S rRNA sequences obtained from PacBio long-read sequencing.** Meditte II was a probe designed against the *M. massiliensis* sequence.

**Table S2.3: Probes designed and tested on full-length 16S rRNA sequences obtained via PacBio long-read sequencing.** In the SG column, "Y" indicates the presence of a detectable signal above background noise, while "N" denotes the absence of such a signal. In the SP column, "Y" confirms specific binding to the target genus, whereas "N" indicates non-specific signals. An example of this is shown in Supplementary Fig. 2**.**

| **No.** | **Probe Name** | **Sequence** | **Target genus** | **SG** | **SP** |
| --- | --- | --- | --- | --- | --- |
| BW156 | Cacco III 488 | TCGAAACCGCTTCGCTCGACTT ctt tgtggagggattgaaggata | *Caccovicinus* | **Y** | **N** |
| BW157 | Cacco IV 488 | CCGCGCCGTCTACGCTC ctc tgtggagggattgaaggata | *Caccovicinus* | **Y** | **Y** |
| BW158 | Lacto I 546 | GTTCCGCTCGCTCGACTTG tgg tgaaaggaatgggttgtggt | *Lactobacillus* | **Y** | **N** |
| BW159 | Lacto II 546 | TCCGCCGCTCGCTTTCC tcc tgaaaggaatgggttgtggt | *Lactobacillus* | **Y** | **Y** |
| BW160 | Lachno I 594 | CTCCAAATCGCTTCGCTCGACT act tagagttgatagagggagaa | *Lachnoclostridium* | **N** |  |
| BW161 | Lachno II 594 | CTTTGCCCACCAACACCTAG tag tagagttgatagagggagaa | *Lachnoclostridium* | **N** |  |
| BW162 | Strepto I 647 | AGGCTCGCCTGAGCAGG agg ttggaggtgtagggagtaaa | *Streptococcus* | **N** |  |
| BW163 | Strepto II 647 | GCTACAAGGCAGGTTACCTACG acg ttggaggtgtagggagtaaa | *Streptococcus* | **Y** | **Y** |
| BW164 | Mediter I 594 | GTATTTAGCCGGTGCTTCTTAG tag tagagttgatagagggagaa | *Mediterraneibacter* | **Y** | **N** |
| BW165 | Mediter II 594 | TCAAGGGCAGGTAAGGTTCTTCG tcg tagagttgatagagggagaa | *Mediterraneibacter* | **Y** | **Y** |
| BW166 | Gallim I 488 | GGTACAGTCACTTTCTTCGTCC tcc tgtggagggattgaaggata | *Gallimonas* | **Y** | **N** |
| BW167 | Gallim II 488 | ACCCCATGCTCCGCTGC tgc tgtggagggattgaaggata | *Gallimonas* | **Y** | **N** |
| BW168 | Pelet I 546 | AAAGGGAGCGTAGACGGA gga tgaaaggaatgggttgtggt | *Pelethomonas* | **Y** | **N** |
| BW169 | Pelet II 546 | ATCTTCCAGCGATAAATCTTTGGCA gca tgaaaggaatgggttgtggt | *Pelethomonas* | **Y** | **N** |
| BW170 | Blauti I 647 | GGCCGTCCACCCTCTCA tca ttggaggtgtagggagtaaa | *Blautia* | **Y** | **N** |
| BW171 | Blauti II 647 | CAGGCTCGCTTCCCTTT ttt ttggaggtgtagggagtaaa | *Blautia* | **Y** | **N** |
| BW172 | Chola I 594 | GTTCTCCCTGGCAGTCC tcc tagagttgatagagggagaa | *Choladocola* | **Y** | **N** |
| BW173 | Chola II 594 | CGGGCGTTGCCAACTCC tcc tagagttgatagagggagaa | *Choladocola* | **N** |  |
| BW174 | Agath I 488 | CCTCCTAAAGGCAGATTGCTCAC cac tgtggagggattgaaggata | *Agathobaculum* | **Y** | **Y** |
| BW175 | Agath II 488 | CCCCGGATTTCACTCCC tcc tgtggagggattgaaggata | *Agathobaculum* | **Y** | **N** |
| BW176 | Acuta I 647 | AGACGTTATCCCCCTCTGAAAG agg ttggaggtgtagggagtaaa | *Acutalibacter* | **Y** | **Y** |
| BW177 | Acuta II 647 | GCGAGCCCATCTTTCAGCG gcg ttggaggtgtagggagtaaa | *Acutalibacter* | **Y** | **N** |
| BW178 | Laws I 488 | AGGTACCGTCACTTGCTTCG tcg tgtggagggattgaaggata | *Lawsonibacter* | **Y** | **N** |
| BW179 | Laws II 488 | GTTTCAAATGCAGGCCACAGGTTG ttg tgtggagggattgaaggata | *Lawsonibacter* | **Y** | **N** |
| BW180 | Flavo I 594 | GGCCATCTCAGAGCGATAAATCTT ctt tagagttgatagagggagaa | *Flavonifractor* | **Y** | **N** |
| BW181 | Flavo II 594 | GTTACTGTCCAGCAATCCGC cgc tagagttgatagagggagaa | *Flavonifractor* | **Y** | **N** |
| BW182 | Lachno III PaBl | ACCGATGGGCTTTGCCC ccc AGAGTGAGTAGTAGTGGAGT | *Lachnoclostridium* | **N** |  |
| BW183 | Lachno IV PaBl | TTTTTGAGATTTGCTCCGGCTC ctc AGAGTGAGTAGTAGTGGAGT | *Lachnoclostridium* | **N** |  |
| BW196 | Lachno V 546 | GCCCACCAACACCTAGTATTC ttc tgaaaggaatgggttgtggt | *Lachnoclostridium* | **Y** | **Y** |
| BW197 | Blauti III 647 | TTGCTGCGGCACCgAAGAGC agc ttggaggtgtagggagtaaa | *Blautia* | **Y** | **N** |
| BW198 | Blauti IV 647 | CACCCCAGTCATCCGTC gtc ttggaggtgtagggagtaaa | *Blautia* | **Y** | **Y** |
| BW199 | Eintest III 488 | GCTTCACCGTCTACGCTC ctc tgtggagggattgaaggata | *Eisenbergiella* | **Y** | **N** |
| BW200 | Eintest IV 488 | CCTGCTGGCTACTAACCATAAGG agg tgtggagggattgaaggata | *Eisenbergiella* | **Y** | **N** |
| BW241 | Gallim III 488 | CGGGTGTAGCCCAGAACGTAAG aag tgtggagggattgaaggata | *Gallimonas* | **N** |  |
| BW242 | Gallim IV 488 | GCACCGAAGTTGAGCCCC tat tgtggagggattgaaggata | *Gallimonas* | **N** |  |
| BW243 | Pelet III 546 | GGCCGGCCAACCTCTCA tca tgaaaggaatgggttgtggt | *Pelethomonas* | **Y** | **Y** |
| BW244 | Pelet IV 546 | GGCACGCAGGGGGTCAG cag tgaaaggaatgggttgtggt | *Pelethomonas* | **N** |  |
| BW245 | Chola III 647 | GCCGGCTTATGCGGTATTAG tag ttggaggtgtagggagtaaa | *Choladocola* | **Y** | **Y** |
| BW246 | Chola IV 647 | TGTCTTTGACACCCGACACC acc ttggaggtgtagggagtaaa | *Choladocola* | **N** |  |
| BW247 | Laws III 594 | CGTTGTGTTATTCCCCACTC ctc tagagttgatagagggagaa | *Lawsonibacter* | **Y** | **Y** |
| BW248 | Laws IV 594 | TTGGCAGCCAGAGTCATGC tgc tagagttgatagagggagaa | *Lawsonibacter* | **Y** | **N** |
| BW249 | Flavo III 488 | TTAGCAACCCTTTCGGGCT gct tgtggagggattgaaggata | *Flavonifractor* | **N** |  |
| BW250 | Flavo IV 488 | CGAGGCCATCTCAGAGCGATAAATC atc tgtggagggattgaaggata | *Flavonifractor* | **Y** | **N** |
| BW253 | Gallim V 488 | CCCATCGTCGGTTTGGTG gtg tgtggagggattgaaggata | *Gallimonas* | **Y** | **N** |
| BW254 | Gallim VI 546 | GGTTAGAACACCGACTTTGGGTACT act tgaaaggaatgggttgtggt | *Gallimonas* | **Y** | **N** |

#### 3. Micro-scale spatial metagenomics (MSSM) method development

Optimising lysis and library preparation is crucial for accurate microbial profiling, as inefficient lysis can bias taxonomic representation, and suboptimal library preparation affects sequencing quality. We systematically tested and refined our workflow to minimise bias, improve data quality and reliability, and ultimately enhance microbiome analysis. Table S3.1 below summarises the lysis methods tested, while Table S3.2 provides an overview of the library preparation conditions used in the MSSM experimental comparisons.

**Table S3.1: Overview of chemical and enzymatic lysis methods used for cell lysis.** The different lysis approaches tested combined the use of different chemical agents (e.g., guanidine salts, detergents like Tween 20 and Triton X-100) with enzymatic components (Lysobac and Protease/Proteinase K). Each method is designated by a code (L01-L08). Abbreviations: guanidine hydrochloride (GuHCl); tris-hcl hydrochloride ph8 (TrisHCl), ethylenediaminetetraacetic acid pH 8.0 (EDTA), polyethylene glycol sorbitan monolaurate (Tween 20), guanidinium isothiocyanate (GITC) and n-lauroylsarcosine sodium salt solution (Sarkosyl). The chemical lysis buffer formulations used are based on standard commercial formulations or protocols reported in the scientific literature. References: 1) Lysis utilised in Winick-Ng *et al.**(4)*, 2) Buffer G2 (Qiagen, Cat. No.: 1014636), 3) Paul V. Haydock and Lynn M. Barker (2012, Patent No. US20130122496A1), 4) Guanidine Isothiocyanate Solution (Invitrogen, Cat. No.: 15577-018), 5) lysis buffer utilised in Lauritsen et al. (5) at 0.20x concentration 6) Composition of STET Lysis solution and 7) “Guidelines for Use” of Lysobac™ (Recombinant Human Lysozyme, InVitria) <https://invitria.com/resources/lysobac-guidelines-for-use/>

| **Code** | **Lysis Approach: Chemical (C) and Enzymatic (E)** | **Ref** |
| --- | --- | --- |
| **L01** | **C:** 800 mM GuHCl; 30 mM Tris•HCl; 2 mM EDTA; 5% Tween 20; 0.5% Triton X-100  **E:** Lysobac (0.1 mg/mL); Protease (2.116 units/mL) | 1 |
| **L02** | **C:** 800 mM GuHCl; 30 mM Tris•HCl; 30 mM EDTA; 5% Tween 20; 0.5% Triton X-100  **E:** Lysobac (0.1 mg/mL); Proteinase K (5.58 mg/mL) | 2 |
| **L03** | **C:** 1M GITC; 12.5 mM Tris•HCl; 5 mM EDTA; 2.5 % Tween 20; 0.25% Triton X-100  **E:** Lysobac (0.1 mg/mL); Proteinase K (5.58 mg/mL) | 3 |
| **L04** | **C:** 1M GITC; 12.5 mM Tris•HCl; 6.25 mM EDTA  **E:** Lysobac (0.1 mg/mL); Proteinase K (5.58 mg/mL) | 4 |
| **L05** | **C:** 1M GITC; 2,5 mM Citrate Buffer; 0.25% Sarkosyl  **E:** Lysobac (0.1 mg/mL); Proteinase K (5.58 mg/mL) | 5 |
| **L06** | **C:** 10 mM Tris•HCl; 0.1 M NaCl; 1 mM EDTA; 5% Triton X-100  **E:** Lysobac (0.1 mg/mL); Proteinase K (5.58 mg/mL) | 6 |
| **L07** | **C:** 100 mM Tris•HCl; 2 mM EDTA; 0.05% Triton X-100  **E:** Lysobac (0.1 mg/mL); Proteinase K (5.58 mg/mL) | 7 |
| **L08** | No chemical lysis (EB buffer)  **E:** Lysobac (0.1 mg/mL); Proteinase K (5.58 mg/mL) |  |

**Table S3.2: Summary of lysis and library preparation conditions used across experimental comparisons.** Lysis refers to the lysis method applied, coded according to Table S3.1. Reaction Volume specifies whether library preparation was conducted using half (50%) of the standard reagent volume (100%). Indexing Strategy indicates whether double indexing (DB) or single indexing (SG) was used, along with the corresponding products catalogue numbers. Method denotes whether library preparation was carried out manually (MN) or with the Tecan Fluent automation system (AS).

| **Comparison** | **Lysis** | **Reaction**  **Volume**  **(100 vs 50%)** | **Indexing**  **Strategy**  **(DB vs SG)** | **Method**  **(MN vs AS)** |
| --- | --- | --- | --- | --- |
| **Lysis mock community (S3.2)** | L01-L08 | 100 | DB (S02215-FG) | MN |
| **Lysis microsamples (S3.2) and Performance across intestinal sections (S3.3)** | L06 & L07 | 100 | DB (S02215-FG) | MN |
| **Resource optimisation: reaction volume (S3.5)** | L07 | 100 & 50 | DB (30219997) | MN |
| **Design considerations: LMD Size** | L06 | 50 | SG (S02366A) | MN |
| **Resource optimisation: automatisation (S3.5)** | L06 | 50 | SG (S02366A) | MN vs AS |
| **Discriminative power and replicability** | L06 | 50 | DB (30219997-30220000) | MN |

##### 1. Impact of lysis conditions on library preparation

Due to the inherently low DNA content in samples collected via laser micro-dissection (LMD), conventional extraction methods are unsuitable, necessitating the development of a streamlined workflow with minimal purification steps to minimise material loss. However, the direct use of crude lysates can hinder reaction efficiency and compromise library quality. To assess the extent of potential inhibitory effects, we compared two lysis methods, one containing chaotropic salts (L01) and one without (L06), on the Ovation® Ultralow V2 DNA-Seq Library Preparation Kit reactions. L01 exhibited significant inhibition, but introducing a magnetic bead-based clean-up (two systems tested: AMPure XP and HighPrep™ PCR beads) before library preparation led to a substantial increase in library yields, with an average molarity increase of 37 ± 27-fold (Table S3.1.1). In contrast, the inhibitory effect of L06 was less pronounced, resulting in a 2 ± 2-fold increase (Table S3.1.1).

Despite the potential loss of initial DNA yield, including a beads clean-up step ensured sufficient yield for sequencing at the tested DNA concentration for most of the reactions and was integrated into the MSSM workflow for further lysis comparisons.

**Table S3.1.1: Comparison of DNA library yields across lysis methods and clean-up conditions.** Indexed sequencing libraries were PCR-amplified using the number of cycles indicated. Final molarity (nmol/L) was quantified using the High Sensitivity NGS Fragment Kit (Agilent, Cat. No. DNF-474-0500) on a 5300 Fragment Analyzer (Agilent). Results are shown for three clean-up conditions: no clean-up (control), AMPure XP (Beckman Coulter) bead-based clean-up, and HighPrep PCR (MagBio) bead-based clean-up, across three biological replicates for lysis method L01 and L06.

| **Lysis** | **Replicate** | **Control** | | **AMPure XP** | | **HighPrep PCR** | |
| --- | --- | --- | --- | --- | --- | --- | --- |
|  |  | **Cycles** | **nmol/L** | **Cycles** | **nmol/L** | **Cycles** | **nmol/L** |
| **L01** | 1 | 20 | 2.3 | 19 | 161.5 | 19 | 46.2 |
|  | 2 | 20 | 6.6 | 19 | 110.7 | 19 | 93.6 |
|  | 3 | 17 | 0.7 | 17 | 53.1 | 17 | 19.1 |
| **L06** | 1 | 17 | 39.6 | 20 | 181.6 | 17 | 57.3 |
|  | 2 | 17 | 54.5 | 19 | 85.2 | 17 | 36.6 |
|  | 3 | 17 | 52.4 | 15 | 15.2 | 17 | 35.2 |

##### 2. Impact of lysis conditions on microbial taxonomic profiling

We initially evaluated eight different lysis methods (Table S3.1) using a mock community composed of four bacterial species (Table S3.2.1), standardised by equal cell numbers and comprising two Gram-positive (*Microbacterium oxydans* and *Paenibacillus amylolyticus*) and two Gram-negative (*Stenotrophomonas rhizophila* and *Xanthomonas retroflexus*). Some lysis methods were found to significantly inhibit the Ovation® Ultralow V2 DNA-Seq Library Preparation Kit (as shown in Table S3.1.1). To account for this in the comparison, DNA fragmentation was performed on diluted crude lysates, and a bead-based clean-up step was incorporated into the MSSM workflow prior to library preparation.

Community profiling results indicated that all eight methods successfully identified the four bacterial species without false positives when a genome-coverage cutoff of 30% was applied. Among these two methods, L06 and L07, containing Triton X-100, Lysobac, and Proteinase K, produced the most accurate composition statistics. Their observed abundance scores (Fig. S3.2.1a, b) closely matched the expected values, each species at 25% relative abundance (i.e. CLR = 0), with a root mean square error (RMSE) of approximately 0.2 (Fig. S3.2.1c). In contrast, methods incorporating other common lysis agents such as guanidine hydrochloride and guanidinium isothiocyanate showed RMSE values exceeding 0.35. Based on these results, L06 and L07 were selected for further evaluation on intestinal sections. The analysis revealed that enzymatic lysis alone (L08, Fig. S3.2.1c) generally underperformed compared to approaches combining enzymatic and chemical agents (RMSE = 0.52). Therefore, we recommend a combined enzymatic-chemical lysis strategy for improved microbial community profiling accuracy, although potential inhibitory effects should be taken into account.

**Table S3.2.1: Composition of the mock community.** Taxonomic Identities, Reference Genomes, and Assembly Metrics (Genome Length and Number of Contigs).

| **Bacterial isolate** | **Reference genome** | **Genome length** | **# Contigs** |
| --- | --- | --- | --- |
| ***Stenotrophomonas rhizophila*** | DSM14405 | 4,648 Mb | 1 |
| ***Xanthomonas retroflexus*** | ASM900143117v1 | 4,641 Mb | 32 |
| ***Microbacterium oxydans*** | NBRC 15586 | 3,892 Mb | 14 |
| ***Paenibacillus amylolyticus*** | FSL H7-0692 | 7,030 Mb | 36 |

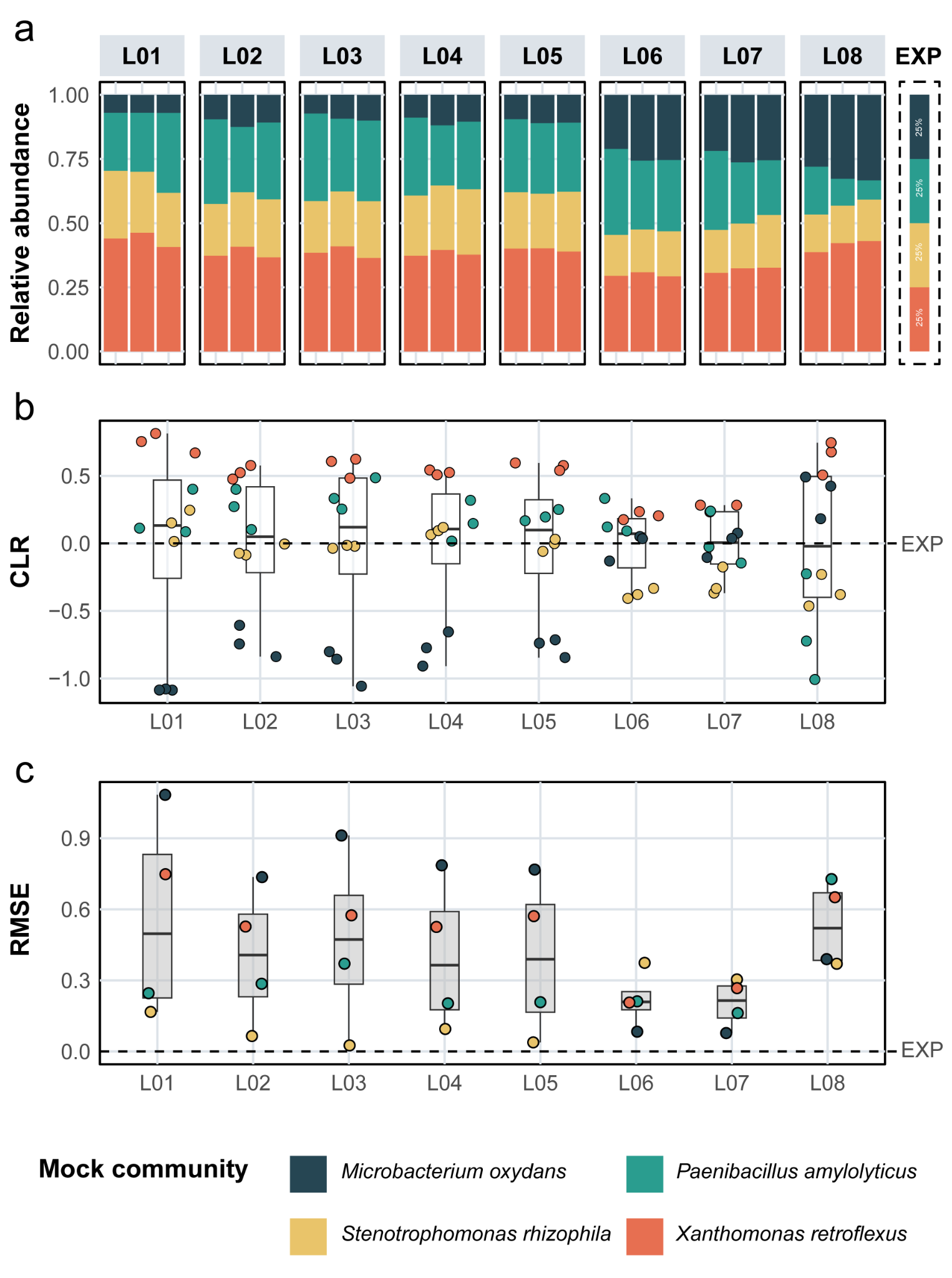

**Fig. S3.2.1: Taxonomic profiling of the mock community across lysis methods. a,** All lysis methods (L01-L08) recovered reads from all four bacterial species, though with varying relative abundances. Conditions L06 and L07 produced profiles closest to the expected equal distribution of 25% for each species. **b,** CLR-transformed relative abundance of the four species for each lysis method. The dashed line indicates the expected CLR value of the mock community. **c,** Root Mean Square Error (RMSE) comparing observed to expected bacterial abundances in the mock community. Methods L06 and L07 showed the best performance with RMSE values around 0.2, whereas other buffers exhibited deviations above 0.35.

No clear performance advantage was observed between methods L06 and L07 in caecum microsamples. L07 produced significantly higher sequencing yields post-quality filtering (GLM_quasipoisson_: *F*_(1, 30)_=6.80, *p*=0.014, Fig. S3.2.2a), and a greater proportion of reads mapping to reference bacterial genomes after removing host and contaminant reads (GLM_quasibinomial_: *F*_(1, 30)_= 4.84, *p*=0.036, Fig. S3.2.2b). However, both methods exhibited similar duplication rates (GLM_quasibinomial_: *F*_(1, 30)_=0.43, *p*=0.519, Fig. S3.2.2c) and no significant differences in richness or neutral diversity (*p*>0.05, Fig. S3.2.2d-e-f). Neither lysis exclusively identified bacterial genera. Although not statistically significant, L06 yielded marginally higher average neutral diversity, a metric integrating both richness and evenness, and was therefore selected for subsequent experiments in both the caecum and colon (Fig. S3.2.2e).

Although L06 was initially selected, a subsequent experiment suggested that L07 may offer enhanced lysis performance in colon microsamples. The use of L07 improved colon results, resulting in increased sequencing yield post-filtering (GLM_quasipoisson_: *F*_(1, 31)_=23.23, *p*<0.001; Fig. S3.2.2a) and higher alpha diversity across all indices including richness (GLM_quasipoisson_: *F*_(1, 31)_=12.99, *p*=0.001; Fig. S3.2.2d) and neutral diversity (LM: *F*_(1, 31)_=4.40, *p*=0.044; Fig. S3.2.2e). Notably, L07 enabled the detection of 10 additional genera (13.7%), whereas L06 revealed only a single unique genus.

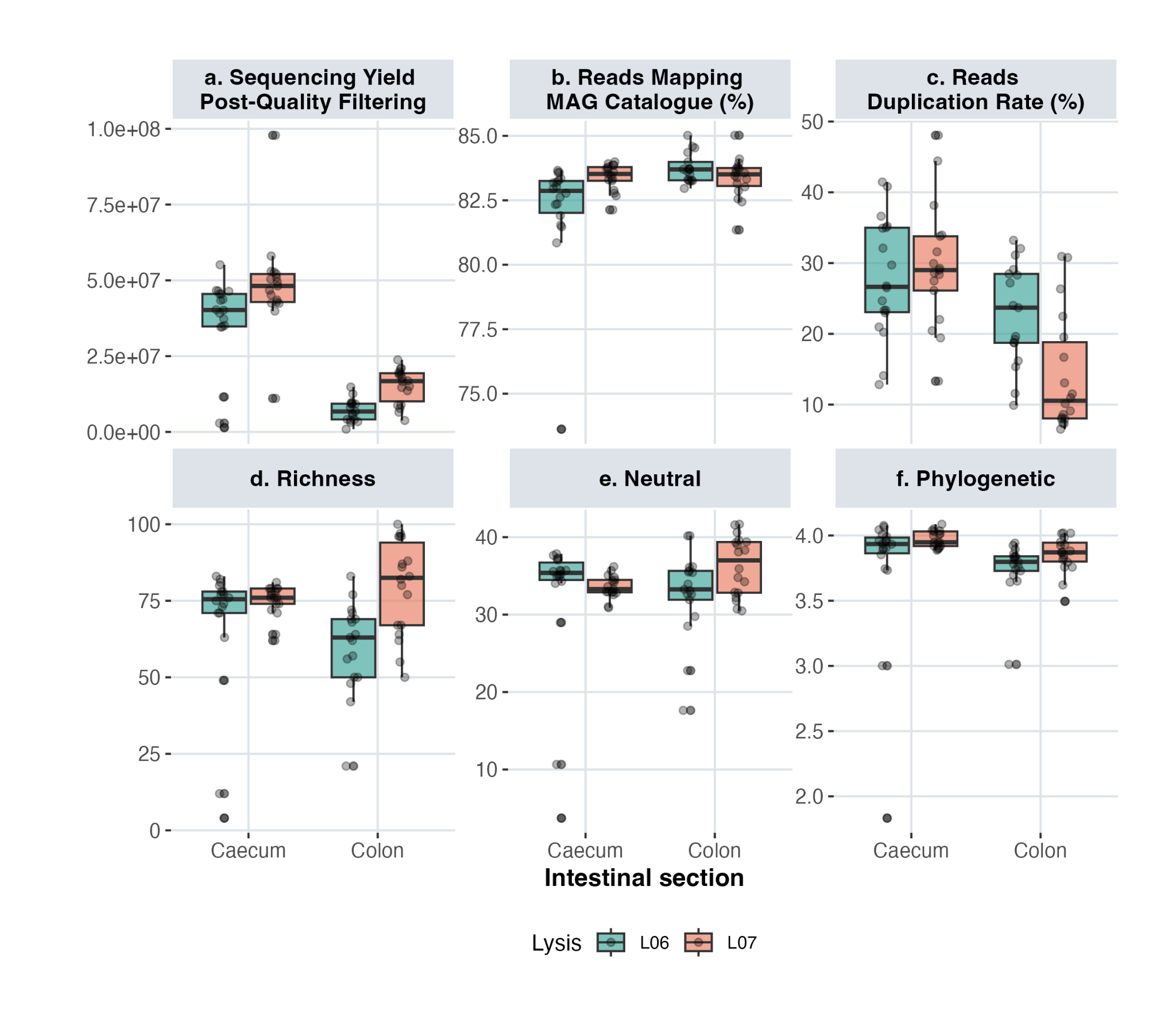
**Fig. S3.2.2: Comparison of sequencing and alpha diversity metrics across lysis methods (L06 and L07) in two intestinal sections (caecum and colon). a,** Sequencing yield measured by the number of reads retained after quality filtering and adapter trimming. **b,** Mapping success, defined as the percentage of quality-filtered reads mapped to the reference genome catalogue. **c,** Data complexity assessed by the percentage of duplicated raw reads. **d,** Species richness derived from each lysis method. **e,** Neutral diversity (exponential Shannon index) observed across methods. **f,** Phylogenetic diversity metrics yielded by the 2 lysis methods.

##### 3. Performance evaluation across intestinal sections

MSSM performance was evaluated using 5,000 μm² microsamples from three sections of the digestive tract (ileum, caecum, and colon) of a 35-day-old animal. After applying a stringent 30% genome coverage cutoff to minimise false identifications caused by cross-mapping of reads between related taxa, the number of samples with retained reads after filtering (>50,000) remained high: 97.2% for the caecum and colon, and 88.8% for the ileum. However, post-filtering yields were highest in the caecum, followed by the colon and then the ileum (GLM_quasipoisson_: *F*_(2, 96)_=60.73, *p*<0.001, Fig. S3.3.1a), despite sequencing all microsamples to a target depth of ~4 GB. Colon and ileum samples had a higher proportion of low-quality reads, with estimates at 51% (95% CI: 40-62) and 35% (95% CI: 25-48) respectively, primarily due to adapter contamination (GLM_quasibinomial_: *F*_(2, 96)_=25.85, *p*<0.001, Fig. S3.3.1b). Host-derived read contamination was minimal in caecum and colon samples but more pronounced in the ileum, where host reads were estimated at 15% (95% CI: 9-23).

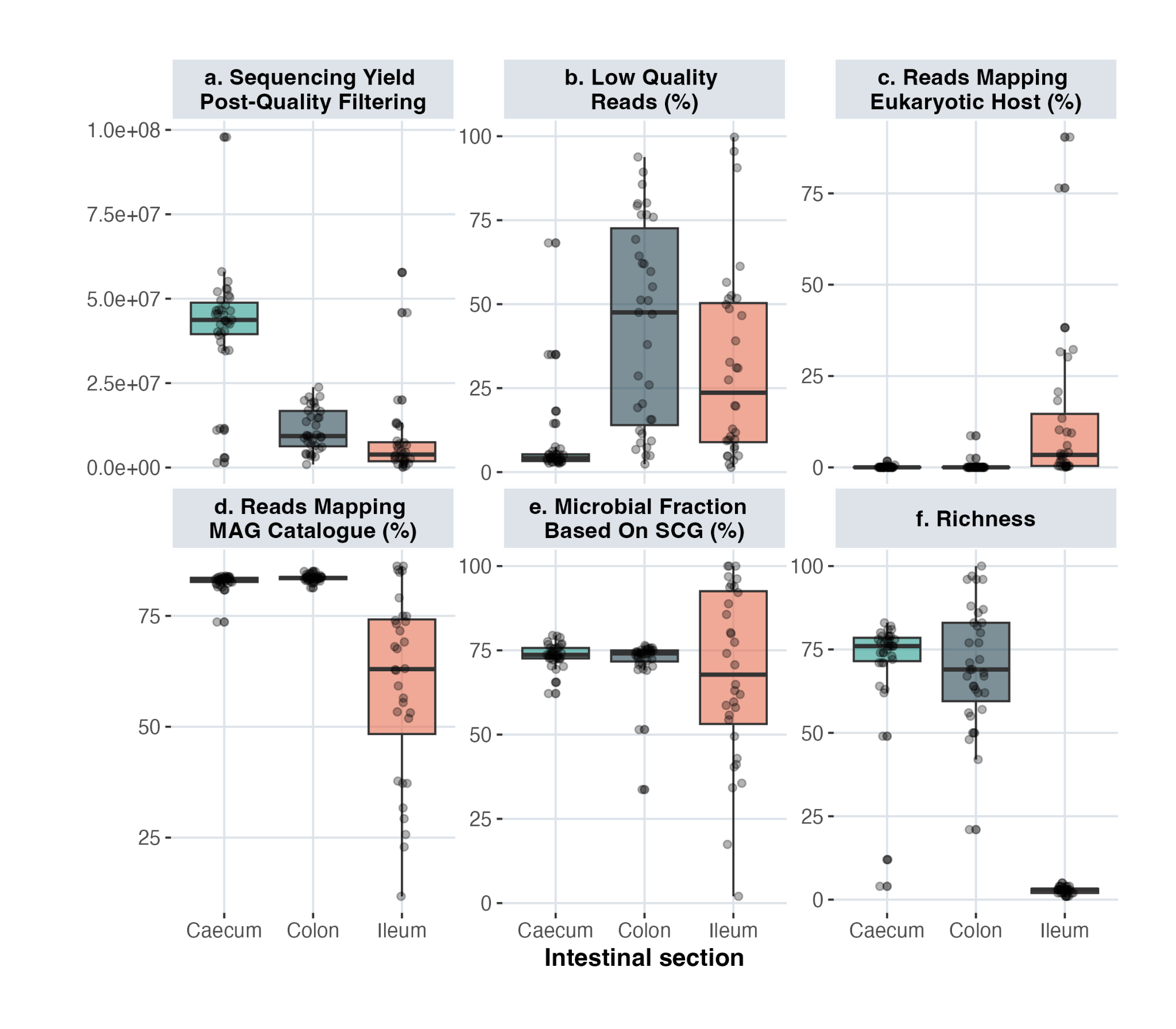
**Fig. S3.3.1: Comparison of micro-scale spatial metagenomics (MSSM) performance across three intestinal sections (caecum, colon, and ileum). a,** Sequencing yield measured by the number of reads retained after quality filtering and adapter trimming. **b,** Sequencing quality measured by percentage of reads discarded due to low quality. **c,** Host interference measured as percentage of reads mapped to the reference chicken genome. **d,** Prokaryotic fraction estimation based on mapping reads to the reference genome catalogue and evaluation of mapping efficiency. **e,** Estimation of the microbial fraction based on mapping against a curated set of single-copy genes (SingleM). **f,** Species richness derived from three intestinal sections.

Despite lower sequencing performance in the colon, post-filtering mapping percentages were comparable between the caecum and colon, with estimated values of 84% (95% CI: 81-87 and 80-87, respectively; Fig. S3.3.1d). In contrast, the ileum exhibited significantly lower and more variable estimates of mapping percentages across samples (62%, 95% CI: 56-67, GLM_quasibinomial_: *F*_(2, 96)_=50.32, *p*<0.001). This variability may arise from the absence of relevant bacterial taxa in the reference catalogue, which was constructed exclusively from caecal content and may not adequately represent the microbial composition of ileal samples. Supporting this interpretation, independent estimates of the microbial fraction based on single-copy gene analysis (Fig. S3.3.1e) showed a moderate correlation with the proportion of reads mapping to the reference post-filtering (Spearman’s *ρ*=0.66). Although microbial fractions in ileum samples tended to be lower and more variable, the difference was not statistically significant, suggesting that both limited catalogue representation and a potentially reduced microbial load contribute to the observed lower performance. These limitations were further reflected in the ileum’s significantly reduced species richness (Fig. S3.3.1f), where only a limited bacterial community was detected, comprising just six species, predominantly *Lactobacillus* spp. Consequently, no additional sequencing was conducted for the ileum, as the combination of low mapping efficiency and likely low biomass precluded reliable characterisation of its microbial community.

Microbial characterization showed that species richness was significantly higher in the caecum and colon than in the ileum, reflecting a more diverse bacterial community in these sections. Nevertheless, variability in MSSM performance between the caecum and colon likely arises from differences in microbial biomass, DNA quality, and technical factors. Efforts to compensate for low biomass in the colon by increasing PCR cycles did not improve the quality of colon results. Increasing the number of PCR cycles from 15 to 19 led to higher read counts before quality filtering (GLM_quasipoisson_: *F*_(1, 61)_=94.32, *p*<0.001). However, reactions with 15 cycles yielded significantly more reads after filtering (GLM_quasipoisson_: *F*_(1, 61)_=4.91, *p*=0.03). This was due to an increase in low-quality reads (GLM_quasibinomial_: *F*_(1, 61)_=31.21, *p*<0.001). Moreover, duplication rates were also significantly increased in the 19-cycle reactions (GLM_quasibinomial_: *F*_(1, 61)_=39.81, *p*<0.001). In the 35-day-old animal, the bacterial communities of the caecum and colon were dominated by the same four orders: *Lachnospirales*, *Oscillospirales*, and *Lactobacillales*, with minor contributions from *Christensenellales* and *Erysipelotrichales* (Fig. S3.3.2). After excluding genera present in fewer than 5% of samples, the two sections shared all genera. A section comparison for the 14-day-old animal was not possible due to poor data quality of the colon dataset. A substantial proportion of reads mapped to the host genome (median 32.86%, IQR = 2.7-70.4%), and none of the 72 sequencing reactions passed the stringent 30% genome coverage cutoff.

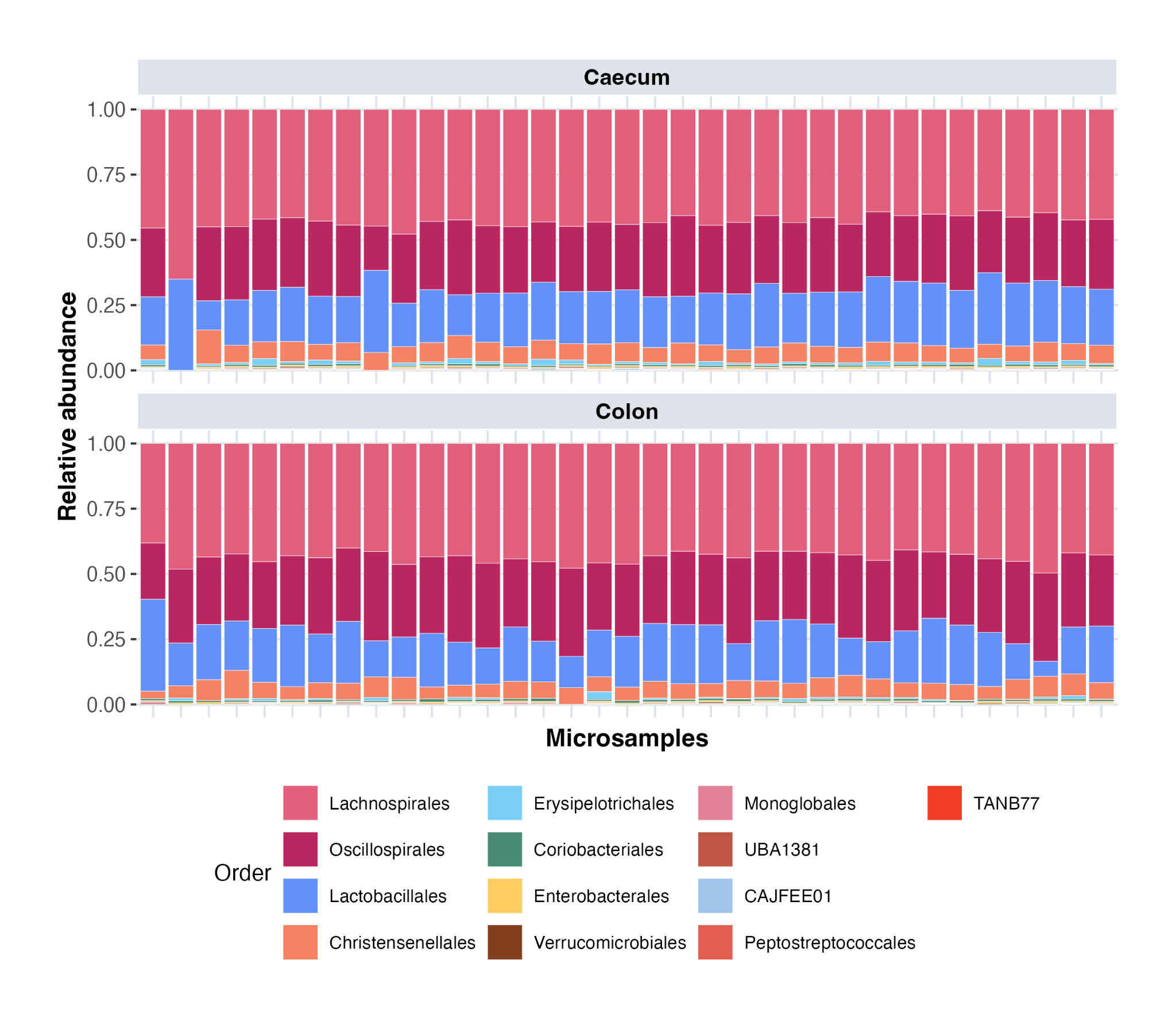

**Fig. S3.3.2: Bacterial community composition (at the order level) of 5,000 μm² microsamples obtained from the caecum and colon.**

To directly assess whether sequencing differences between intestinal sections reflected true variations in microbial biomass or technical variability, confocal microscopy of DAPI-stained microsections from the 35- and 14-day-old animals used for MSSM was performed. This analysis provided direct visual confirmation that differences in bacterial density across intestinal regions (caecum, colon, and ileum) were the primary factor influencing MSSM performance, rather than MSSM technical variability. The images revealed clear visual contrasts between sections (Fig. S3.3.3a). In the 35-day-old animal, quantification of bacterial cells within a 5,000 μm² area using QuPath revealed significantly higher mean bacterial densities in the caecum (648 cells/5,000 μm², IQR = 402-1,112) and colon (535 cells/5,000 μm², IQR = 293-889) relative to the ileum, which exhibited an extremely low count. Furthermore, DAPI staining showed little to no bacterial presence in the colon of the 14-day-old animal, aligning with MSSM’s failure to generate high-quality data from these sections (Fig. S3.3.3a). Through bright-field imaging with the LMD scope and DAPI-staining the same cryosection, we also demonstrated that the noise in our quantification (failed sequencing reactions) may arise primarily from embedding and sectioning artifacts (Fig. S3.3.3b).

Furthermore, the observed MSSM performance closely reflects the recognised biological characteristics of the three intestinal regions. The ileum generally harbors a lower bacterial load(6), consistent with its role as a transitional intestinal segment with high absorption capacity and limited fermentative activity. In contrast, the caeca exhibit the highest bacterial biomass(6)^,^^(7)^ and diversity, serving as the primary site of fermentation in the avian gastrointestinal tract⁴(8). The colon typically contains a moderate bacterial load, influenced by the periodic release of contents from the caecum into the large intestine(9)^,^^(10)^. Despite this, an important limitation of the MSSM approach when interpreting these results is the challenge of sample collection. Although we minimised disturbance to the integrity of the intestinal section by handling the segment at the cut edges, some disruption of the luminal contents may have occurred during sampling. This could have introduced spatial biases and further exacerbated low biomass issues, particularly in samples from the 14-day-old individual.

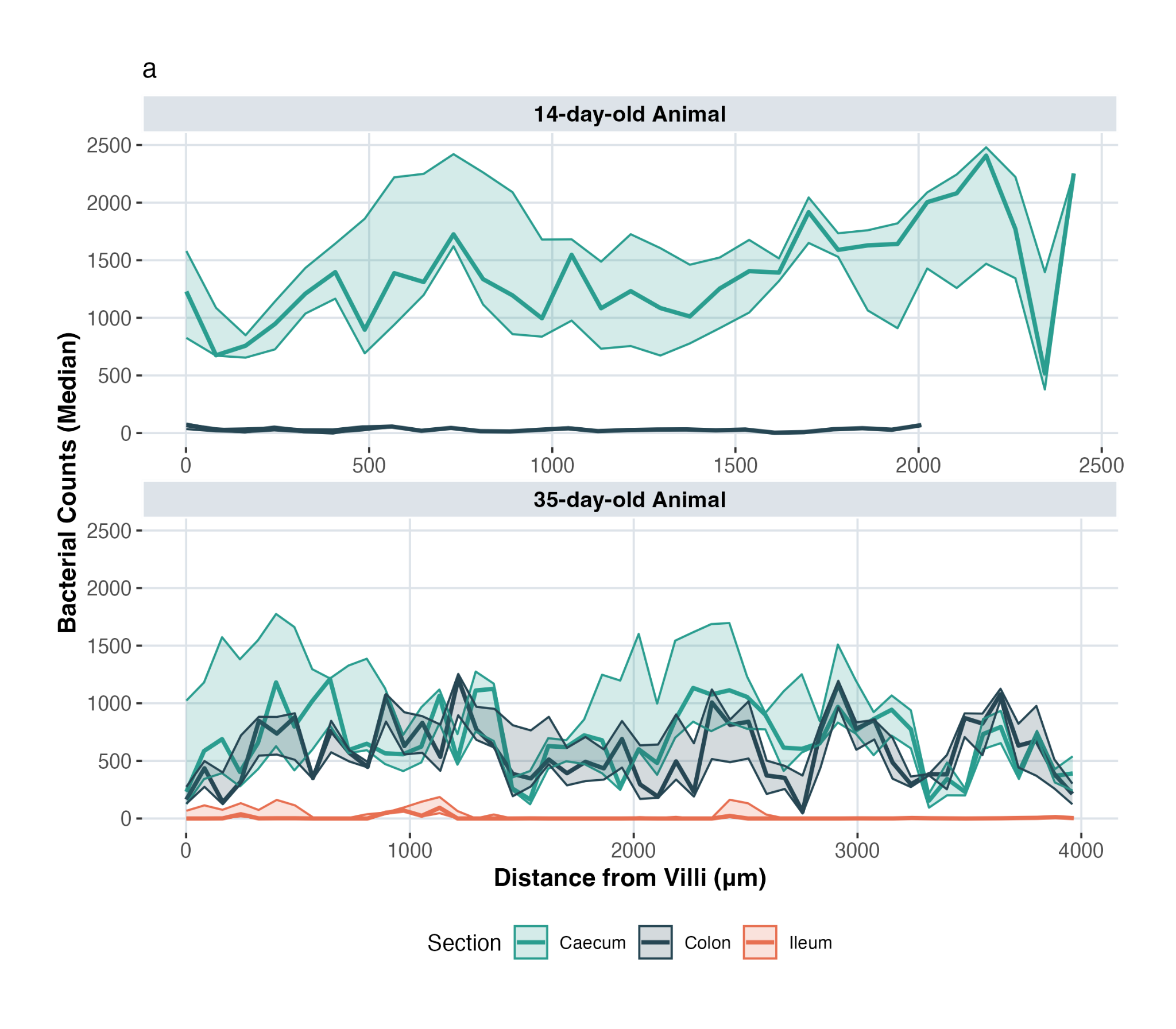

b

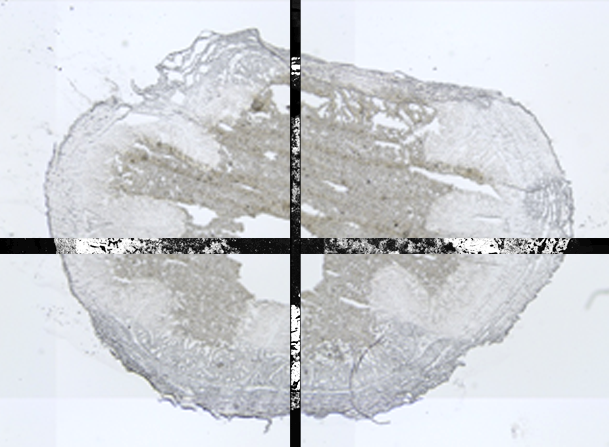

**Fig. S3.3.3: Quantification of bacterial biomass using confocal microscopy of DAPI-stained cryosections. a,** Estimated bacterial biomass (3 replicates) across intestinal sections in 35- and 14-day-old animals. The line indicates the median; shaded areas represent the interquartile range (IQR). Scale bars are 20 μm. **b,** Comparison of bright field versus confocal DAPI-stained images of the same cryosection to reveal impact of biological, embedding and sectioning artifacts.

##### 4. Design considerations: control reactions

With the exception of one sequencing batch (B10), negative samples consistently exhibited lower sequencing output compared to microsamples (Fig. S3.4.1a). This disparity was more evident in certain datasets (excluding B14 and B15) after quality filtering, as a substantial portion of reads in the negatives were discarded due to low quality (Fig. S3.4.1b). Additionally, these datasets exhibited high levels of adapter contamination in the negative samples (Fig. S3.4.1d). In contrast, B14 and B15, which were not affected by compromised quality, showed that a greater proportion of reads from negative samples mapped to the human genome (Fig. S3.4.1d). Despite discrepancies between datasets, the number of retained reads in negative samples remained minimal across all datasets (Fig. S3.4.1e), and mapping rates to the reference catalogue were consistently low (Fig. S3.4.1f).

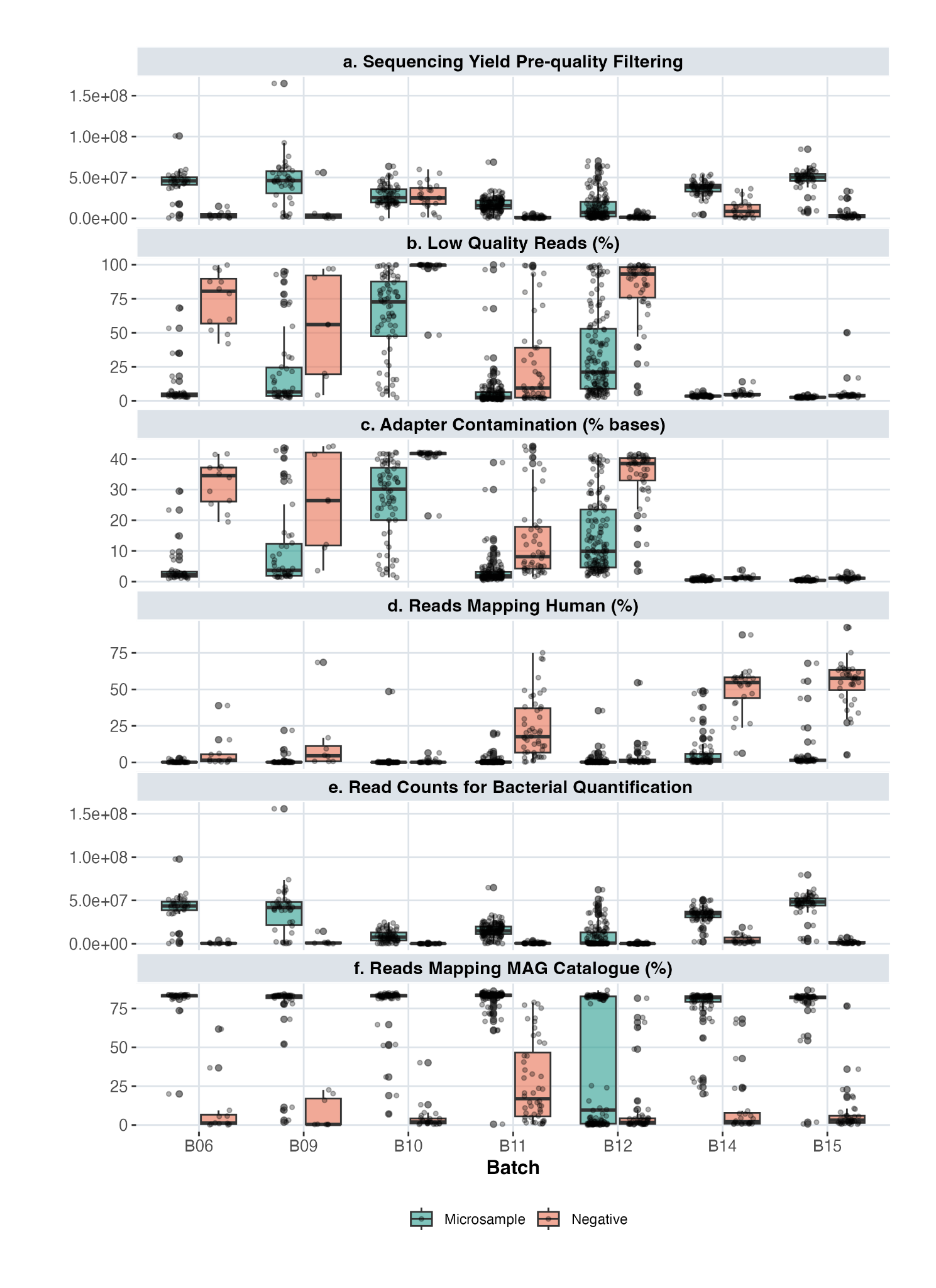
**Fig. S3.4.1: Comparison of sequencing performance and quality metrics across positive and negative control reactions. a,** Number reads obtained by sequencing. **b,** Sequencing quality measured by percentage of reads discarded due to low quality. **c,** Sequencing quality measured by percentage of bases trimmed as sequencing adapters. **d,** Percentage of reads mapped to the reference human genome due to contamination. **e,** Number reads retained after filtering for bacteria quantification. **f,** Mapping efficiency, represented by the percentage of reads mapped to the reference genome catalogue.

Without data filtering, PCA revealed a clear separation between most negative controls and microsample reactions (Fig. S3.4.2). Some microsamples, however, clustered with the negative controls; these were excluded after applying a 30% genome-coverage filtering, indicating low sequencing quality. Among the retained microsamples, separation along PC1 reflected variation between the two animals. In the unfiltered dataset of each animal, differential abundance analysis (ALDEx2) identified certain MAGs as significantly more abundant in negative controls compared to positives; however, those genomes were not retained after 30% coverage filtering, and thus the signal is likely attributable to cross-mapping artefacts.

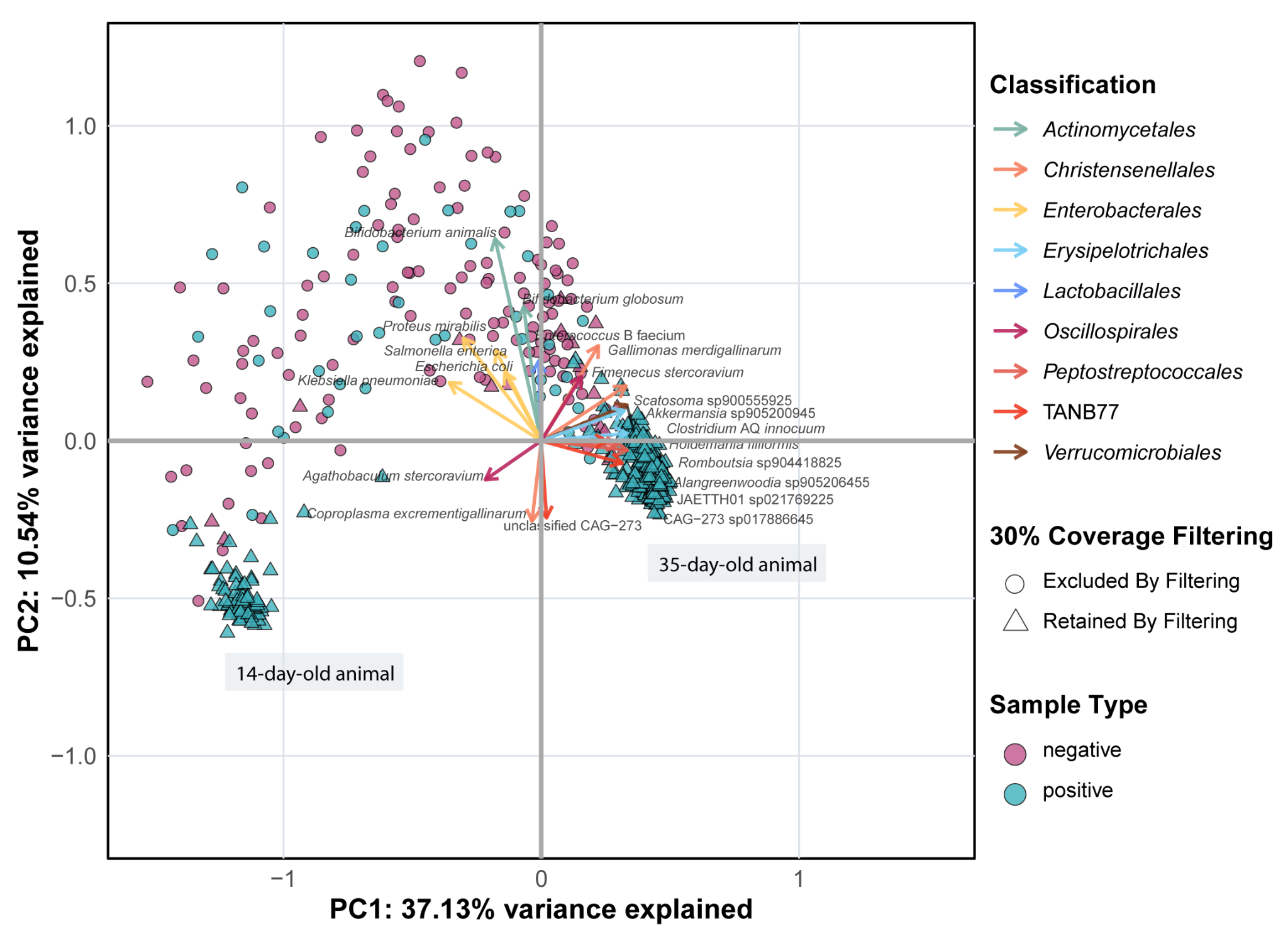

**Fig. S3.4.2: Principal Component Analysis (PCA) illustrates the variance in bacterial composition between control reactions and micro-scale samples.**

This finding aligns with mock community benchmarking studies demonstrating that genome coverage-based filtering is robust and accurate for eliminating false positives, more than traditional relative abundance filtering(11). A 5% genome coverage cutoff achieved a higher average F1 score, a composite metric of precision and recall, than any tested relative abundance criterion(11). Moreover, analysis of short-read metagenomic datasets demonstrated that applying a more stringent 30% genome coverage threshold maximised the F1 score, further supporting its efficacy in minimising false-positive taxa(11). In our MSSM datasets, applying the 30% threshold effectively excluded the majority of control-derived reactions (91%), along with low-quality microsample reactions (17.4%, primarily derived from colon of the 14-day-old animal), which did not meet the 30% genome coverage threshold, indicating insufficient genomic representation or potential artifactual signal (Fig. S3.4.3). While some degree of contamination is likely unavoidable in ultra-low biomass workflows such as MSSM, our results indicate that applying a stringent filtering strategy based on genome coverage can effectively account for such issues.

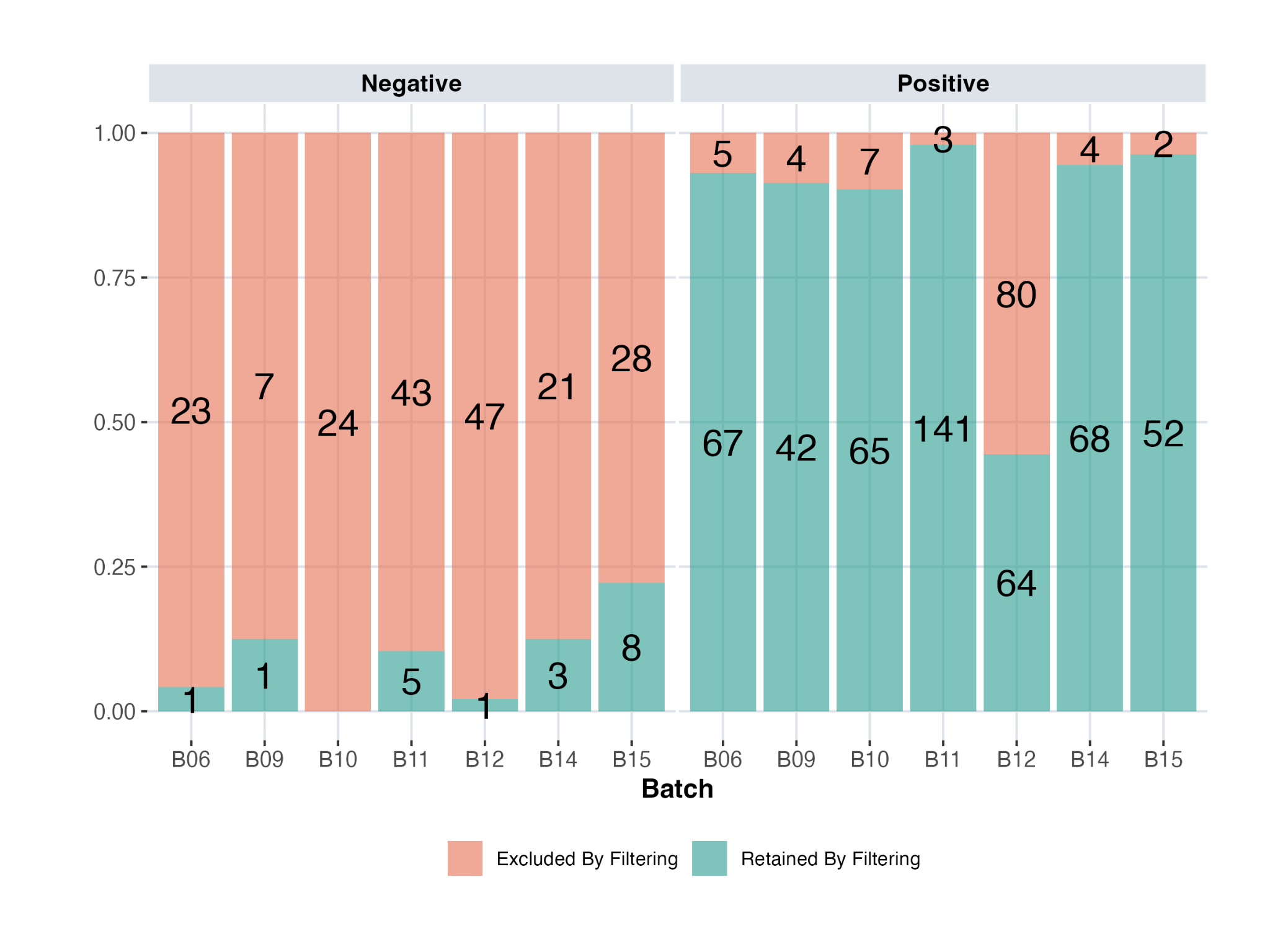

**Fig. S3.4.3: Ratio and absolute number (annotated inside bars) of positive and negative reactions excluded from each sequencing batch after applying the 30% genome coverage threshold.**

##### 5. Resource optimisation and throughput

We evaluated the use of reduced library reaction volumes (50%) across 14 reactions and found that microsample retention rates after quality filtering remained high (93%, with 13 out of 14 reactions successful). Although full-volume reactions yielded more raw sequencing reads (median of 54.46 x 10^6^ vs. 41.86 x 10^6^), post-quality filtering yields were comparable (median of 43.69 x 10^6^ vs. 40.40 x 10^6^). Half-volume reactions produced overall a higher-quality dataset. Only 3.3% of reads were classified as low-quality in the half-volume dataset, compared to 11.95% in the full-volume (Table S3.5.1). This difference was primarily driven by a few samples in full-volume reactions with elevated proportions of low-quality reads (Fig. S3.5.1a) mainly due to an increase in adapter contamination. The percentage of reads mapping to the reference MAG catalogue was comparable between half and full datasets (Table. S3.5.1, increase in the median mapping % by 8.05), indicating that reduced volumes do not impact microbial detectability. Despite a marginally higher fraction of unmapped reads in half-volume reactions (16.2% vs. 15.2%, Table. S3.5.1), the comparable read quality in half-volume reactions maximises the usable sequencing data, making them a more cost-effective and efficient alternative without compromising on performance.

**Table S3.5.1: Proportion of sequencing reads by category across resource optimisation comparisons (100% vs. 50% reaction volume and manual vs. automated library preparation).** Median values are reported for each comparison group to reflect read composition across microsamples. Categories include low quality, host-derived, contaminants-derived such as human and other eukaryotes, microbial reads mapping MAG catalogue, and reads that did not map.

| **Sequencing Reads** | **Cost Optimisation**  **Median %** | | **Throughput Optimisation**  **Median %** | |
| --- | --- | --- | --- | --- |
|  | **100% reaction** | **50% reaction** | **Automation**  **(1 plate set-up)** | **Manual** |
| **Low Quality** | 11.95 | 3.3 | 2.6 | 3.55 |
| **Host** | 0 | 0 | 0 | 0 |
| **Human** | 0 | 0 | 1.35 | 1.2 |
| **Other Eukaryotes** | 0 | 0 | 0 | 0 |
| **MAG catalogue** | 72.4 | 80.45 | 78.85 | 78.65 |
| **Unmapped** | 15.2 | 16.2 | 17 | 16.6 |

The validated 50% library reaction protocol was successfully integrated and customised on a Tecan Fluent automation system (DreamPrep 780), initially implemented as a single-plate setup (96 reactions) to increase processing throughput. As with the previous comparison, automating the library preparation did not affect microbial detectability, as measured by the percentage of reads mapping to the reference genome, with no notable difference observed between manual and automated workflows (Table S3.5.1, Fig. S3.5.1a). Read quality, assessed by the proportion of usable sequencing data remaining after the removal of low-quality reads and human contamination (Table S3.5.1), was marginally lower in the manual preparations dataset (95.25%) compared to the automated process (95.85%). The minor increase in the % of unmapped reads would not justify the continued use of manual processing, as the automated workflow is more time-efficient and offers a more sustainable use of resources (Table S3.5.2).

**Table S3.5.2: Processing time and estimated plastic consumption for manual versus automated implementation of the library preparation workflow.**

| **Method** | **Time** | **Plastic consumption per plate** |
| --- | --- | --- |
| **Manual (Single-Plate)** | 9h | 2980 g |
| **Fluent (Single-Plate)** | 7h | 2192 g |
| **Fluent (Dual-Plate)** | 9h | 2043 g |

To further increase throughput, the automated library preparation was expanded to a dual-plate setup (192 reactions), with interleaved processing steps that maximised efficiency and substantially boosted library preparation capacity. A comparison test between set-ups (single versus dual-plate) was carried out using a standard control (human DNA) at 0.1 ng/μL, with non-specific short fragments (between 100-200 bp) and library (between 200-1000 bp) molarity serving as metrics to evaluate library preparation efficiency. We found no strong evidence supporting a decrease in library preparation efficiency between the single- and dual-plate setups. One of the dual-plate preparations exhibited elevated levels of short fragments compared to both the single-plate preparation and the other plate within the dual-plate setup (Fig. S3.5.1b), suggesting some degree of plate-specific processing variation. However, the two plates in the dual-plate setup did not produce significantly lower library molarity relative to the single-plate preparations (Fig. S3.5.1b). Conversely, the data indicate that the dual-plate configuration may outperform the single-plate setup, given that one plate showed significantly superior performance.

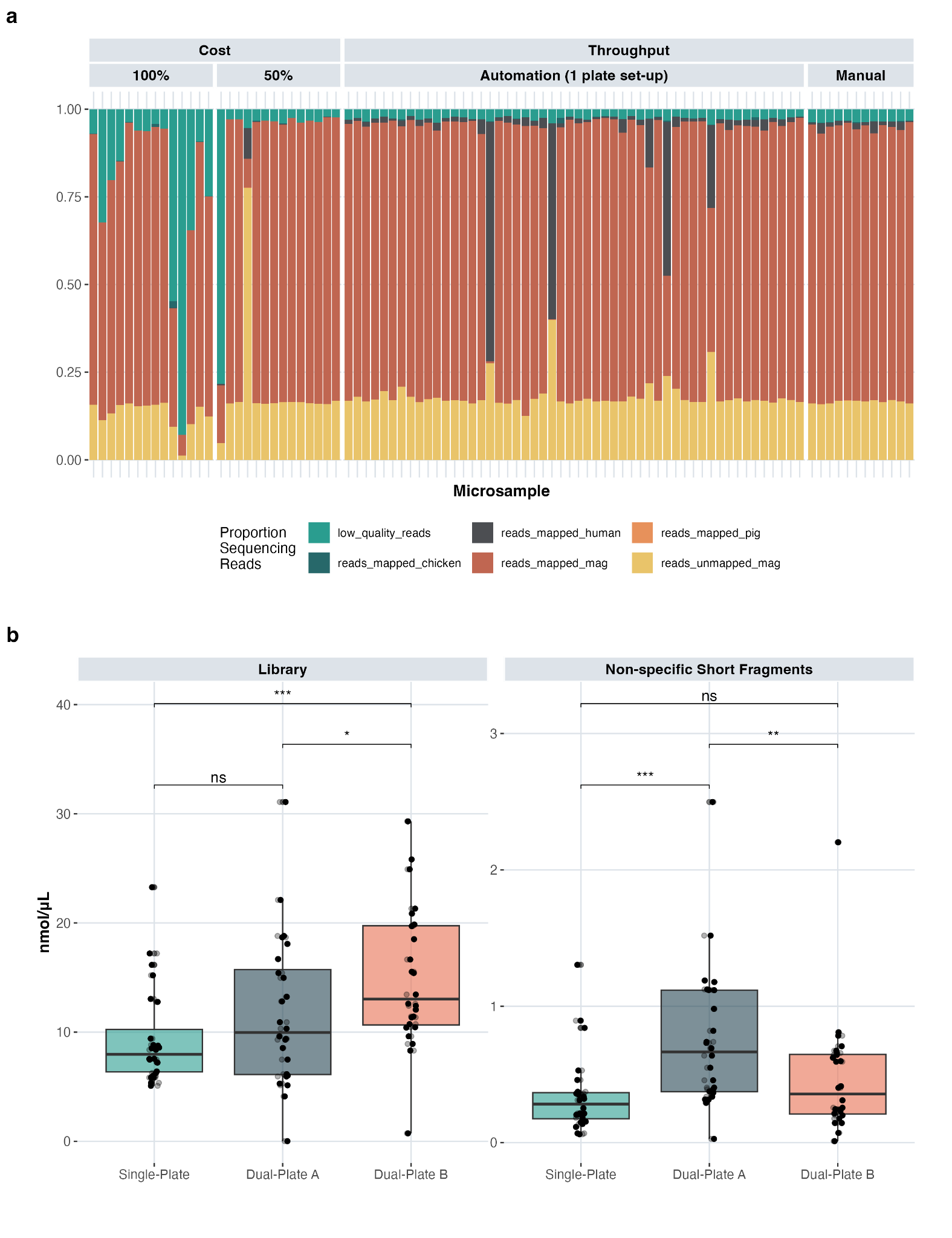

**Fig. S3.5.1: Evaluation of micro-scale spatial metagenomics (MSSM) resource optimisation. a,** Proportion of sequencing reads by type across microsamples. The stacked barplot displays the percentage of reads assigned to different categories (low quality, host-derived, contaminants-derived such as human and other eukaryotes, microbial reads mapping MAG catalogue and reads that did not map) for each comparison. Read proportions were calculated as a percentage of total sequencing reads per microsample. **b,** Comparison of non-specific short fragments (between 100-200 bp) and library (between 200-1000 bp) molarity, grouped by plate setup (single vs. dual A-B). Statistical significance between plates within each concentration was evaluated using Wilcoxon rank-sum tests with Benjamini-Hochberg correction for multiple comparisons. Significance levels are indicated by stars (**p<*0.05; ***p<*0.01; **p<*0.001 and “ns” for *p>*0.05) shown above relevant comparisons.

#### 4. Micro-scale Spatial Metagenomics (MSSM) method implementation

Additive partitioning of microbial diversity showed that the average microsample diversity (alpha component) observed in the caecum and the colon was lower than expected by chance. In contrast, the gain/increase in species richness observed when adding new microsamples within a cryosection (beta component) and the average cryosection diversity were higher than expected by chance (Fig. S4.1). This indicates that the microscale variability within the cryosections accounts for a large portion of the overall microbial diversity observed in the caecum and the colon of the chicken intestine.

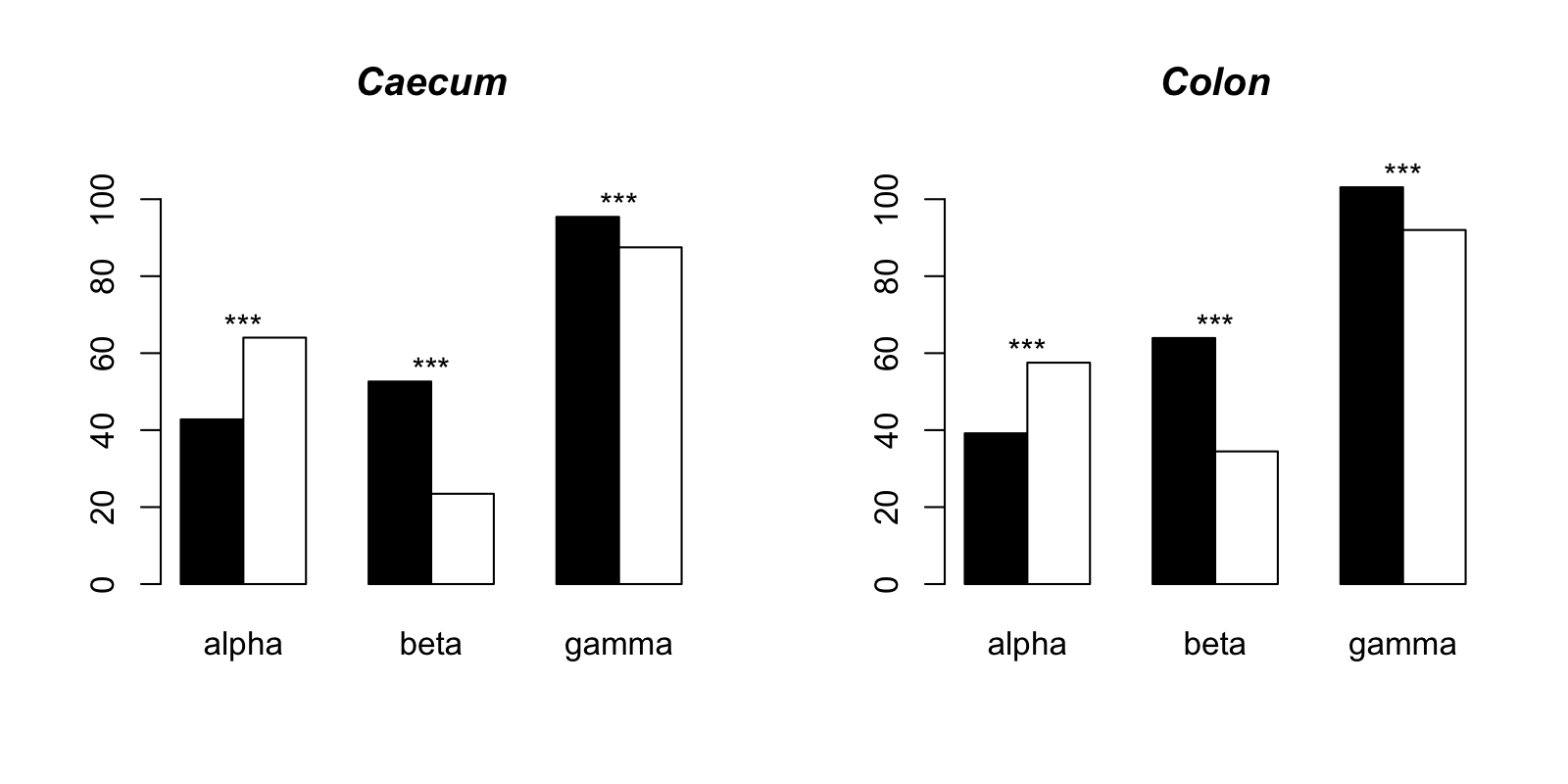

**Fig. S4.1: Additive partitioning of the overall microbial diversity into alpha (average microsample species richness), beta (average number of species when adding new microsamples within a cryosection) and gamma (average cryosection species richness) components in the caecum and the colon.** Black bars depict the observed values and the white bars the expected value under a null model. **p<*0.05; ***p<*0.01; ****p<*0.001.

The significant variability within a cryosection observed in the diversity partitioning for the colon was accompanied by a significant increase in Aitchison distance between microsample communities as the spatial distance between them widened (nperm=10000, *p<*0.001; Fig. S4.2a). As further evidence for fine-scale spatial structuring of microbial communities, the mantel correlogram analysis captured a significant spatial autocorrelation at the shortest spatial distances between microsamples (Fig. S4.2b). In contrast, neither the distance analysis (nperm=10000, *p=*0.110, Fig. S4.2c) nor the mantel correlogram (p>0.05, Fig. S4.2d) detected a significant spatial autocorrelation in the caecum cryosections, suggesting that the microscale spatial variability observed was not spatially structured.

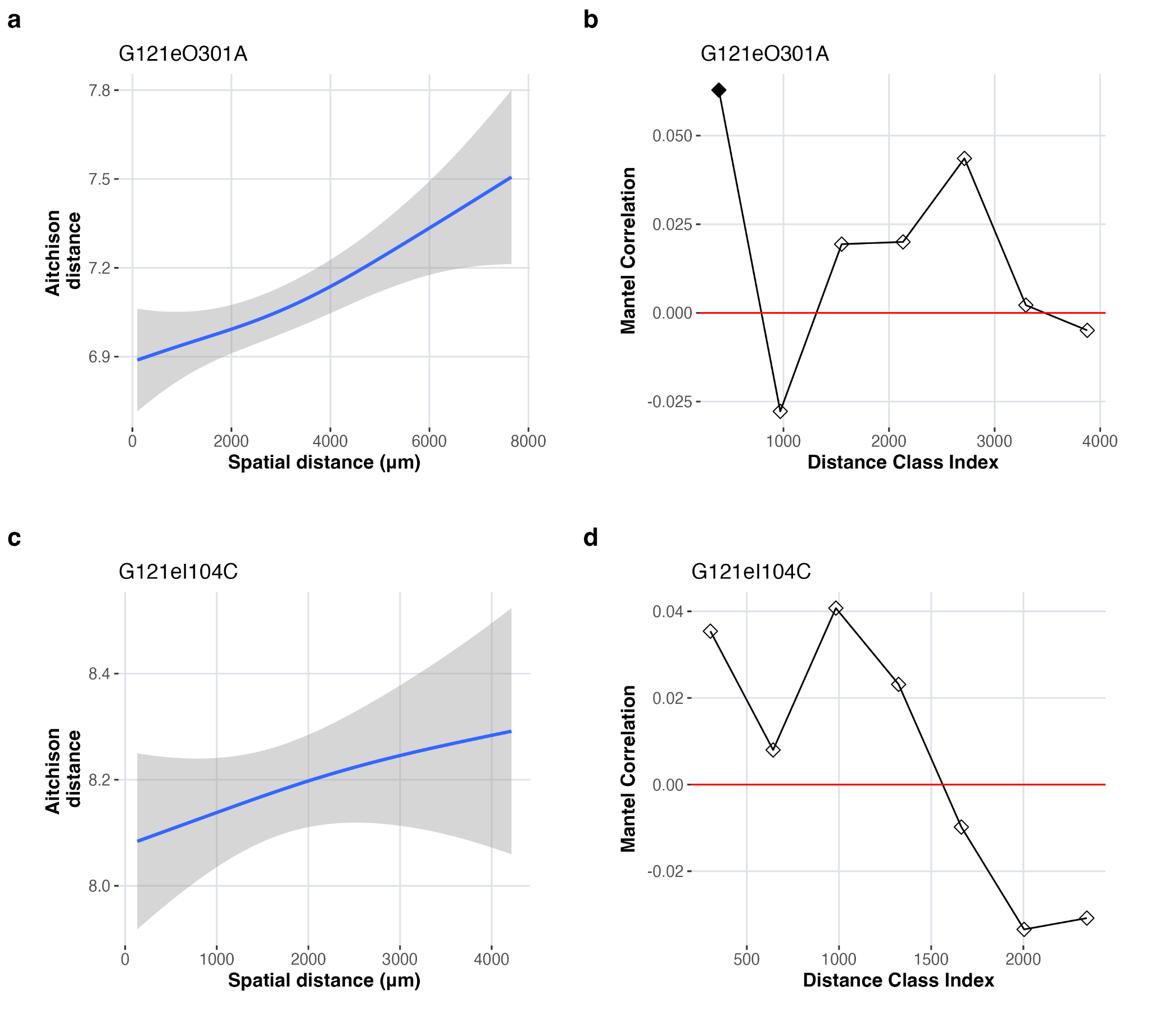

**Fig. S4.2: Distance decay plots of microbial community dissimilarity measured by Aitchison distance and Mantel correlogram analysis. a,** A significant positive correlation (*p*<0.001) between microbial community dissimilarity and spatial distance among microsamples within a colon cryosection (G121eO301A). **b,** Mantel correlogram showing spatial autocorrelation of microbial communities across defined distance classes for the colon cryosection (G121eO301A); filled squares indicate significant autocorrelation. **c,** No significant positive correlation (*p*=0.21) between microbial community dissimilarity and spatial distance was observed among microsamples within a caecum cryosection (G121eI104C). **d,** Mantel correlogram indicating no significant spatial autocorrelation of microbial communities across distance classes for the caecum cryosection (G121eI104C); filled squares indicate significant autocorrelation.

To delve deeper into the spatial structures observed in the colon microbial communities, we applied an extended RLQ analysis based on data from 65 microsamples obtained from a single cryosection. This multivariate method linked the spatial and environmental (microbial species diversity or richness and sequence counts) structure observed in microbial communities with the functional attributes and phylogenetic characteristics of their microbial taxa. Bacterial diversity and sequence counts showed a nearly significant positive spatial autocorrelation at the shortest distances (Moran’s I test,: *p=*0.063 and *p=*0.097 for microbial diversity and sequence counts respectively). Similarly, most studied functional traits showed a phylogenetic signal, with the exception of B04 (Short-Chain Fatty Acids biosynthesis) and B09 (Metallophore biosynthesis), which were excluded from the analysis.

The extended RLQ analysis revealed that there was a spatially structured gradient in the microbial community (Fig. 5b). The gradient varied from microsamples with low microbial diversity and low numbers of sequence counts to those with higher number of species and sequence counts (Fig. S4.3a). Microorganisms exhibited a phylogenetically structured response to this gradient (Fig. S4.3b). Microsamples with lower diversity tended to be dominated by Lactobacillaceae and Lachnospiraceae (Table S4.2.1), characterised by higher predicted capacity to degrade polysaccharides (D02) and sugars (D03) and lower capacity to degrade lipids (D01) (Fig 5c). In contrast, the microsamples with higher diversity were dominated by Oscillospiraceae (Table S4.2.2), associated with a higher predicted capacity to degrade lipids and low capacity to degrade polysaccharides and sugars (Fig 5c).

The data employed in the extended RLQ analysis were derived from colon microsamples subjected to two distinct lysis treatments, one of which produced higher sequence counts and diversity metrics (Supplementary note S3.2). To validate the robustness of our results, all the analyses were repeated independently for each lysis treatment. Results remained consistent, confirming the overall conclusion of the extended RLQ analysis applied to the full set of microsamples.

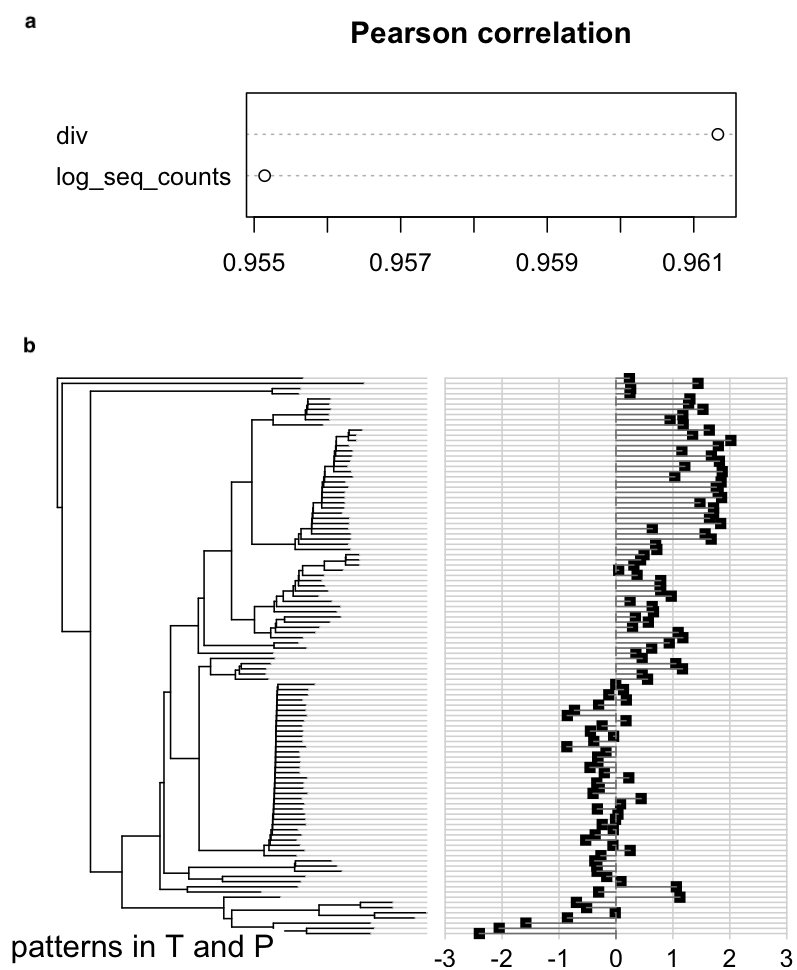

**Fig. S4.3: Results of the extended RLQ analysis for the ﬁrst axis in the colon cryosection (G121eO301A).** **a,** For quantitative environmental variables (div=microbial diversity and log_seq_count=sequencing yield post-quality filtering), Pearson correlations (based on raw data) between the variables and the global coordinates of the sites on the canonical axis are given. **b,** The coordinates of the species, deﬁned as the combination of trait-based coordinates and of phylogeny-based coordinates, expressed on a Cleveland dot plot arranged according to their phylogenetic position.

**S4.2.1: Genomes associated with low-diversity microsamples.** Only top-5 genomes are displayed, while the rest can be found in the code repository.

| **Genome** | **Score** | **Family** | **Species** |
| --- | --- | --- | --- |
| **TG5_28_bin_000004** | -2.1375890 | Lactobacillaceae | *Lactobacillus johnsonii* |
| **D300479_bin_000001** | -1.8214777 | Lactobacillaceae | *Lactobacillus crispatus* |
| **TG5_35_bin_000001** | -0.9768024 | Streptococcaceae | *Streptococcus alactolyticus* |
| **D300452_bin_000016** | -0.9684922 | Lachnospiraceae | *Eisenbergiella* sp904392525 |
| **D300511_bin_000002** | -0.9255171 | Lachnospiraceae | *Eisenbergiella merdigallinarum* |

**S4.2.2: Genomes associated with high-diversity microsamples.** Only top-5 genomes are displayed, while the rest can be found in the code repository.

| **Genome** | **Score** | **Family** | **Species** |
| --- | --- | --- | --- |
| **GPB_bin_000080** | 2.155511 | Oscillospiraceae | *Lawsonibacter asaccharolyticus* |
| **GPB_bin_000197** | 2.123523 | Oscillospiraceae | *Flavonifractor avistercoris* |
| **GPB_bin_000028** | 2.080568 | Oscillospiraceae | *Unclassified Intestinimonas* |
| **GPB_bin_000022** | 2.060220 | Oscillospiraceae | *Dysosmobacter pullicola* |
| **GPB_bin_000015** | 2.046999 | Oscillospiraceae | *Lawsonibacter pullicola* |
